# Lactate promotes longevity through redox-driven lipid remodeling in *Caenorhabditis elegans*

**DOI:** 10.1101/2025.09.25.678523

**Authors:** Arnaud Tauffenberger, Payton J. Netherland, Hubert Fiumelli, Joshua D. Meisel, Frank C. Schroeder, Pierre Magistretti

## Abstract

Lactate has emerged as a key metabolite involved in multiple physiological processes, including memory formation, immune response regulation, and muscle biogenesis. However, its role in aging and cellular protection remains unclear. Here, we show that lactate promotes longevity in *C. elegans* through a mechanism that requires early-life intervention, indicating a hormetic priming effect. This pro-longevity action depends on its metabolic conversion via LDH-1 and NADH, which drives redox-dependent metabolic reprogramming. Multi-omics approaches revealed that lactate induces early-stage metabolic adaptations, with a strong modulation of lipid metabolism, followed by late-life transcriptional remodeling. These shifts are characterized by enhanced stress response pathways and suppression of energy-associated metabolic processes. Our genetic screening identified *sir-2.1*/SIRT1 and *rict- 1/*RICTOR as essential for lactate-mediated lifespan extension. Our findings establish lactate as a pro-longevity metabolite that couples redox signaling with lipid remodeling and nutrient-sensing pathways. This work advances our understanding of lactate’s dual role as a metabolic intermediary and geroprotector signaling molecule, offering insights into therapeutic strategies for age-related metabolic disorders.

## Introduction

Lactate, once considered merely a metabolic byproduct, is now recognized as a pleiotropic signaling molecule with systemic effects across tissues and species. Its physiological impact extends beyond its classical role as an energy metabolite. Lactate is oxidized to pyruvate via lactate dehydrogenase (LDH)^1^. Emerging evidence supports direct mitochondrial and nuclear lactate transport^2^, challenging traditional metabolic paradigms. Lactate modulates cellular metabolism through NAD^+^/NADH ratio shifts associated with its conversion to pyruvate^3^, influencing glycolysis, oxidative phosphorylation^4^, and redox-sensitive pathways such as NMDA receptor-mediated synaptic plasticity^5,6^. Tissue-specific effects are prominent: in muscle, lactate activates pro-growth and stress-resistance pathways (e.g., mitochondrial biogenesis), while in the brain, it enhances synaptic plasticity and memory via BDNF, Arc, and c-fos induction^6–8^. The discovery of GPR81/HCAR1, a lactate-sensing GPCR in adipose tissue and other organs, has further expanded our understanding of lactate as an intercellular signal^9–11^.

Recent advances have revealed lactate’s role as a post-translational modifier. Histone lactylation, mediated enzymatically via p300 using lactyl-CoA and reversed by sirtuins (SIRT1- 3) or HDACs^12^, links lactate flux to gene regulation in physiological and pathological contexts^13–16^. Concurrently, lactate shapes immune responses through mechanisms ranging from metabolic reprogramming to direct receptor interactions^17–19^. Lactate, via its oxidation into pyruvate and TCA-mediated acetyl-CoA production, has also been reported to increase histone acetylation^2^. However, the impact of lactylation or lactate-promoted acetylation in the context of aging and stress resistance has not been investigated. Most evidence supporting a cell-protective role for lactate stems from studies showing that lactate injections improve neuronal survival under hypoxic or glucose-deprived conditions^20–22^. During aging, astrocytes, the primary lactate-producing cells in the brain, undergo senescence, which reduces their metabolic support and thereby promotes cellular stress and accelerated degeneration^23–26^. Maintaining the expression of the lactate transporter MCT1 during aging has been shown to improve peripheral neuron myelination, thereby supporting nerve conduction and sensory function^27^. Lactate supports protection against glutamate toxicity, with the latter dependent on PI3K signaling^28,29^ and NADH-associated metabolism^30^. Lactate also influences ROS balance^31–35^. In this context, we recently showed that this lactate-mediated mechanism regulates stress resilience^36^. Despite these observations, the specific signaling cascades that regulate lactate’s ability to maintain cellular homeostasis remain to be identified. While evidence suggests a role for ROS and lipid droplet exchanges between neurons and astrocytes^37,38^, a comprehensive understanding of how lactate mediates this interaction, and how aging impacts these processes, is lacking. Here, we used transcriptomics and untargeted metabolomics to decipher the role of lactate on longevity using *C. elegans*. We uncover that lactate increases longevity through its metabolic conversion into pyruvate and increased mitochondrial ROS. Redox changes are accompanied by an increase in protein acetylation, regulating the longevity-promoting action of lactate. Our genetic analysis revealed that lactate- promoted longevity relies on skn-1/Nrf-2, *sir-2.1*/SIRT1, and TORC2 complex kinase *rict- 1*/RICTOR as part of a metabolic hub that coordinates redox processes, regulation of protein acetylation, and lipid mobilization.

## Methods

### Strains and maintenance

The strains used in this study were obtained at the *C. elegans* genomic center (CGC). All strains were maintained on Nematode Growth Media (NGM) using standard procedure, at 20 °C unless specified. Mutant lines were outcrossed at least 3 times. The strain list used in this study can be found in Table I.

### Phenotyping

Animals were grown on regular NGM or NGM + supplemented diet. For lifespan and paralysis experiments, 30 L4 stage animals were transferred to new NGM/NGM + diet, in triplicate, without the use of progeny blocking compounds unless specified.

### Lifespan

Survival was scored every 2 days until no animals were alive. Nematodes were considered dead if they failed to react to prodding with a pick. Animals that displayed bagging or desiccation on the side of the dish were censored. **Paralysis:** Mobility was scored every day until day 12 of adulthood. Nematodes were considered paralyzed if they failed to produce forward or backward locomotion after prodding with a worm pick, for a minimum of 30 s. **Aldicarb:** Animals at different aging stages (Day 1, 5, and 9) were placed on media containing 1 mM aldicarb (Sigma-Aldrich #33386). Paralysis was measured every 30 min until the entire population failed to produce locomotion.

### Lipid Staining

Cultures were synchronized from a mix stage culture on regular NGM. 1000 eggs were transferred to NGM or NGM + supplement. For the multi-day experiments, animals were transferred to media with 5-Fluoroacil to block progeny production. For all days evaluated, worms were washed from plates with PBS-T 0.01% and centrifuged 20 s, 250*g* at room temperature (RT). Worms were fixed using 40% (v/v) isopropanol for 3 min under agitation, followed by centrifugation 20s, 250*g* at RT. Animals were stained with a Nile Red (NR) solution (5mg/ml DMSO) diluted in 40% isopropanol (6 μl NR/ml isopropanol 40%). Animals were stained 2h at RT in 1 ml of NR, protected from light. Worms were centrifuged 20 s, 250*g* at RT, resuspended in PBS-T 0.01% and washed for 30 min, protected from light. Worms were centrifuged 20 s, 250*g* at RT, and mounted on 2% agar pad to be imaged on a Leica M205 microscope equipped with a Leica DFC7000 T camera. All image acquisition was performed using Leica LasX software.

### ROS staining

Cultures were synchronized from a mix stage culture, on regular NGM. 1000 eggs were transferred on NGM or NGM + supplement. 72h post-synchronization (adult day 1), worms were washed from plate with M9 + 0.01% Triton X-100 and centrifuged for 20 s, 250*g* at room temperature (RT). Worms were washed an additional time to remove bacteria, followed by centrifugation for 20s, 250*g* at RT. Mitotracker H2X-ROS was diluted in DMSO (10 mM stock solution) and added to a culture of heat-inactivated OP50-1 to a final concentration of 50 μM. Animals were stained 4h at RT, protected from light. After staining, worms were centrifuged for 20 s, 250*g* at RT, washed in M9-T 0.01%. Worms were mounted on 2% agar pad in a 5 mM Levamisole solution and imaged on a Leica M205 microscope equipped with a Leica DFC7000 T camera. All image acquisition was performed using Leica LasX software.

### Fluorescent reporter imaging

Cultures were synchronized from a mix stage culture on regular NGM. 1000 eggs were transferred to NGM or NGM + supplement. For the multi-day experiments, animals were transferred to media with 5-Fluoroacil to block progeny production. Strain imaging was performed on adult animals (D1, 3, and 5) grown on NGM or NGM + supplement (see figure legends). Briefly, animals were washed from plates using 1 mL M9 Buffer, centrifuged at 250*g*, and placed on a 2% agarose pad + 5 mM levamisole. Images were acquired on a Leica M205 using a Leica DFC7000 T camera and LasX software. Fluorescence quantification was measured using Fiji (imageJ) software.

### Immunoblot

#### Cultures

Nematodes were synchronized from a mix stage culture on regular NGM. 2500 eggs were transferred to 6x10 cm NGM or NGM + supplement. For the multi-day experiments, animals were subsequently transferred to media with 5-Fluoroacil to block progeny production at L4 stage. At day 1 or 5 of adulthood, animals were washed with M9 buffer in 15 ml conical tubes and centrifuged for 30 s, 1000 RPM at RT. Pellet washed an additional time using M9 to remove bacteria using same centrifugation parameters. Pellets were transferred into 1.5 ml tubes, centrifuged for 20 s, 250*g* at RT to remove water, and frozen on dry ice.

Protein extraction: Nematode pellets were thawed on ice, in 100 μl RIPA buffer (150 mM NaCl; 1% Triton X-100; 0.5% Na-Deoxycholate; 0.1% SDS; 50 mM Tris pH:8.0) + 1x Halt proteinase- phosphatase inhibitor (ThermoFisher #1861281). Once thawed, pellets were sonicated in a refrigerated water bath. Lysate was subsequently centrifuged for 10 min, 14000*g* at 4 °C, and supernatant transferred to new 1.5 ml tube.

#### Protein quantification and sample preparation

Proteins were measured using a BCA kit following the manufacturer’s instruction. 30 μg of protein were used for immunoblotting.

#### Electrophoresis and transfer

Gels were prepared as 4% acrylamide (stack) and 12% acrylamide (run) gels. Proteins were loaded in 1X sample buffer complemented with ddH_2_O to an equal volume for each sample. Samples were run in 1X SDS running buffer (tris 3g/L; glycine 72 g/l; SDS 1 g/l) at RT together with a BioRad kaleidoscope protein ladder (#1610375). Samples were subsequently transferred on a PVDF membrane (#IPVH00010) using semi-dry transfer (BioRad) following the manufacturer’s instructions.

#### Antibodies staining

Membranes were blocked for 45 min at RT using PBS-T 0.1% + 5% Bovine Serum Albumin (BSA) (REF#). Antibodies were applied overnight at 4 °C in PBS-T 0.1% + 5% BSA. The antibody list and working concentration are as follows: Anti-Lactylysine (PTMBio #1401RM - 1/1000), Anti-Acetyllysine (PTMBio #105RM - 1/1000), Anti-tubulin (REF# - 1/5000). After overnight incubation, membranes were washed three times, 5 min at RT, using PBS-T 0.1%. Secondary antibodies: goat-ms-HRP and goat-Rb-HRP (ThermoFisher #G21234) were applied for 1h at RT as a 1/5000 dilution. Membranes were subsequently washed 3x5 min with PBS-T 0.1% and imaged using Pierce ECL (ThermoFisher #32209) following manufacturer’s instructions.

### RNA extraction

Synchronized cultures (10,000 eggs/condition) were grown on a control or lactate-supplemented diet (10 mM). At adult day 1 or day 5, nematodes were collected using M9, centrifuged 1000 RPM for 1 min at 20 °C. Pellet were washed twice to remove bacteria and frozen on dry ice and stored at -80 °C until processed. Total RNA from WT *C. elegans* was isolated using TRIzol – Phenol/Chloroform method. Worm pellets were thawed on ice with 1 volume of TRIzol, vortexed briefly, and let at room temperature for 5 min. 1/3 volume (TRIzol) of Phenol-Chloroform was added and incubated at RT for 3 min. Samples were centrifuged 13000 RPM at 4 °C for 15 min, and the aqueous phase was transferred into a new clean 1.7 ml tube. RNA was precipitated using 1/2 volume (TRIzol) of isopropanol and incubated 15 min at RT. Samples were then centrifuged 14000 RPM at 4 °C for 15 min. Supernatant was removed and pellets washed using 75% ethanol. Pellets were dried for 10 min at RT and resuspended in 30 μl of ddH_2_O.

### RNA sequencing

Concentration, purity, and integrity of the RNA were assessed with a NanoDrop spectrophotometer (NanoDrop 2000, ThermoFisher Scientific), and a 2100 Bioanalyzer (Agilent).

Total RNA with an RNA Integrity Number above 9.5 was used to construct libraries using the TruSeq Stranded mRNA Sample Kit (Illumina) following the protocol’s instructions. Briefly, mRNA was enriched using oligo dT-attached magnetic beads, fragmented, and converted into cDNA. Fragments of cDNA went through an end repair process, 3’ ends were adenylated, universal bar-coded adapters were ligated, and cDNA fragments were amplified by PCR to yield the final libraries. The sequencing libraries were evaluated using a 2100 Bioanalyzer (Agilent). Paired-end read (2 x 150 bp) multiplex sequencing from pooled libraries was performed on an Illumina HiSeq 4000 machine by Macrogen Inc. (Seoul, South Korea). An average of 60-70 million reads was obtained for each sample. Sequencing data have been deposited in the NCBI SRA database under the project accession number PRJNA1306788.

Raw read quality was evaluated with the FastQC tool (https://www.bioinformatics.babraham.ac.uk/projects/fastqc/). Low-quality reads were filtered out and adapter sequences trimmed using Trimmomatic version 0.36^39^ with the following parameters: ILLUMINACLIP/TruSeq3-PE-2.fa:2:30:10, LEADING:3, TRAILING:3, SLIDINGWINDOW:4:15, MINLEN:36. Reads from each sample replicate were mapped to the *Caenorhabditis elegan*s reference genome (WBCel235) using STAR version 2.6.0a^40^ with default parameters except for outFilterMultimapNmax set to 1. Mapped reads for protein-expressing genes were summarized with the featureCounts program (Subread package, version 1.5.2,^41^), and the differential expression analysis was performed with the Bioconductor package DESeq2^42^ in the R programming environment. To minimize background noise and to focus on more significant genes in terms of biological impact, we removed genes with very low expression levels, excluding genes that failed to total an average count above 10 in any conditions. Differentially expressed genes (DEG) were considered in pairwise comparisons with a threshold including a fold change expression ≥ 1.5 and q-value (or False Discovery Rate, FDR) < 0.05. To obtain a functional representation of the lists of DEG, we performed gene ontology (GO) and pathways enrichment analyses using the online database Wormcat^43^.

### Liquid cultures

Mixed stage cultures were grown in S-complete^44^ with added concentrated OP50-1 for approximately 7-8 days. Cultures were checked daily to ensure enough food, and we kept nematode concentration to a maximum of 5 animals/μl. Cultures were synchronized using bleaching solution (20% bleach, 1N NaOH). Then 80,000 synchronized eggs were added to 125 ml Erlenmeyer flasks containing 30 ml of S-complete + concentrated OP50-1 and incubated at 20 °C with shaking at 160 rpm.

On day 1 of adulthood, cultures were centrifuged (1,000 rpm, RT, 30 s) in 15 ml conical tubes. Supernatants were collected in separate tubes and frozen on dry ice. Pellets were washed with M9 and split in half (40,000 animals). One half was frozen on dry ice, and the other was transferred to 30 ml S-complete + OP50-1 and incubated at 20 °C with shaking at 160 rpm. For all following days (2 to 5), cultures were centrifuged (1,000 rpm, 30 sec at RT), and the supernatant was collected in a separate tube and frozen on dry ice. Pellets were washed with M9 to remove larvae using a settling method. Once pellets were cleared from larvae, animals were transferred into fresh 30 ml S-complete + OP50-1 and incubated at 20 °C while shaking at 160 rpm for 24 h. On day 5, pellets were frozen on dry ice. OP50 only cultures were grown in parallel to nematodes. Once collected and frozen, all samples were kept at -20 °C until processed.

### Sample preparation

Frozen samples were lyophilized using a Virtis sentry 2.0 lyophilizer for 24 h to 48 h. Pellets were then sonicated in 2 ml of methanol using Qsonica Q700 Ultrasonic Processor with a water bath cup horn adaptor (Qsonica 431C2). After sonication, 3 ml of methanol was added, and samples were incubated at RT on an orbital shaker. Supernatant samples were resuspended in 5 ml of methanol and incubated overnight at RT on an orbital shaker. Methanol suspension was centrifuged (2,500*g*, 10 °C, 5 min) to remove any precipitate, and the supernatant was carefully transferred to 8 ml glass vials. Samples were dried via an SC250EXP SpeedVac (ThermoFisher Scientific) vacuum concentrator. Dried materials were resuspended in 1 ml of methanol, vortexed for 30 s, and sonicated for 10 min at RT. Suspensions were transferred into 1.7-ml Eppendorf tubes and centrifuged at 14,000*g* for 5 min at 10 °C. Supernatants were transferred in HPLC vials. Samples were then dried a second time using the SpeedVac, resuspended in either 200 μl (exo-metabolome) or 100 μl (endo- metabolome). Samples were then vortexed for 30 s, sonicated for 10 min at RT, and transferred to 1.7 ml tubes. After a 5 min, 14,000*g* centrifugation, supernatants were transferred back to HPLC vials with an insert and stored at -20 °C until analyzed by mass spectrometry.

### Mass spectrometry

Liquid chromatography was performed on a Vanquish HPLC system controlled by Chromeleon Software (ThermoFisher Scientific) and coupled to an Orbitrap Q Exactive High Field mass spectrometer controlled by Xcalibur software (ThermoFisher Scientific). Methanolic extracts prepared as described above were separated on a Thermo Hypersil Gold C18 column (150 mm × 2.1 mm, particle size 1.9 μM; 25002-152130) maintained at 40°C with a flow rate of 0.5 mL/min. Solvent A: 0.1% formic acid (Fisher Chemical Optima LC/MS grade; A11750) in water (Fisher Chemical Optima LC/MS grade; W6-4); solvent B: 0.1% formic acid in acetonitrile (Fisher Chemical Optima LC/MS grade; A955-4). A/B gradient started at 1% B for 3 min after injection and increased linearly to 98% B at 20 min, followed by 5 min at 98% B, then back to 1% B over .1 min and finally held at 1% B for the remaining 2.9 min to re-equilibrate the column (28 min total method time). Mass spectrometer parameters: spray voltage, −3.0 kV/+3.5 kV; capillary temperature 380 °C; probe heater temperature 400 °C; sheath, auxiliary, and sweep gas, 60, 20, and 2 AU, respectively; S-Lens RF level, 50; resolution, 120,000 at *m/z* 200; AGC target, 3E6. Each sample was analyzed in negative (ESI−) and positive (ESI+) electrospray ionization modes with *m/z* range 100–1200.

Parameters for MS/MS(dd-MS2): MS1 resolution, 60,000; AGC Target, 1E6. MS2 resolution, 30,000; AGC Target, 2E5. Maximum injection time, 60 msec; Isolation window, 1.0 *m/z*; stepped normalized collision energy (NCE) 10, 30; dynamic exclusion, 5 s; top 8 masses selected for MS/MS per scan. Inclusion lists with 20 sec windows were generated in Metaboseek for targeted MS/MS^45^ .

Hydrophilic Interaction Liquid Chromatography (HILIC) was performed on a Waters XBridge Amide column (150 mm × 2.1 mm, particle size of 3.5 μm) maintained at 40 °C (solvent A, 0.1% ammonium formate in 90% acetonitrile–10% water; solvent B, 0.1% ammonium formate in 30% acetonitrile–70% water). A flow rate of 0.5 ml min^−1^ was used, and the A–B gradient was as follows: being isocratic at 1% B for 3 min, linearly increasing to 25% B at 20 min, linearly increasing to 100% B at 22 min, keeping at 100% B for 4.9 min, shifting back to 1% B in 0.1 min and holding at 1% B until 30 min (mass spectrometer parameters: spray voltage, +3.5 kV; capillary temperature, 380 °C; sheath gas, 60 psi; auxiliary gas, 20 psi; spare gas, 1 psi; probe heater temperature, 300 °C; S-lens RF level, 50; resolution, 240,000 at an *m*/*z* ratio of 200; AGC target, 3 × 10^6^). The instrument was calibrated with positive ion calibration solution (Thermo Fisher Scientific) Reversed-phase post-column ion-pairing chromatography (PCI) was performed using the same system as described; extracts were separated on a Thermo Scientific Hypersil Gold column (150 mm × 2.1 mm, particle size 1.9 μm, part no. 25002-152130) or on a Kinetex Evo C18 (150 mm × 2.1 mm, particle size 1.7 μm, part no. 00F-4726-AN) maintained at 40 °C with a flow rate of 0.5 mL/min. Solvent A: 0.1% ammonium acetate in water; solvent B: acetonitrile. A/B gradient started at 5% B for 3 min after injection and increased linearly to 98% B at 20 min, followed by 5 min at 98% B, then back to 5% B over 0.1 min and finally held at 5% B for an additional 2.9 min. A second pump (Dionex 3000) controlling a solution of 800 mM ammonia in methanol was run at a constant flow rate of 0.015 mL/min for the duration of the method and mixed via micro-splitter valve (Idex #P-460S) with the eluate line from the column. HPLC-HRMS RAW data were converted to mzXML file format using MSConvert (v3.0, ProteoWizard) and were analyzed using Metaboseek software (v0.9.9.0) with the following settings: 5 ppm, 320 peakwidth, 3 snthresh, 3100 prefilter, FALSE fitgauss, 1 integrate, true firstBaselineCheck, 0 noise, wMean mzCenterFun, −0.005 mzdiff. Default settings for XCMS feature grouping: 0.2 minfrac, 2 bw, 0.002 mzwid, 500 max, 1 minsamp, FALSE usegroups. Metaboseek peak filling used the following settings: 5 ppm_m, 5 rtw, TRUE rtrange, FALSE areaMode. Quantification was performed with Metaboseek software or via integration using Xcalibur QualBrowser v4.1.31.9 (Thermo Fisher Scientific) using a 5-ppm window around the m/z of interest. ‘Basic analysis’ and ‘Fast peak shapes’ were selected in the subsequent Metaboseek analysis.

### Statistics and software

Fiji ImageJ was used for image quantification. GraphPad Prism 10 was used for statistical analyses. Statistical details of experiments can be found in the figures’ legends. The log-rank (Mantel–Cox) method was used to compare survival curves. Mann–Whitney test with Benjamini–Hochberg test for multiple hypothesis correction was used to compare transcript levels. The paired t-test was used to compare the expression of antibody signals in immunofluorescence samples and western blots. For RNA-seq analysis, DEGs were identified using the Wald test. For all experiments, P values <0.05 were considered significant. No statistical method was used to predetermine the sample size. No data were excluded from the analyses. The experiments were not performed blinded, but worms were arbitrarily distributed for all experiments.

### Data availability

All raw and processed sequencing data for RNA-seq libraries can be found under NCBI Gene SRA database under PRJNA1306788.

The HPLC-HRMS data generated during this study have been deposited in the MassIVE database under accession code MSV000099134.

Source data are provided with this article. Other original data will be available from the Lead Contact upon request.

## Results

### Lactate promotes longevity in a timing -specific manner

We previously demonstrated that high concentrations of L-lactate (100 mM, hereafter referred to as lactate) increase resistance to oxidative stress induced by juglone but paradoxically shorten lifespan in C. *elegans*^36^. To determine whether lower concentrations of lactate could promote longevity, we tested various concentrations. We found that lactate significantly extended lifespan at 5 and 10 mM **(Fig. 1A)**. In contrast, higher concentrations failed to improve longevity, consistent with our previous studies **(Fig. S1A)**. Because live bacteria can metabolize lactate, we tested whether lactate’s effects depended on bacterial metabolism. Lifespan extension was still observed in animals fed heat-killed *E. coli* **(Fig. S1B)**, indicating that the effect is independent of bacterial metabolism. Lactate exists as two enantiomers, L- and D-lactate. To assess whether the stereochemistry of lactate influences its effects on longevity, we supplemented animals with D-lactate but observed no lifespan extension **(Fig. S1C)**.

**Figure 1:**
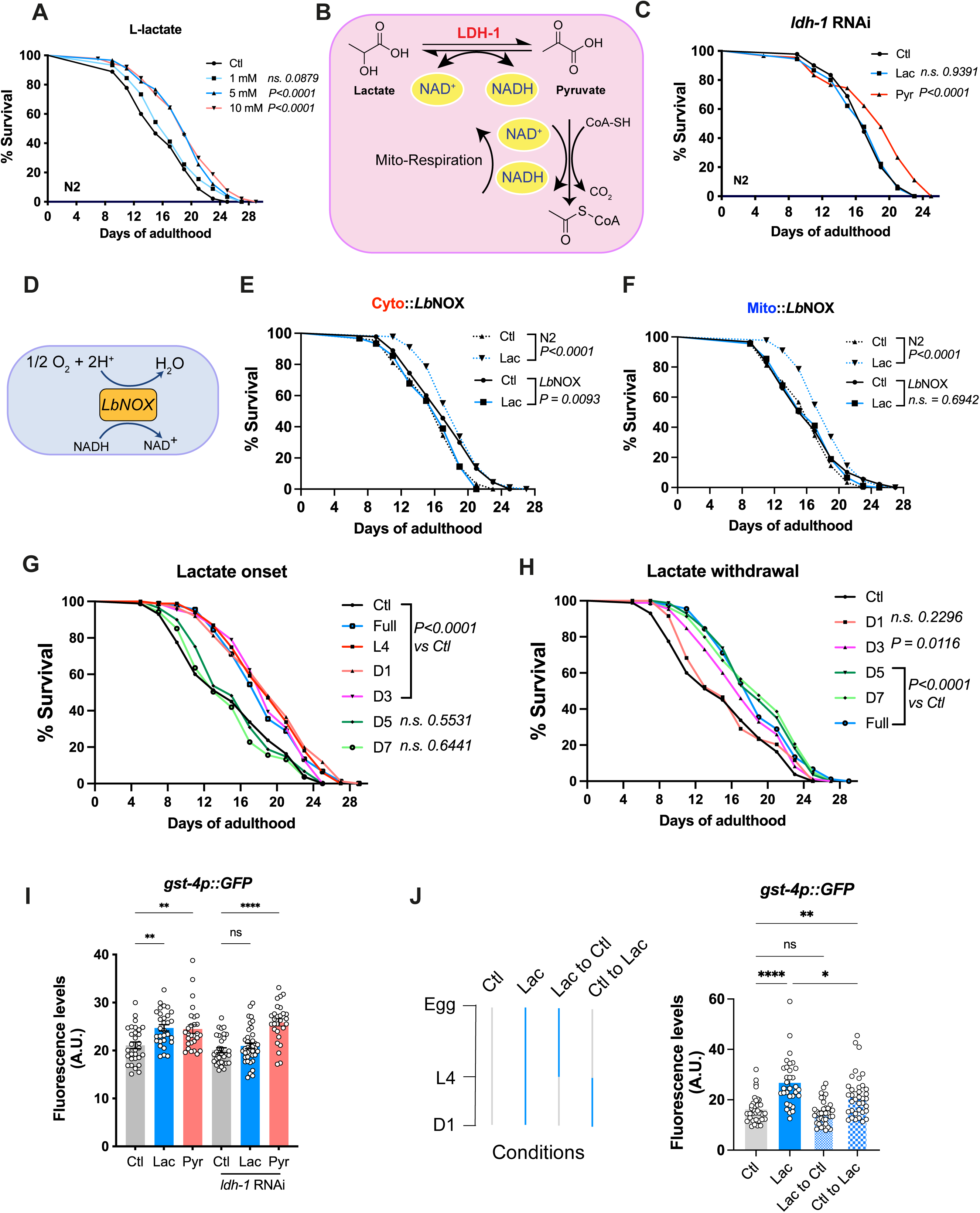
**Lactate catabolism regulates longevity**. **A)** Lifespan curves of N2 animals under control (black line) and lactate-supplemented (colored lines) diets. **B)** Schematic representation of metabolic pathways associated with the lactate dehydrogenase reaction and subsequent pyruvate oxidation to acetyl-CoA. **C)** Lifespan curves of N2 animals treated with *ldh-1* RNAi, under control (black) or lactate- (blue) and pyruvate-supplemented (red) diets. 90 animals were scored per condition. Treatments were applied from the embryonic stage until death. **D)** Schematic representation of the *Lb*NOX-dependent mechanism. **E-F)** Lifespan curves for animals expressing *Lb*NOX in the cytoplasm **(E)** or in the mitochondria **(F)**. Animals were grown on control (black) or lactate-enriched diet (blue). **G-H)** Lifespan curves of N2 animals under control diet (black) or different temporal lactate supplementation. Full treatment refers to continuous lactate exposure from the embryonic stage until death. Other conditions refer to stages when lactate supplementation was initiated and continued until the death of the animals tested. **I)** Quantification *gst-4*p::GFP fluorescence in day 1 adult animals treated with *ldh-1* RNAi. Animals were grown under control (grey), lactate- (blue) or pyruvate-supplemented (red) diet. **J)** Experimental design for mid-developmental stages. For the Lac to Ctl condition, L4 animals were transferred from lactate-supplemented to control diet. For the Ctl to Lac condition, L4 animals were transferred from control to lactate-supplemented diet. *gst-4*p::GFP expression was quantified in day 1 adult animals, and each dot represents one animal. Statistics for lifespan curves were done using log-rank (Mantel–Cox) method. For each condition, 90 animals were analyzed (in triplicate), and each experiment was repeated at least twice. P values for fluorescence measurements were calculated by unpaired, two-sided t-test with Welch correction. *P<0.05; **P<0.01; ****P<0.0001. ns: Not significant. Each fluorescence experiments were performed at least 3 times, and between 25-40 animals were measured per condition.

To better understand the metabolic basis of lactate-mediated longevity, we investigated the oxidation of lactate to pyruvate by lactate dehydrogenase (LDH), a reaction that reduces NAD+ to NADH. **(Fig. 1B)**. Pyruvate supplementation at the same concentrations as lactate also extended lifespan in N2 animals to a similar extent **(Fig. S1D–E)**. To assess whether lactate-driven longevity depends on the conversion of lactate to pyruvate, we knocked down *ldh-1* by RNAi. KD of *ldh-1* abrogated the lifespan extension conferred by lactate, but not pyruvate **(Fig. 1C)**, indicating that LDH-mediated conversion of lactate into pyruvate is essential for lactate’s activity.

Pyruvate is further metabolized into acetyl-CoA and CO_2_, generating a second equivalent of NADH **(Fig. 1B)**. To test whether lactate- or pyruvate-derived NADH generation could underlie the observed longevity effects, we used strains expressing the *Lactobacillus brevis* NADH oxidase (*Lb*NOX), an enzyme that oxidizes NADH to NAD⁺^46,47^, bypassing the canonical regeneration of NAD^+^ via mitochondrial respiration **(Fig. 1D)**. Expression of *Lb*NOX in either the cytoplasm or mitochondria prevented lifespan extension by lactate **(Fig. 1E–F, Fig. S1H)**, suggesting that either high NADH levels or increased NAD^+^ regeneration via mitochondrial respiration are required for lactate’s pro-longevity effects.

Next, we examined the effect of lactate on *C. elegans* aging in greater detail. While lactate supplementation improved mid-life survival, its impact on maximum lifespan was less pronounced. To further investigate the timing of lactate’s effects on longevity, we exposed wild-type animals to lactate at different life stages. Lactate extended lifespan similarly regardless of whether treatment was started at embryonic stages, at late larval stages (L4), or up until day 3 of adulthood, but failed to increase lifespan when treatment began later in life **(Fig. 1G, S1F)**. Conversely, interrupting lactate supplementation during early adulthood (day 3) largely abolished its pro-longevity effects, whereas withdrawal at later stages did not impair lifespan benefits **(Fig. 1H)**. These findings underscore the significance of timing in metabolic interventions designed to delay aging.

Having previously demonstrated that lactate increases the expression of glutathione S-transferase 4 (*gst-4*), a well-established downstream target of the SKN-1/Nrf2 oxidative stress response pathway^48^, we first validated that treatment with 10 mM lactate robustly activates GST-4 expression in *C. elegans*. Using a GFP-tagged *gst-4* transcriptional reporter, which showed a marked increase in GFP fluorescence following lactate exposure^49^ **(Fig. S1I)**. Similar to its lifespan effects, this induction required metabolic conversion of lactate via LDH, as RNAi knockdown of *ldh-1* abolished GST-4 activation by lactate but not by pyruvate **(Fig. 1I)**. To further investigate the temporal dynamics of this response, we monitored GST-4 expression over time. Lactate supplementation during the L4-to-adult day 1 transition significantly increased GST-4 levels within 24 hours **(Fig. 1J)**. This induction was reversible: GFP signal returned to baseline within 24 hours after removing lactate from the diet **(Fig. 1J)**. These results indicate that lactate triggers a rapid and transient activation of stress response pathways, which may underlie its longevity-promoting effects.

In addition to promoting longevity, lactate demonstrated neuroprotective properties. In wild-type animals, lactate did not acutely affect neuromuscular junction (NMJ) function in young adults (day 1), as measured by aldicarb-induced paralysis assays. However, lactate significantly delayed age-related NMJ decline in older animals, improving resistance to paralysis at day 5 and day 9 adulthood **(Figs. S2A–C)**. To further explore lactate’s neuroprotective effects, we tested its impact in two distinct models of neurodegeneration. In *unc-47(e307)* mutants, which lack the vesicular GABA transporter and exhibit progressive motor deficits, lactate treatment delayed the onset and slowed the progression of motor dysfunction. Similarly, in a pan-neuronal polyglutamine disease model expressing 67 glutamine repeats, lactate reduced the severity and delayed the onset of motility impairments **(Figs. S2D–G)**.

Importantly, unlike many pro-longevity interventions, lactate supplementation did not impair reproductive fitness. Brood size and egg-laying timing remained unchanged in treated animals compared to controls **(Fig. S2G)**. Together, these findings highlight lactate’s dual role as a regulator of aging and neuronal health. This lack of reproductive trade-offs sets lactate apart from other longevity-promoting interventions and suggests a unique capacity to balance acute stress signaling with sustained physiological benefits.

### Lactate regulates longevity via redox changes

The observation that lactate promoted healthspan in a narrow temporal window and had very limited late-life benefits **(Fig. S2H)** led us to hypothesize that sustained stress signaling might be the cause of this phenomenon. Since both deletion and activation of *skn-1*/Nrf-2 can reduce lifespan^50,51^, and lactate induces GST-4 expression, we examined how GST-4 level varies over time. We observed that indeed, lactate supplementation maintains high GST-4 expression over 5 days of adulthood **(Fig. 2A)**. RNAi-mediated knockdown of *skn-1* abolished both lactate-mediated longevity **(Fig. 2B)** and GST-4 induction **(Fig. 2C)**, confirming the requirement of SKN-1. To assess whether these responses are redox-dependent, we treated animals with the antioxidant N-acetylcysteine (NAC). The use of 5 mM NAC completely prevented lactate-induced GST-4 activation **(Fig. 2D)**. It blocked the pro-longevity effect of lactate, while also strongly reducing the longevity in N2 animals **(Fig. 2E)**, as previously reported^52^. Finally, we measured ROS levels in N2 animals using Mitotracker and found that lactate induced a modest increase in mitochondrial ROS **(Fig. 2F)**, consistent with our prior observation^36^. These results support a model in which lactate extends lifespan by sustaining redox-sensitive stress responses via SKN-1 activation.

**Figure 2:**
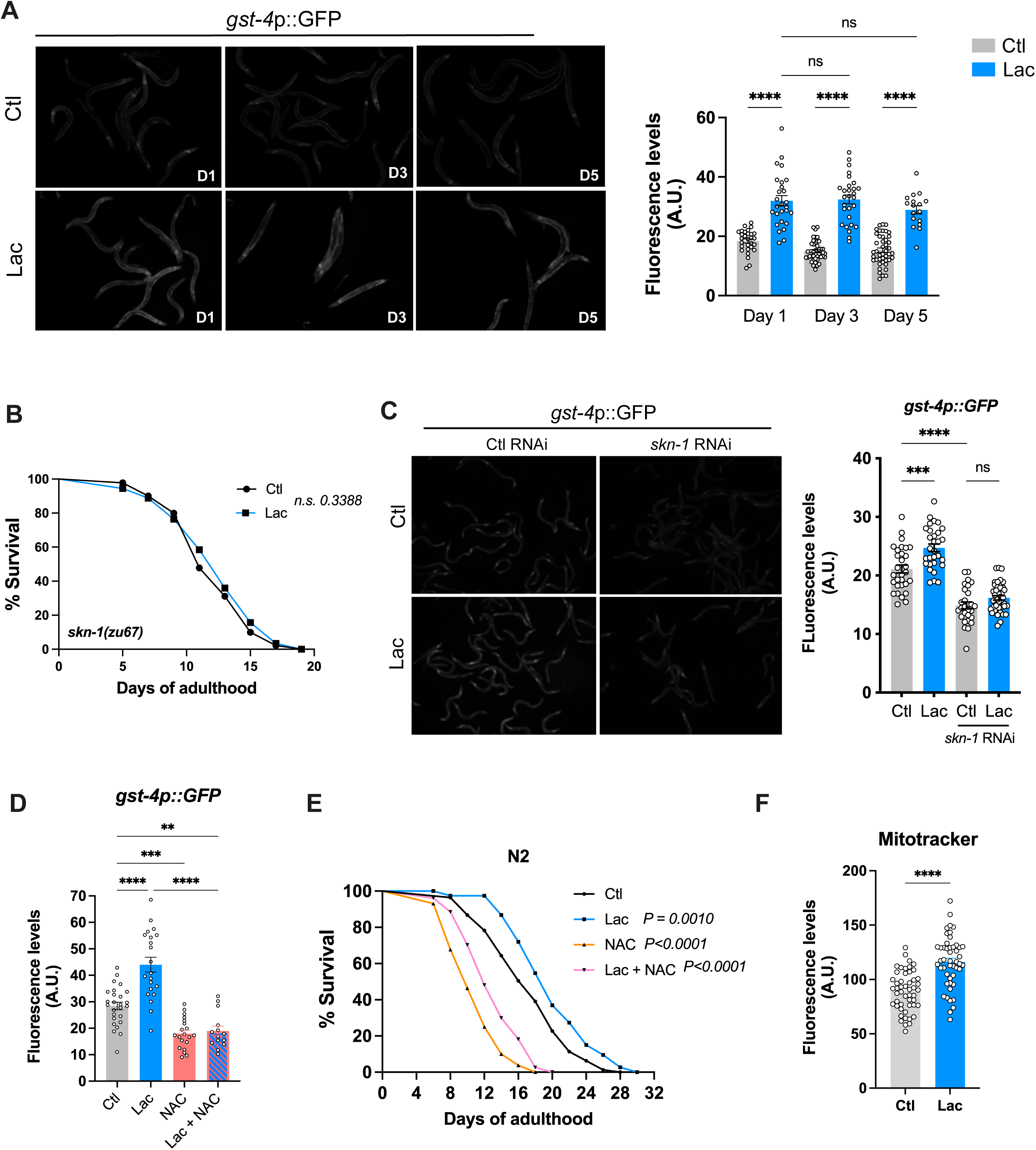
Lactate extends longevity through redox changes A) Representative images of *gst-4*p::GFP reporter animals at day 1 (D1), day 3 (D3), and day 5 (D5) of adulthood under control or lactate-supplemented (10 mM) diets. Quantification of *gst-4*p::GFP fluorescence intensity from animals shown in the images. Data represent pooled measurements from three independent experiments, each including between 20 and 40 animals per condition. B) Lifespan curve for *skn-1(zu67)* mutant animals maintained on control (circles, black line) or lactate-supplemented (squares, blue line, 10 mM) diets. 90 animals were scored per condition. C) Images of *gst-4*p::GFP reporter in day 1 animals. Animals were grown on L4440 RNAi vector and transferred on *skn-1* RNAi at L4 stage. Quantification of *gst-4*p::GFP expression in animals treated with *skn-1* RNAi. D) Quantification of *gst-4*p::GFP expression in animals treated under control, 10 mM lactate (lac), 5 mM N-acetylcysteine (NAC) or lactate + NAC (Lac + NAC) diets. Between 20 and 40 animals were scored per condition. Experiment was repeated three times. E) Lifespan curves of N2 animals treated under control, 10 mM lactate (lac), 5 mM N- acetylcysteine (NAC) or lactate + NAC (Lac + NAC) diets. F) Quantification of mitotracker-H2Xros staining of day 1 N2 animals. Samples were grown on control of lactate-supplemented diet. Statistics for lifespan curves were done using log-rank (Mantel–Cox) method. For each condition, 90 animals were analyzed (in triplicate), and each experiment was repeated at least twice. P values for fluorescence measure were calculated by unpaired, two-sided t-test with Welch correction. *P<0.05; **P<0.01; ****P<0.0001. ns: Not significant. Each fluorescence experiment was performed at least 3 times, and between 25-40 animals were measured per condition.

### Lactate induces late transcriptomic changes

To investigate the transcriptional changes underlying lactate’s pro-longevity effects, we performed bulk RNA sequencing on wild-type *C. elegans* treated with 10 mM lactate or maintained on control diet, collected at early adulthood (day 1) and mid-life (day 5) **(Fig. 3A)**. Differential gene expression analysis was conducted using a fold change threshold of ≥ 1.5 and an adjusted p-value < 0.05 to define significance. At day 1, lactate induced surprisingly few transcriptional changes, with only 3 genes meeting these criteria **(Fig. S3A)**. In contrast, by day 5, lactate elicited a substantial transcriptional response. To assess the interaction between aging and lactate at the transcriptome level, we compared day 5 control and day 5 lactate-treated animals with day 1 controls **(Fig. 3B)**. Aging alone (day 5 vs. day 1 control) led to the upregulation of 1,411 transcripts and downregulation of 620 transcripts **(Fig. 3C)**. Under lactate supplementation, the transcriptional response was broader, with 1,653 transcripts upregulated and 2,090 downregulated relative to day 1 control animals **(Fig. 3C)**. To identify genes specifically regulated by lactate during aging, we compared DEGs from the day 5 lactate-treated animals (*D5 Lac vs. D1 Ctl*) with those from the day 5 control animals *(D5 Ctl vs. D1 Ctl*). This revealed 889 transcripts uniquely upregulated and 1,550 uniquely downregulated in lactate-treated animals **(Fig. 3C**, **Table I and II)**, defining a lactate-specific aging signature.

**Figure 3:**
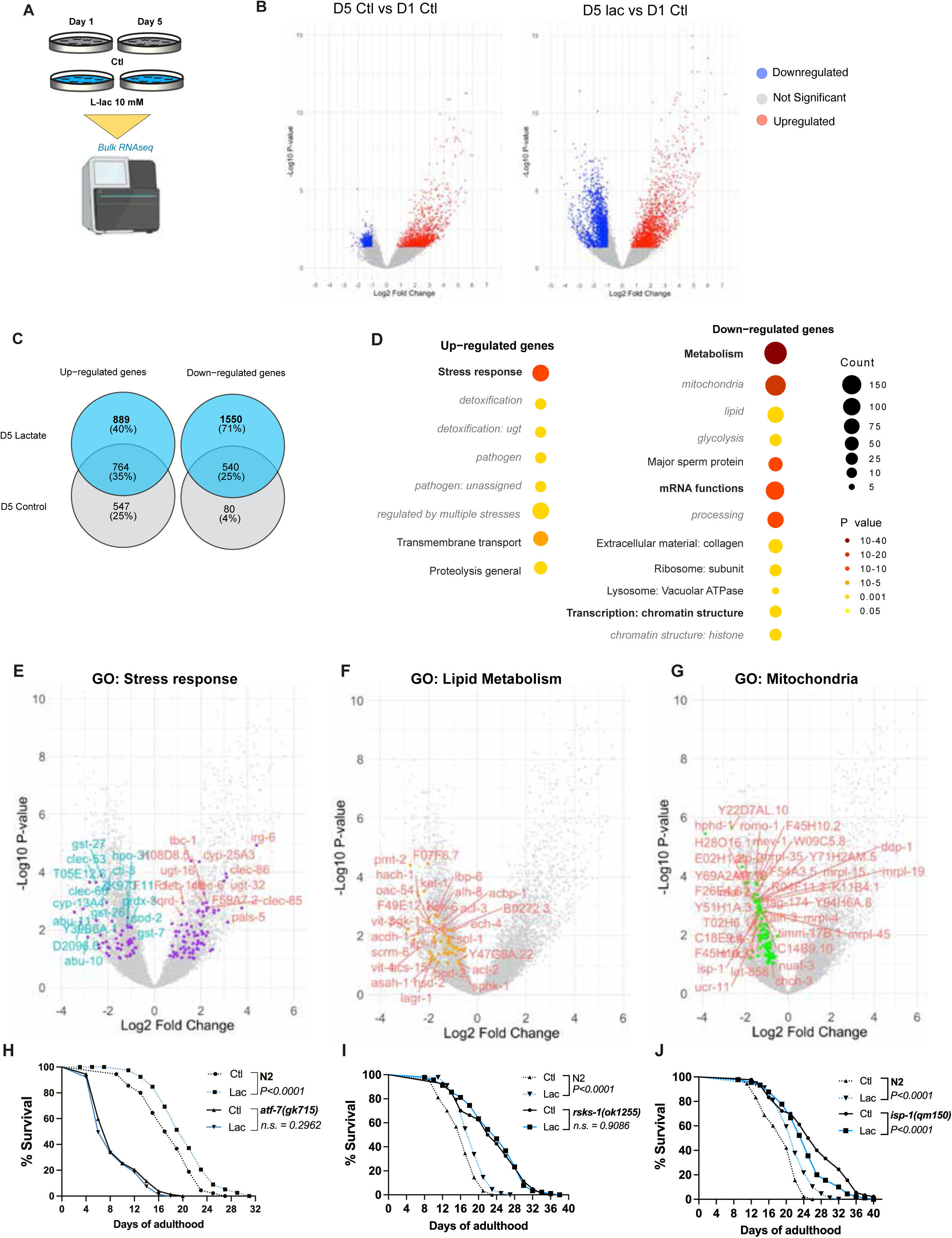
Lactate induces late transcriptomic changes B) Schematic representation of the experimental design used for RNA sequencing. C) Volcano plots representing differentially expressed genes between day 1 controls (D1 Ctl) and day 5 controls (D5 Ctl) (left) or day 5 lactate-supplemented animals (D5 Lac) (right). Significantly up-regulated (red) and down-regulated (blue) genes were identified using thresholds of fold-change ≥ 1.5 and false discovery rate (FDR) <0.05. D) Venn diagram showing numbers of differentially expressed genes (DEG) in aging (D1 Ctl vs D5 Ctl, grey) and in lactate-treated aging (D1 Ctl vs D5 Lac, blue). E) Gene Ontology (GO) analysis (Biological Process) for DEGs between day 5 lactate-treated animals and day 5 controls, performed using *Wormcat*^43^ . Enriched categories, with p adjusted value < 0.05, are shown. **E-J)** Lifespan curves for *atf-7(gk715), rsks-1(ok1255)*, and *isp-1(qm150)* mutants under control (black) or lactate-supplemented (blue) diets. Differential gene expression (DEG) between D5 lactate and D1 control conditions was assessed using the unpaired, nonparametric Mann–Whitney test followed by Benjamini– Hochberg correction for multiple comparisons. Cutoff for significance was fold change ≥1.5 and P-adjusted value of ≤0.05. A list of transcripts specifically regulated by lactate was generated by comparing DEG D5 Lactate vs D1 control vs D5 control vs D1 control. Gene ontology analysis for differentially regulated genes between day 5 ctl and day 5 lac animals was performed using *Wormcat*^43^ . Enriched categories with an adjusted P-value ≤0.05 were considered significant. Lifespan curves were compared using the log-rank (Mantel–Cox) method. For each condition, 90 animals were analyzed (in triplicate). Each experiment was repeated at least twice.

Next, we performed gene ontology (GO) enrichment analysis on these lactate-specific DEGs to identify biological processes associated with the transcriptional response to lactate during aging. Genes associated with specific stress response pathways were significantly overrepresented in our analysis **(Fig. 3D)**. This included both up- and downregulated genes **(Fig 3E)**. Transcripts involved in detoxification **(Fig. S3B)** and pathogen response **(Fig. S3C)** were predominantly upregulated, consistent with activation of defense mechanisms. In contrast, downregulated transcripts were primarily associated with core metabolic processes, including mitochondrial function, lipid metabolism, glycolysis, and, more broadly, mRNA processing **(Fig. 3D)**. Cellular component analysis further revealed a marked enrichment of genes involved in lipid metabolism **(Fig. 3F, Fig. S3E),** mitochondrial respiration **(Fig. 3G),** and glycolysis **(Fig. S3D)**. While these pathways were already modestly downregulated during aging (day 5) in control animals, lactate supplementation induced a significantly stronger suppression **(Figs. S3E, F)**, pointing to a coordinated metabolic reprogramming. These transcriptional signatures suggest that lactate promotes a metabolic shift away from energy-intensive processes, while enhancing stress-resilience pathways. Consistent with an increased stress response, cellular components involved in transcription and mRNA processing were downregulated in day 5 adults treated with lactate **(Fig.S3G-I)**. Overall, these findings support a model in which lactate induces stress response programs while driving mid-adulthood remodeling of metabolic, transcriptional, and translational processes.

We next validated whether the transcriptional changes induced by lactate underlie its capacity to increase longevity. Responses to pathogens involve a plethora of components. However, based on our early findings that lactate-driven longevity is dependent on *skn-1*, we turned to two key elements of the innate immunity pathway, *pmk-1*/p38-MAPK, and the downstream transcription factor *atf-7*/ATF7, both required for the transcriptional activation of defense genes^53^. Loss-of-function mutation for *atf-7(gk715)* abolished the lifespan extension conferred by lactate supplementation, while lactate-driven longevity was not impaired in *pmk-1(km25)* mutants **(Fig. 3H, Fig. S4A)**. We further examined additional stress-responsive pathways and found that disrupting the unfolded protein response (UPR) in either the mitochondria (*atfs- 1*/ATFS1) or the endoplasmic reticulum (ER) (*atf-6*/ATF6 and *xbp-1*/XBP1) also blunted lactate’s pro-longevity effects **(Fig. S4B-D)**. These findings establish that intact stress defense mechanisms, including p38 MAPK-ATF-7 signaling and organellar UPR, are essential mediators of lactate-induced longevity.

We demonstrated that lactate triggers a redox signal during early adulthood, accompanied by a mild increase in mitochondrial ROS **(Fig. 1 and 2)**, a pattern also observed in our previous work^35^. This early redox signaling may prime the strong transcriptional response observed later in life (mid-adulthood) that underlies increased longevity. Building on this model and considering the changes in translation and mRNA processing revealed by our transcriptomic analysis, we next investigated potential roles of the mechanistic target of rapamycin (mTOR), a key regulator of stress responses and cellular metabolism. Previous studies have implicated mTOR in lactate-driven adaptations in muscle^54^ and innate immunity^55^. Because *let-363* (mTOR) mutants are embryonic lethal, we tested two downstream effectors: *rsks- 1(ok1255)*/S6K *and ife-2(ok306)/*eIF4E, both known to modulate longevity. In these mutants, lactate failed to further extend lifespan **(Fig. 3I and S4E)**, suggesting that inhibition of mTOR signaling contributes to the pro-longevity effects of lactate. We further tested for potential roles of mitochondrial respiration, a process linked to longevity via ROS modulation. Lactate supplementation failed to extend lifespan in *isp-1(qm150)* animals, carrying a mutation in mitochondrial complex III **(Fig. 3J)**, supporting the idea that intact mitochondrial function is required for lactate’s pro-longevity activity, and suggesting that lactate may exert its effects in part through modulation of mitochondrial metabolism^54^.

Finally, we evaluated several canonical aging and metabolic pathways^55–58^. Mutants for *daf- 16*/FOXO3A, *hsf-1*/HSF1, insulin/IGF signaling (*age-1*/PDK1, *akt-1*/AKT1), *hlh-30*/TFEB, and *aak-2*/AMPK, all retained sensitivity to lactate-mediated lifespan extension **(Figs. S4F–K)**, indicating these pathways are dispensable for its effects. Collectively, our data show that lactate induces late-life transcriptional reprogramming which shifts the metabolic signature toward reduced lipid mobilization while activating cellular defense. The observation that early-life exposure is necessary for lifespan extension **(Fig. 1G-H)** suggests that redox-driven metabolic adaptations may act early in life to prime these transcriptional changes, ultimately conferring longevity.

### Lactate longevity depends on SIR-2.1 activity

Considering evidence that lactate affects PTMs, including acetylation and lactylation that impact immune and metabolic pathways^2,59–61^, we noted expression changes of several genes involved in protein modification in our transcriptomic profiles, in particular chromatin modifiers **(Fig. S3I)**. Therefore, we measured acetyl-lysine levels, including histone acetylation levels, in N2 worms, which revealed a significant increase of acetylation signal in the whole-cell protein extract **(Fig. 4A)**. In contrast, global lactylation levels remained unchanged **(Fig. S5A)**. Antioxidant treatment (NAC) did not alter lactate-induced acetylation **(Fig. S5B)**, suggesting either that redox signaling and acetylation operate independently or that ROS induction occurs downstream of acetylation. To further investigate the role of acetylation, we targeted *cbp- 1*/CBP1, an acetyltransferase reported also to possess lactyltransferase^62^ activity and to regulate SKN-1-dependent stress responses^63^ as well as other cellular defense pathways^64,65^. RNAi-mediated knockdown of *cbp-1* reduced the lifespan under control conditions and abolished lactate’s pro-longevity effects **(Fig. 4B)**. Instead, lactate reduced the lifespan of animals treated with *cbp-1* RNAi. These results suggest that increased protein is required for lactate-mediated longevity.

**Figure 4:**
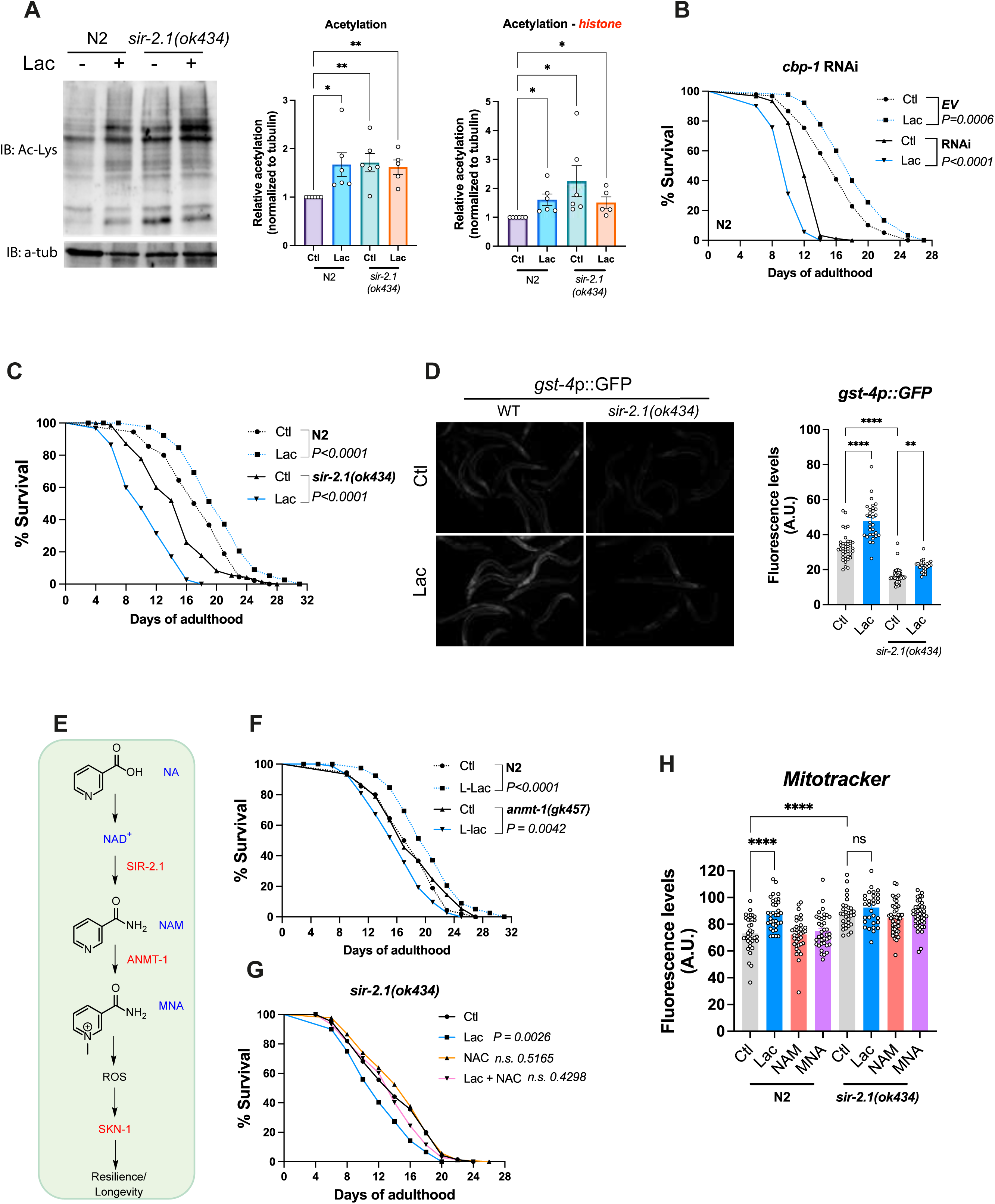
Protein acetylation regulates lactate-mediated longevity. **A)** Protein acetylation level as measured by western blot, for N2 and *sir-2.1(ok434)* animals under control and lactate-enriched diet. Tubulin is displayed as a loading control, and protein level is measured relative to control condition. **B)** Lifespan curve for N2 animals treated with *cbp-1* RNAi. Worms were grown on control RNAi, under control (black) or lactate-supplemented (blue) diets, and transferred to *cbp-1* RNAi at L4 stage. **C)** Lifespan curve for N2 (dotted lines) and *sir-2.1(ok434) (*full lines) animals under control (black) or lactate-supplemented (blue) diets. 90 animals were scored per condition. **D)** Images of *gst-4*p::GFP reporter in wild-type and *sir-2.1(ok434)* background. Quantification *gst-4*p::GFP expression in day 1 animals under control or lactate-supplemented diet. Each dot represents one animal. **E)** Schematic of SIR-2.1-dependent ROS production via nicotinamide (NAM) and methyl icotinamide (MNA). **F)** Lifespan curve for N2 (dotted lines) and *anmt-1(gk457) (*full lines) animals under control (black) or lactate-supplemented (blue) diets. **G)** Lifespan curves for *sir-2.1(ok434)* animals. Animals were treated under control (Ctl), 10 mM lactate (Lac), 5 mM N-acetylcysteine (NAC) or 10 mM lactate + 5 mM NAC (Lac + NAC) diets. **H)** Quantification of mitotracker-H2Xros staining in N2 and *sir-2.1(ok434)* animals. Staining was applied to day 1 animals, under different treatments. Control (grey), Lactate 10 mM (blue), NAM 100 µM (red), MNA 1µM (purple) Statistics for lifespan curves were done using log-rank (Mantel–Cox) method. For each condition, 90 animals were analyzed (in triplicate), and each experiment was repeated at least twice. P values for fluorescence measure were calculated by unpaired, two-sided t-test with Welch correction. *P<0.05; **P<0.01; ****P<0.0001. ns: Not significant. Each fluorescence experiment was performed at least 3 times and between 25-40 animals were measured per condition. Acetylation levels were normalized to tubulin and represented relative to the control condition.

The observed increase in acetylation levels, together with the role of lactate in regulating NAD/NADH ratio and mitochondrial function, prompted us to explore the involvement of sirtuins, a class of NAD^+^-dependent histone deacetylases. As expected, *sir-2.1/*SIRT1 mutants exhibited baseline hyperacetylation, which was not further increased by lactate supplementation **(Fig. 4A)**. Intriguingly, lactate supplementation of *sir-2.1(ok434)* mutants not only failed to increase lifespan, but instead resulted in a significant lifespan decrease, similar to the effects of *cbp-1* RNAi **(Fig. 4C)**. This effect was specific to *sir-2.1*, as mutations in the mitochondrial sirtuins *sir-2.2* and *sir-2.3* did not impair lactate-induced lifespan extension **(Fig. S5C)**. Additionally, loss of *sir-2.1* reduced GST-4 expression under control diet and strongly suppressed lactate-induced GST-4 expression **(Fig. 4D)**. These data suggest that lactate promotes longevity via increased protein acetylation and subsequent deacetylation via SIR- 2.1, resulting in increased NAD^+^ consumption.

To determine whether SIR-2.1-dependent NAD⁺ metabolism contributes to lactate’s effects, we quantified key metabolites in the *sir-2.1*-NAD⁺ signaling cascade using high-performance liquid chromatography coupled to high-resolution mass spectrometry (HPLC–HRMS). Notably, steady-state NAD⁺ and NADH levels were unaffected by lactate supplementation **(Fig. S6A)**, indicating that the increase inNADH levels driven by lactate must be offset by increased regeneration of NAD^+^ via mitochondrial respiration. However, several NAD^+^-related metabolites were altered in lactate-treated animals. Nicotinic acid (NA) levels **(Fig. S6B)** were decreased while nicotinamide (NAM) levels increased **(Fig. S6C)**. NAM is produced from NAD^+^ during SIR-2.1-catalyzed lysine deacetylation, and thus increased NAM levels in lactate-supplemented animals could be the result of increased sirtuin activity. Additionally, kynurenine, an intermediate in the *de novo* NAD⁺ synthesis pathway, was elevated under lactate treatment **(Fig. S6D)**, possibly suggesting increased flux through this pathway **(Fig. S6E)**. Levels of 1-*N*-methylnicotinamide (MNA) **(Fig. S6F)** and nicotinamide riboside (NR) were either unchanged or not detected.

These results suggested the possibility that the lactate-driven increase in *sir-2.1* activity may promote longevity by generating reactive oxygen species (ROS), as previously demonstrated^66^ **(Fig. 4E)**. This signaling cascade involves *sir-2.1*-dependent conversion of NAD⁺ to nicotinamide (NAM) and subsequently to MNA via the enzyme amine *N*- methyltransferase (*anmt-1*). Consistent with this model, loss of *anmt-1* inverted the effect of lactate on lifespan, similar to the effects of loss of *sir-2.1* or *cbp-1* **(Fig. 4F)**. Next, we tested whether dietary supplementation with NA, NAM, or MNA affects longevity; however, none of these metabolites extended lifespan in wild-type animals in our hands **(Fig. S7A–B).** We also measured the impact of aldehyde oxidase 1 (AOx1)/*gad-3* on lactate-mediated longevity. GAD-3 was reported to extend longevity by inducing hydrogen peroxide (H_2_O_2_) levels by acting on MNA downstream of sirtuin activity^66^. However, we found that loss of *gad-3* did not block lactate-mediated longevity **(Fig. S7C)**.

Lastly, we investigated the role of ROS by combining lactate supplementation with antioxidant (NAC) treatment. Loss of *sir-2.1* blocked lactate-mediated lifespan extension and rendered mutant animals largely unresponsive to NAC **(Fig. 4G)**. Given lactate’s ability to elevate mitochondrial ROS levels in wild-type animals **(Fig. 2F)**, we hypothesized that if ROS induction depends on the *sir-2.1*-NAM/MNA axis, loss of *sir-2.1* would prevent ROS accumulation. However, mitochondrial ROS levels were already elevated in *sir-2.1(ok434)* mutants, and lactate supplementation did not further increase ROS. Additionally, supplementation with previously reported doses^66^ of NAM (100 μM) and MNA (1 μM) did not increase ROS levels in either wild-type animals or *sir-2.1* mutants **(Fig. 4H)**, nor did these metabolites induce expression of the oxidative stress reporter GST-4 **(Fig. S7E, F)**. In fact, higher concentrations of NAM (1 and 2 mM) reduced GST-4 expression **(Fig. S7E)**.

Together, these findings support a model in which lactate promotes longevity by increasing the demand for mitochondrial respiration in two ways. First, via the two-step conversion of lactate into acetyl-CoA (via pyruvate), concomitant with large amounts of NADH, which necessitates increased NAD^+^ regeneration. Second, via the increased protein acetylation induced by acetyl-CoA and CBP-1 activity. Both cascades require an increased SIR-2.1 activity, which consumes NAD^+^, to limit ROS accumulation through the activation of a SKN-1- dependent stress response and potentially act as a regulatory feedback on protein acetylation. Mitochondrial respiration thus appears to be the proximal source of the lactate-induced increase in ROS. In contrast, NAM/MNA metabolism downstream of SIR-2.1 activity is not the primary driver of stress-response activation or lifespan extension. Nonetheless, deletion of *anmt-1* diminishes lactate’s longevity-promoting effect, indicating that this methyltransferase has an additional role in lactate-mediated changes, potentially via regulation of autophagy and S-adenosylmethionine (SAM) levels^67^.

### Lactate regulates lipid-associated molecules during early adulthood

To investigate how lactate-induced redox changes influence metabolic networks, we used untargeted metabolomics on N2 animals treated with 10 mM lactate at days 1 and 5 of adulthood using HPLC–HRMS **(Fig. 5A)**. Using MS/MS molecular networking, we identified four distinct clusters representing metabolite classes consistently downregulated by lactate **(Fig. 5B)**. These metabolites shared two defining characteristics: they were lipid-derived structures with long-chain unsaturated fatty acyl groups, and they played functional roles in membrane dynamics, signaling, and mitochondrial transport.

**Figure 5:**
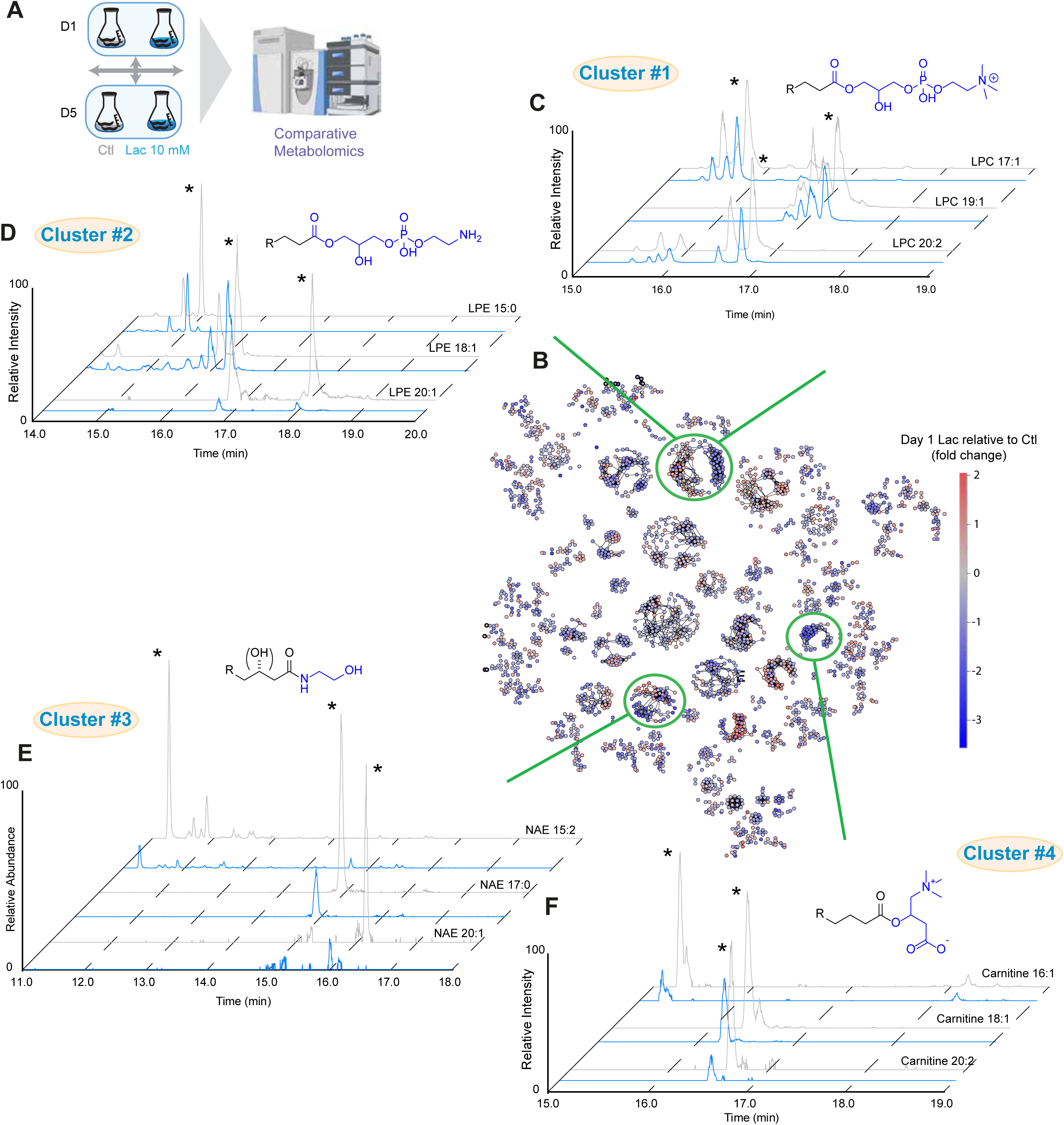
Lactate impacts lipid metabolism **A)** Schematic representation of the untargeted metabolomic pipeline used in this study **B)** Comparative metabolite network between day 1 animals treated with lactate (10 mM) vs control. Metabolite levels are represented as a Log(2) fold change over day 1 control samples. **C-F)** Chromatogram of three lactate-regulated lysophosphatidylcholine **(C)**, lysophatidylethanolamine **(D)**, N-acylethanolamine (NAE) **(E)**, and fatty-acylcarnitine **(F)**. MS2 spectra for the network were acquired in ESI(+) mode. Asterisk (*) indicates the measured peak. R in each schematic correspond of a fatty-acyl chain of variable length. Fig. S7 contains measures of all differential compounds related to this figure.

Clusters #1 and #2 were composed of lysophospholipids. Namely, we identified lysophosphatidylcholine (LPC) and lysophosphoethanolamine (LPE) (**Fig. 5C, D, S8A, B**). Lysophospholipids are membrane-derived lipids that possess hydrophilic choline or ethanolamine moieties^68^. Although their cellular function is unclear, their concentration has been shown to change during aging and inflammation^69^. Cluster #3 was found to represent *N*- acylethanolamines (NAEs) of varying chain lengths, some of which featured additional oxygenation of the fatty acid chain (**Fig. 5E, S8C, D**). These lipid-derived molecules are similar to the mammalian endocannabinoid system and have been associated with development and longevity in *C. elegans*, in a diet-dependent manner^70,71^. Finally, cluster #4 represented a series of fatty-acylcarnitine (**Fig. 5F**). Carnitine acts as a lipid carrier, and acyl-carnitine formation is a prerequisite for the transport of fatty acids into the mitochondria. Given that abundances of four dominant families of lipids were significantly decreased under lactate supplementation, we asked whether concentrations of free fatty acids were also altered by lactate treatment. We observed that most saturated and unsaturated free fatty acids were also less abundant in animals supplemented with lactate. **(Fig. S8F)**.

Next, we explored the redox dependence of these lipid-associated changes by exposing wild-type animals to paraquat (PQ), a low-dose ROS-inducing agent known to extend lifespan when administered early in life^72,73^. PQ treatment replicated most of the metabolic changes observed with lactate supplementation **(Figs. S8G–I)**, supporting the hypothesis that ROS induction drives the metabolic changes caused by lactate.

Finally, we assessed how these identified classes of molecules vary during aging. LPC levels declined significantly during aging, while LPE showed a more modest decrease **(Figs. S9A, B)**. NAEs exhibited no significant changes over time **(Fig. S9C)**, while fatty-acylcarnitine increased with age **(Fig. S8D)**. Notably, lactate supplementation did not further modulate NAE and fatty-acylcarnitine at later stages (day 5) **(Figs. S9E–H)**.

These findings suggest that lactate reprograms lipid metabolism through redox-dependent mechanisms. Lactate’s suppression of lysophospholipids, NAEs, and fatty-acylcarnitine mirrors metabolic shifts observed in lifespan-extending interventions such as caloric restriction, indicating that lipid depletion may prime stress-resilience pathways. Furthermore, the shared metabolic signatures between lactate and PQ imply that early-life ROS triggers conserved lipid remodeling processes, consistent with hormetic models of aging. Reduced levels of fatty-acylcarnitine may also limit β-oxidation, thereby lowering mitochondrial ROS production while enhancing metabolic efficiency, a hallmark of longevity-promoting pathways.

### Lactate regulates lipid metabolism

Our untargeted metabolomics findings prompted us to investigate how lactate regulates lipid metabolism to extend lifespan. Using Nile Red (NR) and Oil Red O staining, we observed that lactate supplementation increased neutral lipid storage in day 1 and day 3 adults compared to controls, though this effect diminished by day 5 **(Figs. 6A, S9A)**. Pyruvate similarly elevated fat levels in young adults **(Fig. 6B)**, suggesting that lactate and pyruvate have similar overall metabolic effects.

**Figure 6:**
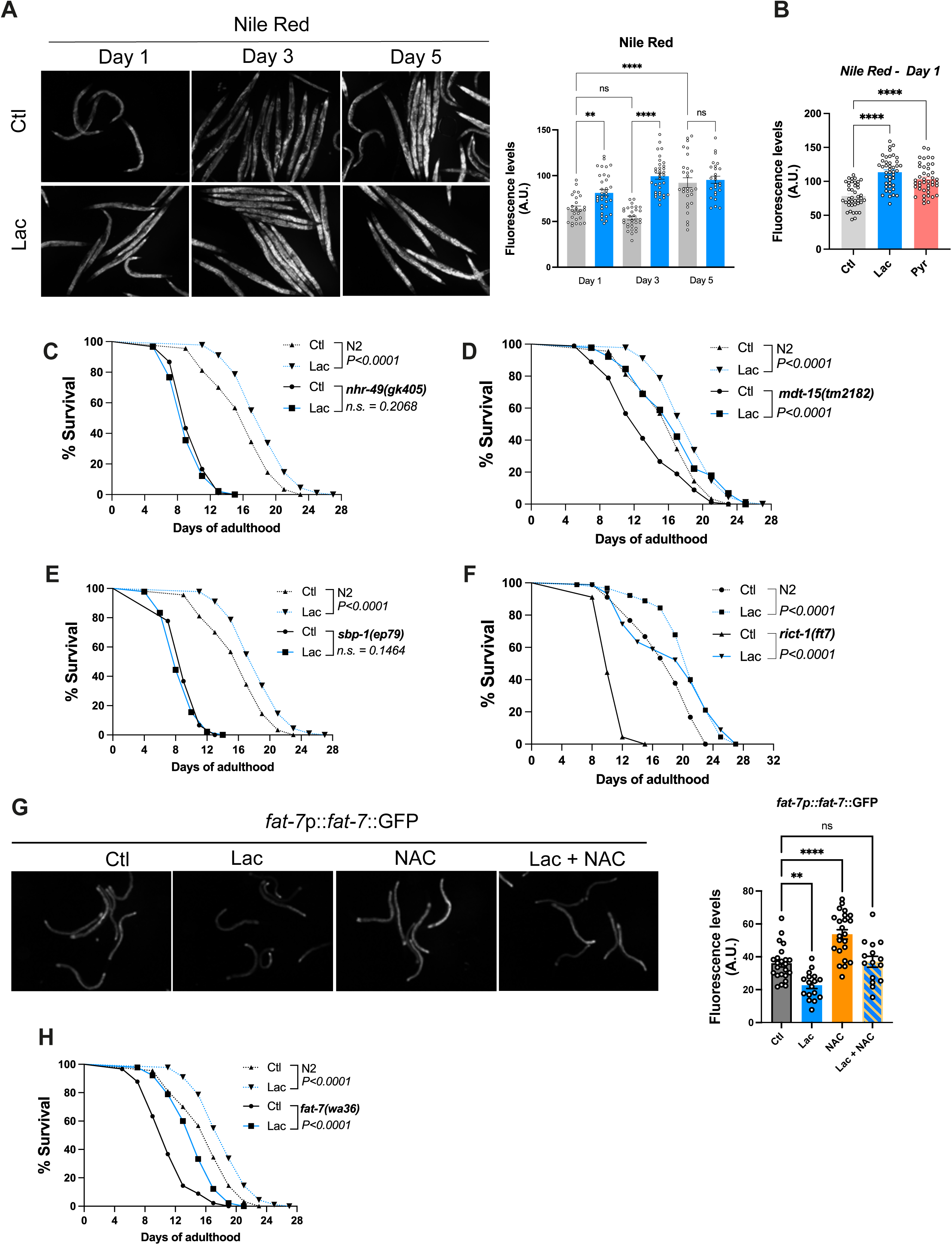
Lactate-mediated longevity is influenced by lipid regulators A) Representative Images and quantification of N2 animals stained with Nile red. Animals were grown on control or lactate-supplemented (10 mM) diet and stained at day 1, 3 and 5 of adulthood. B) Quantification of Nile red staining experiment. Animals were grown on a control, lactate (10 mM), and pyruvate (10 mM) diet and stained at day 1 post-adult. **C-F)** Lifespan curves for *nhr-49(gk405)*, *mdt-15(tm2182)*, *sbp-1(ep79)*, and *rict-1(ft7)* loss-of- function mutants under control (black) and lactate-enriched diet (blue). The dotted line corresponds to WT lifespan curves. Animals were grown on a control or lactate-supplemented diet. G) Representative images and quantification of *fat-7*::GFP reporter in day 1 animals. Animals were treated under a controlled diet, lactate (10 mM), N-acetylcysteine (5 mM), and a combination of lactate (10 mM) and N-acetylcysteine (5 mM). Between 20 and 40 animals were imaged at day 1 of adulthood. H) Lifespan curve for *fat-7(wa36)* animals under control or lactate-enriched (10 mM) diets. Statistics for lifespan curves were done using log-rank (Mantel–Cox) method. For each condition, 90 animals were analyzed (in triplicate), and each experiment was repeated at least twice. P values for fluorescence were calculated by unpaired, two-sided t-test with Welch correction. *P<0.05; **P<0.01; ****P<0.0001. ns: Not significant. Each fluorescence experiment was performed at least 3 times, and between 25 and 40 animals were measured per condition.

Fat metabolism has been associated with longevity and homeostasis in multiple ways^68,74^. Guided by our transcriptomic analysis, which showed strong changes in lipid metabolism enzymes, such as the desaturase *fat-7*/SCD1 **(Fig. 3D, Fig. S3E)**, we investigated how the loss of *nhr-49(gk405)*, *sbp-1(ep79)*, *and mdt-15(tm2182)* would impact lactate-mediated longevity. *Nhr-49*/PPARα is a conserved transcription factor that plays a central role in the regulation of lipid metabolism^75–77^ as well as survival to stress and longevity^78^. The loss of *nhr- 49* blocked lactate-mediated longevity **(Fig. 6C)**. We next examined the transcription factor *sbp-1*/SREBP1^79–81^ and co-activator mdt-15/MED15^82,83^, and only the loss of *sbp-1* abrogated lactate-driven longevity (**Fig. 6D, E**). Finally, we also assessed the role of *rict-1*/RICTOR in our phenotyping panel. RICT-1 is a key component of the TORC2 complex and has been associated with lipid metabolism and mitochondrial integrity^84–86^ and is a known negative regulator of SKN-1^87^. The other mTOR complex, TORC1, has previously been associated with lactate’s physiological effects^31^. Loss-of-function mutants for *rsks-1*/S6K and *ife-2*/eIF4E, both downstream of TORC1^88^, blocked lactate-mediated longevity (**Fig. 3**). Surprisingly, lactate supplementation dramatically extended the lifespan of *rict-1(ft7)* animals, which are normally short-lived. Both mean (15% vs 210%) and maximal lifespan (20% vs 80%) were increased to a much greater extent by lactate in *rict-1(ft7)* than in N2 animals (**Fig. 6F**). These data support that lactate-mediated longevity is dependent on lipid regulation.

Redox regulation emerged as a key modulator of lipid metabolism under lactate treatment. While lactate suppressed FAT-7 (Δ9 desaturase regulated by NHR-49 and SBP-1) expression **(Fig. 6G, S3E)**, antioxidant (NAC) supplementation restored FAT-7 levels. Combined lactate and NAC treatment resulted in intermediate FAT-7 expression **(Fig. 6G)**, indicating the presence of both redox-dependent and independent mechanisms. Conversely, FAT-6 expression remained unchanged **(Fig. S9B)**. Knockdown of *fat-7* enhanced lactate’s pro-longevity effects **(Figs. 6H, S9C)**, indicating that lactate’s suppression of desaturase activity may optimize lipid composition for stress resilience.

Neutral lipid accumulation in *sir-2.1* and *rict-1* mutants provided further mechanistic insights. Both mutants exhibited elevated baseline fat stores^84,89^, but lactate failed to modulate lipid levels in these backgrounds (**Fig. 7A, B)**. Similarly, *sir-2.1* mutants showed elevated FAT-7 expression unresponsive to lactate, while *rict-1* mutants displayed constitutively low FAT-7 levels **(Fig. 7C)**. The pronounced effect of *rict-1* mutant on lactate-mediated longevity, appeared to be dependent on redox signaling as the use of NAC, with no effects on its own, partially reduced lactate-driven longevity phenotype **(Fig. 7D)**, suggesting that TORC2 operates downstream of the redox-dependent arm of the lactate pathway. Genetic epistasis analysis further revealed that *sir-2.1* acts upstream of *rict-1*, as double mutants phenocopied the short lifespan of *rict-1* mutants, and *sir-2.1* mutants abolished lactate-induced benefits on both longevity and GST-4 activity, while the loss of *rict-1* alone increased GST-4 expression **(Fig. 7E–G)**.

**Figure 7:**
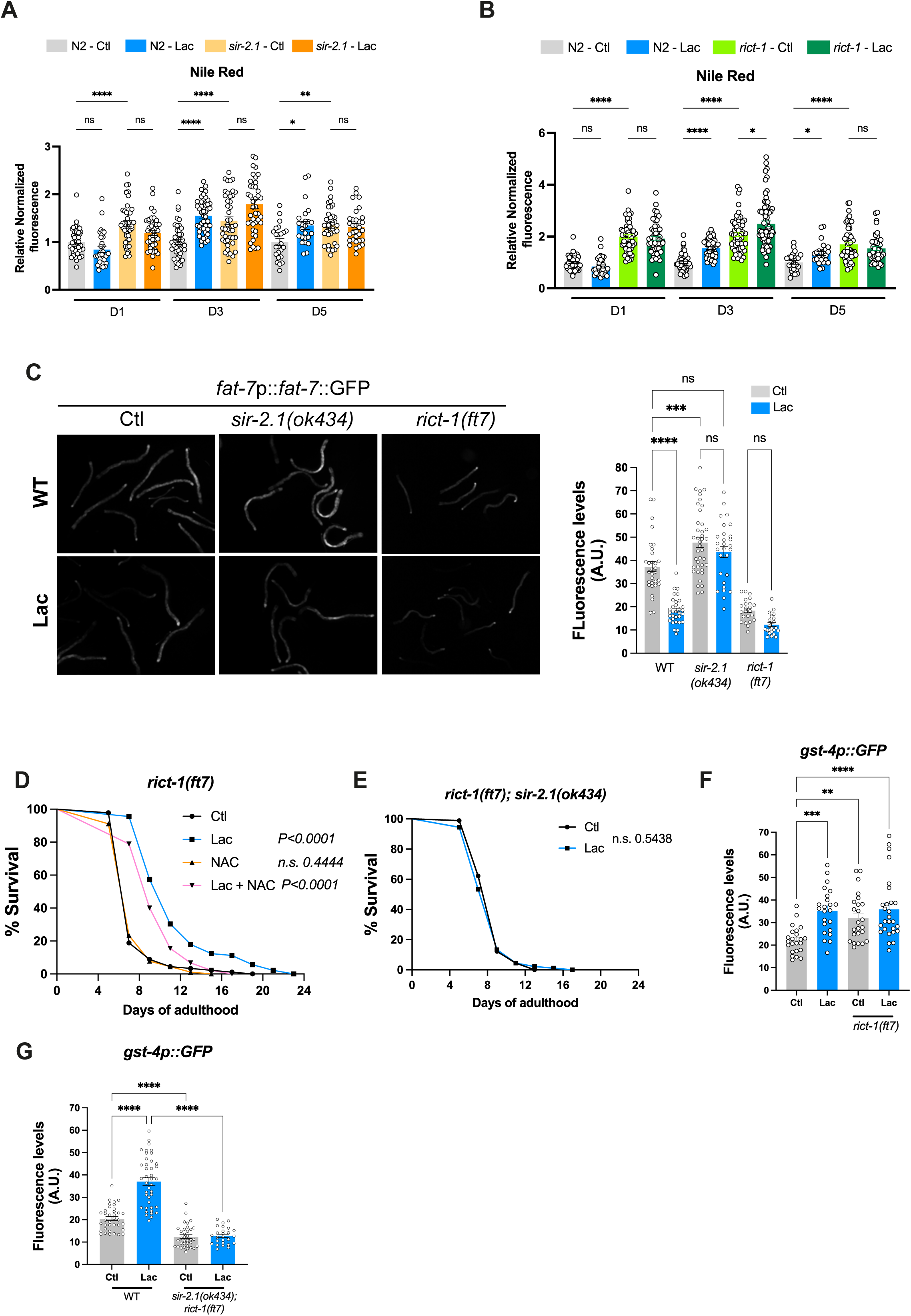
Lactate-dependent lipid changes require sir-2.1 and rict-1 **A-B)** Quantification of Nile red staining in *sir-2.1(ok434)* **(A)** and *rict-1(ft7)***(B)** under control or lactate-enriched diets. Fat content was measured at three life-stages (adult day 1, 3, and 5). 40 to 50 animals were scored per condition/ per day. C) Images of *fat-7*::GFP reporter in wild-type, *sir-2.1(ok434)* and *rict-1(ft7)* backgrounds. Animals were grown on a control or lactate-enriched diet, and the expression level was measured in day 1 adults. Quantification of *fat-7*::GFP level. Each dot represents an animal. D) Lifespan curves for *rict-1(ft7)* animals. Animals were treated under control (Ctl), 10 mM lactate (Lac), 5 mM N-acetylcysteine (NAC), or 10 mM lactate + 5 mM NAC (Lac + NAC) diets. E) Lifespan curves for *sir-2.1(ok434); rict-1(ft7)* animals. Animals were grown on a control or a lactate-enriched diet. F) Quantification of *gst-4*p::GFP reporter in WT and rict-1(ft7) mutants. Animals were grown on a control or lactate-enriched diet, and fluorescence was measured in day 1 adults. G) Quantification of *gst-4*p::GFP reporter in WT and *sir-2.1(ok434); rict-1(ft7)* background. Animals were grown on a control or lactate-enriched diet, and fluorescence was measured in day 1 adults. Statistics for lifespan curves were done using log-rank (Mantel–Cox) method. For each condition, 90 animals were analyzed (in triplicate), and each experiment was repeated at least twice. P values for fluorescence were calculated by unpaired, two-sided t-test with Welch correction. *P<0.05; **P<0.01; ***P<0.001 ****P<0.0001. ns: Not significant. Each fluorescence experiment was performed at least 3 times, and between 25 and 40 animals were measured per condition. Nile-red staining levels were normalized to animals’ size to account for the size difference between N2, *sir-2.1(ok434)*, and *rict-1(ft7)*.

### Lactate inhibits lipid metabolism to promote longevity

Next, we explored how the loss of *sir-2.1* affects levels of lipid-derived molecules regulated by lactate. We screened lactate-regulated compounds using HPLC-MS in day 1 adults **(Fig. 8A)**. The first observation was that lactate supplementation did not significantly alter the abundance of LPC, fatty-acylcarnitine, and NAE levels in the *sir-2.1* mutant background when compared to untreated *sir-2.1* control. The second observation was that LPC and fatty-acylcarnitine levels were globally reduced in untreated *sir-2.1* mutants compared to WT, while NAE levels were comparable between N2 and *sir-2.1* mutants **(Fig. 8B-D)**. A similar metabolomic approach using *rict-1* mutants revealed that LPC and fatty-acylcarnitine levels were not as significantly reduced compared to untreated WT and lactate treatment, although minimally, appeared to increase the levels of these metabolites in *rict-1* mutants **(Fig. S11)** NAE levels significantly reduced in untreated *rict-1* mutants compared to WT, and lactate treatment showed a similar trend, of increased level in *rict-1* mutants.

**Figure 8:**
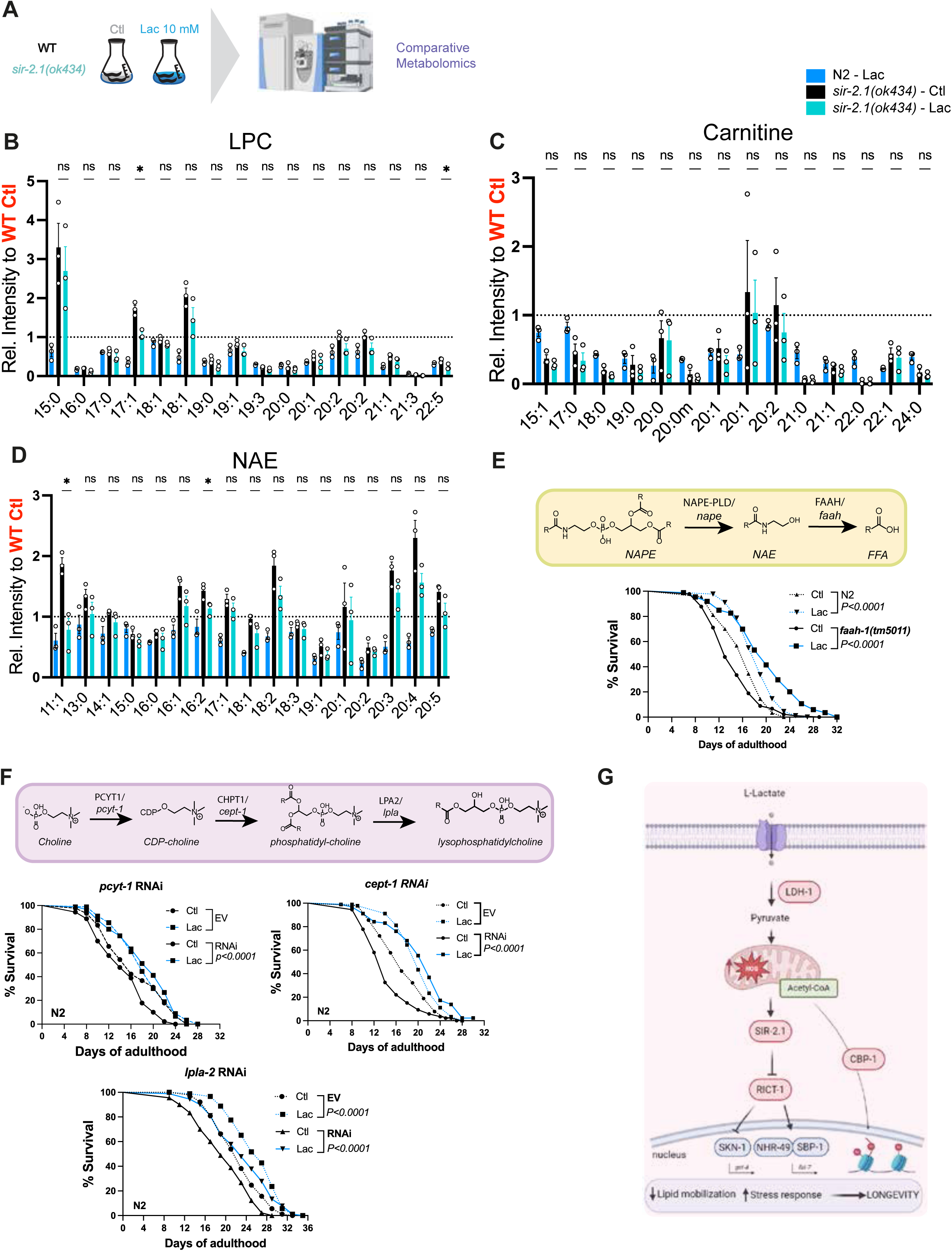
Lipid associated molecules biosynthetic pathways regulate lactate longevity. **A)** Schematic of metabolomics experiments comparing N2 and *sir-2.1(ok434)* animals on a control or lactate-enriched diet. Samples were harvested at day 1 adult stage. **B-D)** Quantification of lactate-regulated LPC **(B)**, fatty-acylcarnitine **(C)**, and N-acylethanolamine (NAE – **D**) in *sir-2.1(ok434) (teal)* mutants. F) Schematic of NAE biosynthesis pathway and lifespan curves of *faah-1(tm5011)* and *faah- 4(lt121)* animals grown on control diet or lactate (10 mM). Dotted lines correspond to wildtype curves. 90 animals were scored per condition. G) Schematic of LPC/LPE biosynthesis pathway and lifespan curves for N2 animals treated with L4440 (empty vector), *cept-1*, *pcyt-1, or lpla-2* RNAi. Animals were grown on a control diet or lactate (10 mM). 90 animals were scored per condition. Statistics for lifespan curves were done using log-rank (Mantel–Cox) method. For each condition, 90 animals were analyzed (in triplicate), and each experiment was repeated at least twice. P-values for metabolite levels were calculated by an unpaired, two-sided t-test with Welch correction. *P<0.05; **P<0.01; ****P<0.0001. ns: Not significant. Metabolite measures represent one experiment, performed in triplicate. Each experiment was performed three times. Graphical summary of the proposed model from this study, highlighting the key mechanisms and significant findings.

To assess the functional relevance of these lactate-induced lipid changes, we investigated key biosynthetic pathways involved in NAE and phospholipid metabolism **(Fig. 8E, F)**. NAEs are synthesized by N-acyl phosphatidyl ethanolamine-specific phospholipase D (NAPE-PLD/*nape*) and hydrolyzed by fatty-acyl amide hydrolase (FAAH/*faah*)^70^. Interestingly, *nape- 1* and *nape-2* lof mutants did not significantly affect lactate-mediated longevity **(Fig. S12A, B)**, possibly due to enzymatic redundancy. However, the *faah-1(tm5011)* lof mutant exhibited enhanced sensitivity to lactate supplementation. In *faah-1(tm5011)* mutants, lactate treatment increased mean lifespan by 54% compared to 18% in N2 controls **(Fig. 8E)**. In contrast, the *faah-4* mutant, which preferentially hydrolyzes 2-arachidonylglycerol (2-AG), did not display increased sensitivity to lactate **(Fig. S12C)**.

RNAi-mediated knockdown of genes involved in lysophospholipid synthesis^81^ revealed that disruption of this pathway amplifies lactate’s pro-longevity effects. Specifically, targeting *pcyt- 1* (PCYT1) or *cept-1* (CHPT1), key enzymes in the CDP-choline pathway^82^, and regulated by lactate **(Fig. 3F)**, markedly enhanced the lifespan extension induced by lactate. In contrast, knockdown of *lpla-2*, a lysosomal phospholipase A2, had no significant effect. The mean lifespan increase under lactate supplementation was 85% in *pcyt-1*, 57% in *cept-1*, and *21% in lpla-2* RNAi-treated worms, compared to an 18% increase in control vector-treated animals **(Figure 8F)**. These findings suggest that suppression of phospholipid synthesis sensitizes animals to lactate, supporting the idea that downregulation of lipid mobilization contributes to its pro-longevity mechanism.

Together, our findings support a model in which lactate-driven redox changes modulate lipid metabolism and regulate longevity. We propose that lactate-driven mitochondrial respiration generates ROS, which initiates a signaling cascade involving protein acetylation, activation of SIR-2.1, and metabolic remodeling. This cascade acts on downstream targets like RICT-1 to reduce lipid mobilization, ultimately leading to lifespan extension **(Fig. 8G)**. These results provide novel insights into the interplay between lactate metabolism, redox signaling, and lipid regulation in the control of longevity. Further studies are needed to elucidate the molecular details of this pathway and assess its relevance in mammalian aging.

### Discussion and conclusion

Lactate is a potent signaling molecule, involved in both physiological and pathological contexts^90^. We previously demonstrated that high concentrations of lactate enhance stress resilience through a hormetic action involving a mild increase in reactive oxygen species (ROS)^36^. In this study, we investigated the mechanisms underlying lactate-mediated longevity and found that lactate promotes lifespan and healthspan in *C. elegans* through distinct signaling pathways. A longstanding debate is how lactate influences cellular processes, through direct signaling or via its conversion to pyruvate. Our findings reveal that the oxidation of lactate to pyruvate via LDH-1 is essential for its longevity-promoting effects. This process also further generates NADH, which appears to play a critical role in mediating downstream effects. Pyruvate is metabolized further to produce acetyl-CoA and CO_2_, which generates an additional equivalent of NADH. Our finding that reducing NADH levels by expressing *Lb*NOX abolishes lactate-driven longevity strongly suggests that the build-up of NADH resulting from lactate treatment plays a central role. Regulation of the NAD^+^/NADH ratio is critical for metabolic homeostasis, and thus, the increase of NADH levels driven by lactate must be offset by increased regeneration of NAD^+^ via mitochondrial respiration.

Our proposed longevity mechanism (**Fig. 8G**) links increased ROS levels resulting from increased mitochondrial respiration and increased protein acetylation due to elevated acetyl-CoA levels (via pyruvate) with increased SIR-2.1 activity, stress response activation, and changes in lipid metabolism **(Fig. 8G)**. While NAD^+^, the substrate of SIR-2.1/SIRT1, has been previously associated with the physiological impacts of lactate^91,92^, our finding that lactate-mediated longevity is strictly dependent on *sir-2.1* was unexpected. Deletion of *sir-2.1* results in starkly increased protein acetylation levels, which appear to be detrimental, given that lactate supplementation significantly reduces the lifespan of *sir-2.1* mutants. This suggests that the pro-longevity effect of lactate hinges on increased protein deacetylation by *sir-2.1*. The signaling cascade proposed here also requires the methyltransferase *anmt-1,* which methylates NAM, derived from *sir-2.1* activity, and produces MNA. In a previous study^66^, *anmt- 1*-derived MNA was reported to increase ROS production via the oxidase GAD-3, leading to activation of stress response pathways, including *gst-4*. However, in our hands, MNA did not significantly increase ROS or *gst-4* activity, and the loss of *gad-3* did not prevent lactate-driven longevity. This suggests that ANMT-1 may methylate additional substrates leading to activation of *gst-4* or that *anmt-1*-dependent conversion of NAM to MNA regulates NAM metabolism by additional mechanisms.

Taken together, our model proposes that lactate supplementation increases NADH levels, triggering increased mitochondrial respiration to regenerate NAD+ as well as protein hyperacetylation that activates SIR-2.1, which in turn initiates a signaling cascade that activates *skn-1*/Nrf-2 and promotes the expression of detoxification enzymes such as GST-4 as well as changes in lipid metabolism. This model is supported further by our finding that knocking down *cbp-1*/CBP1, a key acyltransferase previously implicated stress response^63,65,95^, not only abolished lactate-mediated longevity, but also, similar to what we observed in *sir-2.1* mutants, led to reduced lifespan in response to lactate. While *cbp-1* has previously been implicated with protein lactylation, our results indicate that lactylation levels were not significantly increased after lactate treatment, in contrast to a drastic increase in acetylation levels. Clarifying the exact role of *cbp-1* for lactate-dependent changes in protein acylation levels will require more detailed characterization of the substrate scope of CBP-1 as well as of the changes to the protein acylation landscape in response to lactate treatment.

Lactate is known for its protective effects^96,97^, but chronic high levels can be harmful^98–100^. In our study, lactate conferred SKN-1/Nrf2–dependent protection, markedly improving mid-life survival in N2 animals, with a smaller effect on maximum lifespan **(Fig. 2)**. Lactate sustained GST4 activity through day 5, which may underlie its later detrimental impact, as prolonged SKN-1 activation is linked to reduced lifespan^101^. This biphasic effect parallels other interventions that are beneficial early but deleterious later in life^102,103^ and matches reports on the dual role of early reactive oxygen species exposure in promoting resilience and longevity^72,104^. Furthermore, lactate-induced protein acylation may play a role, with transient acetylation aiding stress adaptation, but persistent acetylation accelerating aging^105,106^.

The final key observation in this study was a marked shift in metabolic pathways under lactate supplementation. Our untargeted metabolomics analysis revealed that changes in energy metabolism were detectable as early as day 1, preceding the transcriptomic alterations observed at day 5. We propose that lactate stimulates mitochondrial activity, thereby increasing protein acetylation and subsequently suppressing the expression of metabolic genes, including those related to lipid mobilization. This aligns with previous findings linking elevated acetylation to downregulation of metabolism^107^ through sirtuin activity^108^. Although lactate’s physiological role in lipid regulation is not fully understood^109,110^, we found that supplementation decreased levels of several lipid-derived metabolites **(Fig. 5)** and required the lipid-associated factors NHR-49 and RICT-1 for lifespan extension. Notably, RICT-1 has not been implicated in lactate signaling before; it integrates nutrient cues and regulates metabolism, longevity, and Nrf-2/skn-1 activity. Interestingly, *rict-1* knockdown, despite reducing lifespan on its own, enhanced lactate-mediated longevity **(Fig. 6)**.

Our investigation into lipid changes associated with lactate-driven longevity demonstrated that RNAi knockdown of the biosynthetic pathways for lysophospholipids (LPC/LPE) and for *N*- acylethanolamines (NAE) produced distinct effects. Specifically, inhibition of LPC/LPE synthesis and suppression of NAE catabolism both significantly enhanced lactate-mediated lifespan extension **(Fig. 8)**. These findings indicate that lactate promotes healthy aging, at least in part, by modulating lipid metabolic pathways.

We propose a mechanistic model in which lactate promotes mitochondrial activity, resulting in increased metabolic activity through the production of pyruvate and acetyl-CoA. Increased mitochondrial function induces ROS and leads to increased protein acetylation, which in turn activates SIR-2.1 (**Fig. 8G**). This signaling cascade triggers cellular detoxification mechanisms mediated by SIR-2.1 and SKN-1. Concurrently, lactate, possibly through SIR- 2.1-mediated pathways, downregulates RICT-1, reducing lipid metabolism. Together, these coordinated metabolic and redox adaptations, occurring particularly in later stages of life, may contribute to enhanced longevity.

### Study limitations

This model offers a framework for understanding the complex interplay between lactate metabolism, stress response pathways, and longevity regulation. However, several questions remain. First, the precise molecular mechanisms by which lactate-induced ROS promotes longevity are still unclear. Although our transcriptomic analyses revealed gene expression changes at later life stages, we hypothesize that early effects of lactate may be mediated through post-translational modifications, including acetylation. To address this, the effects of lactate at different life stages warrant further investigation. Second, while we investigated the systemic effects of lactate, its temporal and spatial dynamics remain to be elucidated. Because metabolic and signaling characteristics vary across tissues, it is crucial to determine whether lactate’s redox and lipid effects occur in the same or distinct tissues.

## Supporting information

Table III

## Acknowledgments

We thank Dr. Sylvia Lee and Dr. Bennett Fox for their technical assistance and constructive comments. Thanks to the *C. elegans* Genetic Center and National Bioresource Project for the different strains used in this study. Support for this study was provided by King Abdullah University of Science and Technology (PJM), Canadian Institutes of Health Research (AGE-477545) (AT), NIH R00GM140217 (JDM) and NIH R35GM131877 (FCS).

## Author contributions

AT, PJN and JDM performed the experiments. AT, HF, FCS and PJM designed the experiments. All the authors commented on the results and reviewed the manuscript.

## References

1. Fantin, V.R., St-Pierre, J., and Leder, P. (2006). Attenuation of LDH-A expression uncovers a link between glycolysis, mitochondrial physiology, and tumor maintenance. Cancer Cell 9, 425–434. 10.1016/j.ccr.2006.04.023.

2. An, Y.J., Jo, S., Kim, J.-M., Kim, H.S., Kim, H.Y., Jeon, S.-M., Han, D., Yook, J.I., Kang, K.W., and Park, S. (2023). Lactate as a major epigenetic carbon source for histone acetylation via nuclear LDH metabolism. Exp. Mol. Med., 1–10. 10.1038/s12276-023-01095-w.

3. Li, H., Rai, M., Buddika, K., Sterrett, M.C., Luhur, A., Mahmoudzadeh, N.H., Julick, C.R., Pletcher, R.C., Chawla, G., Gosney, C.J., et al. (2019). Lactate dehydrogenase and glycerol- 3-phosphate dehydrogenase cooperatively regulate growth and carbohydrate metabolism during Drosophila melanogaster larval development. Development 146, dev175315. 10.1242/dev.175315.

4. Lin, Y., Wang, Y., and Li, P. (2022). Mutual regulation of lactate dehydrogenase and redox robustness. Front. Physiol. 13, 1038421. 10.3389/fphys.2022.1038421.

5. Margineanu, M.B., Mahmood, H., Fiumelli, H., and Magistretti, P.J. (2018). L-Lactate Regulates the Expression of Synaptic Plasticity and Neuroprotection Genes in Cortical Neurons: A Transcriptome Analysis. Front Mol Neurosci 11, 375. 10.3389/fnmol.2018.00375.

6. Yang, J., Ruchti, E., Petit, J.-M., Jourdain, P., Grenningloh, G., Allaman, I., and Magistretti, P.J. (2014). Lactate promotes plasticity gene expression by potentiating NMDA signaling in neurons. Proc National Acad Sci 111, 12228–12233. 10.1073/pnas.1322912111.

7. Suzuki, A., Stern, S.A., Bozdagi, O., Huntley, G.W., Walker, R.H., Magistretti, P.J., and Alberini, C.M. (2011). Astrocyte-Neuron Lactate Transport Is Required for Long-Term Memory Formation. Cell 144, 810–823. 10.1016/j.cell.2011.02.018.

8. Fiumelli, H., Herrera-López, G., Lemtiri-Chlieh, F., Mottier, L., Girgis, J., Ben-Adiba, C., Jourdain, P., Carrano, N., Mahmood, H., Ooi, A., et al. (2024). Lactate potentiates NMDA receptor currents via an intracellular redox mechanism targeting cysteines in the C-terminal domain of GluN2B subunits: implications for synaptic plasticity. bioRxiv, 2024.11.21.624499. 10.1101/2024.11.21.624499.

9. Bozzo, L., Puyal, J., and Chatton, J.-Y. (2013). Lactate Modulates the Activity of Primary Cortical Neurons through a Receptor-Mediated Pathway. Plos One 8, e71721. 10.1371/journal.pone.0071721.

10. Abrantes, H. de C., Briquet, M., Schmuziger, C., Restivo, L., Puyal, J., Rosenberg, N., Rocher, A.-B., Offermanns, S., and Chatton, J.-Y. (2019). The Lactate Receptor HCAR1 Modulates Neuronal Network Activity through the Activation of G α and G βγ Subunits. J Neurosci 39, 4422–4433. 10.1523/jneurosci.2092-18.2019.

11. Herrera-López, G., and Galván, E.J. (2018). Modulation of hippocampal excitability via the Hydroxycarboxylic Acid Receptor 1. Hippocampus, 1–38. 10.1002/hipo.22958.

12. Moreno-Yruela, C., Zhang, D., Wei, W., Bæk, M., Liu, W., Gao, J., Danková, D., Nielsen, A.L., Bolding, J.E., Yang, L., et al. (2022). Class I histone deacetylases (HDAC1–3) are histone lysine delactylases. Sci. Adv. 8, eabi6696. 10.1126/sciadv.abi6696.

13. Zhang, N., Jiang, N., Yu, L., Guan, T., Sang, X., Feng, Y., Chen, R., and Chen, Q. (2021). Protein Lactylation Critically Regulates Energy Metabolism in the Protozoan Parasite Trypanosoma brucei. Front Cell Dev Biol 9, 719720. 10.3389/fcell.2021.719720.

14. Hagihara, H., Shoji, H., Otabi, H., Toyoda, A., Katoh, K., Namihira, M., and Miyakawa, T. (2021). Protein lactylation induced by neural excitation. CellReports 37, 109820. 10.1016/j.celrep.2021.109820.

15. Huang, A., Li, Y., Duan, J., Guo, S., Cai, X., Zhang, X., Long, H., Ren, W., and Xie, Z. (2022). Metabolomic, proteomic and lactylated proteomic analyses indicate lactate plays important roles in maintaining energy and C:N homeostasis in Phaeodactylum tricornutum. Biotechnology Biofuels Bioprod 15, 61. 10.1186/s13068-022-02152-8.

16. Jiang, J., Huang, D., Jiang, Y., Hou, J., Tian, M., Li, J., Sun, L., Zhang, Y., Zhang, T., Li, Z., et al. (2021). Lactate Modulates Cellular Metabolism Through Histone Lactylation- Mediated Gene Expression in Non-Small Cell Lung Cancer. Front. Oncol. 11, 647559. 10.3389/fonc.2021.647559.

17. Husain, Z., Huang, Y., Seth, P., and Sukhatme, V.P. (2013). Tumor-Derived Lactate Modifies Antitumor Immune Response: Effect on Myeloid-Derived Suppressor Cells and NK Cells. J Immunol 191, 1486–1495. 10.4049/jimmunol.1202702.

18. Peter, K., Rehli, M., Singer, K., Renner-Sattler, K., and Kreutz, M. (2015). Lactic acid delays the inflammatory response of human monocytes. Biochem Bioph Res Co *457*, 412418. 10.1016/j.bbrc.2015.01.005.

19. Hoque, R., Farooq, A., Ghani, A., Gorelick, F., and Mehal, W.Z. (2014). Lactate Reduces Liver and Pancreatic Injury in Toll-Like Receptor– and Inflammasome-Mediated Inflammation via GPR81-Mediated Suppression of Innate Immunity. Gastroenterology 146, 1763–1774. 10.1053/j.gastro.2014.03.014.

20. Buscemi, L., Price, M., Castillo-González, J., Chatton, J.-Y., and Hirt, L. (2022). Lactate Neuroprotection against Transient Ischemic Brain Injury in Mice Appears Independent of HCAR1 Activation. Metabolites 12, 465. 10.3390/metabo12050465.

21. Berthet, C., Lei, H., Thevenet, J., Gruetter, R., Magistretti, P.J., and Hirt, L. (2009). Neuroprotective role of lactate after cerebral ischemia. Journal of Cerebral Blood Flow & Metabolism 29, 1780–1789. 10.1038/jcbfm.2009.97.

22. Berthet, C., Castillo, X., Magistretti, P.J., and Hirt, L. (2012). New Evidence of Neuroprotection by Lactate after Transient Focal Cerebral Ischaemia: Extended Benefit after Intracerebroventricular Injection and Efficacy of Intravenous Administration. Cerebrovasc Dis 34, 329–335. 10.1159/000343657.

23. Bastian, C., Zerimech, S., Nguyen, H., Doherty, C., Franke, C., Faris, A., Quinn, J., and Baltan, S. (2022). Aging astrocytes metabolically support aging axon function by proficiently regulating astrocyte-neuron lactate shuttle. Exp Neurol 357, 114173. 10.1016/j.expneurol.2022.114173.

24. Mayorga-Weber, G., Rivera, F.J., and Castro, M.A. (2022). Neuron-glia (mis)interactions in brain energy metabolism during aging. J Neurosci Res. 10.1002/jnr.25015.

25. Limbad, C., Oron, T.R., Alimirah, F., Davalos, A.R., Tracy, T.E., Gan, L., Desprez, P.- Y., and Campisi, J. (2020). Astrocyte senescence promotes glutamate toxicity in cortical neurons. Plos One 15, e0227887. 10.1371/journal.pone.0227887.

26. Cohen, J., and Torres, C. (2019). Astrocyte senescence: Evidence and significance. Aging Cell 18, e12937. 10.1111/acel.12937.

27. Jha, M.K., Lee, Y., Liu, Y., Rothstein, J.D., and Morrison, B.M. (2019). Monocarboxylate transporter 1 in Schwann cells contributes to maintenance of sensory nerve myelination during aging. Glia 68, 161–177. 10.1002/glia.23710.

28. Jourdain, P., Rothenfusser, K., Ben-Adiba, C., Allaman, I., Marquet, P., and Magistretti, P.J. (2018). Dual action of L-Lactate on the activity of NR2B-containing NMDA receptors: from potentiation to neuroprotection. Sci. Rep. 8, 1–16. 10.1038/s41598-018-31534-y.

29. Jourdain, P., Allaman, I., Rothenfusser, K., Fiumelli, H., Marquet, P., and Magistretti, P.J. (2016). L-Lactate protects neurons against excitotoxicity: implication of an ATP- mediated signaling cascade. Sci. Rep. 6, 21250. 10.1038/srep21250.

30. Llorente-Folch, I., Rueda, C.B., Perez-Liebana, I., Satrustegui, J., and Pardo, B. (2016). L-Lactate-Mediated Neuroprotection against Glutamate-Induced Excitotoxicity Requires ARALAR/AGC1. Journal of Neuroscience 36, 4443–4456. 10.1523/jneurosci.3691-15.2016.

31. Nikooie, R., Moflehi, D., and Zand, S. (2020). Lactate regulates autophagy through ROS- mediated activation of ERK1/2/m-TOR/p-70S6K pathway in skeletal muscle. Journal of Cell Communication and Signaling 394, 1–17. 10.1007/s12079-020-00599-8.

32. Galardo, M.N., Regueira, M., Riera, M.F., Pellizzari, E.H., Cigorraga, S.B., and Meroni, S.B. (2014). Lactate Regulates Rat Male Germ Cell Function through Reactive Oxygen Species. PLoS ONE 9, e88024. 10.1371/journal.pone.0088024.t002.

33. Hashimoto, T., Hussien, R., Oommen, S., Gohil, K., and Brooks, G.A. (2007). Lactate sensitive transcription factor network in L6 cells: activation of MCT1 and mitochondrial biogenesis. The FASEB Journal 21, 2602–2612. 10.1096/fj.07-8174com.

34. Coco, M., Caggia, S., Musumeci, G., Perciavalle, V., Graziano, A.C.E., Pannuzzo, G., and Cardile, V. (2012). Sodium L-lactate differently affects brain-derived neurothrophic factor, inducible nitric oxide synthase, and heat shock protein 70 kDa production in human astrocytes and SH-SY5Y cultures. J. Neurosci. Res. 91, 313–320. 10.1002/jnr.23154.

35. Zelenka, J., Dvořák, A., and Alán, L. (2015). L-Lactate Protects Skin Fibroblasts against Aging-Associated Mitochondrial Dysfunction viaMitohormesis. Oxidative Medicine and Cellular Longevity 2015, 1–14. 10.1155/2015/351698.

36. Tauffenberger, A., Almustafa, S., and Magistretti, P.J. (2019). Lactate and pyruvate promote oxidative stress resistance through hormetic ROS signaling. Cell Death and Disease 10, 653. 10.1038/s41419-019-1877-6.

37. Liu, L., Zhang, K., Sandoval, H., Yamamoto, S., Jaiswal, M., Sanz, E., Li, Z., Hui, J., Graham, B.H., Quintana, A., et al. (2015). Glial lipid droplets and ROS induced by mitochondrial defects promote neurodegeneration. Cell 160, 177–190. 10.1016/j.cell.2014.12.019.

38. Liu, L., MacKenzie, K.R., Putluri, N., Maletić-Savatić, M., and Bellen, H.J. (2017). The Glia-Neuron Lactate Shuttle and Elevated ROS Promote Lipid Synthesis in Neurons and Lipid Droplet Accumulation in Glia via APOE/D. Cell Metabolism, 1–36. 10.1016/j.cmet.2017.08.024.

39. Bolger, A.M., Lohse, M., and Usadel, B. (2014). Trimmomatic: a flexible trimmer for Illumina sequence data. Bioinformatics 30, 2114–2120. 10.1093/bioinformatics/btu170.

40. Dobin, A., Davis, C.A., Schlesinger, F., Drenkow, J., Zaleski, C., Jha, S., Batut, P., Chaisson, M., and Gingeras, T.R. (2012). STAR: ultrafast universal RNA-seq aligner. Bioinformatics 29, 15–21. 10.1093/bioinformatics/bts635.

41. Liao, Y., Smyth, G.K., and Shi, W. (2014). featureCounts: an efficient general purpose program for assigning sequence reads to genomic features. Bioinformatics 30, 923–930. 10.1093/bioinformatics/btt656.

42. Love, M.I., Huber, W., and Anders, S. (2014). Moderated estimation of fold change and dispersion for RNA-seq data with DESeq2. Genome Biol. 15, 31–21. 10.1186/s13059-014-0550-8.

43. Holdorf, A.D., Higgins, D.P., Hart, A.C., Boag, P.R., Pazour, G.J., Walhout, A.J.M., and Walker, A.K. (2020). WormCat: An Online Tool for Annotation and Visualization of Caenorhabditis elegans Genome-Scale Data. Genetics 214, 279–294. 10.1534/genetics.119.302919.

44. Stiernagle, T. (2006). Maintenance of C. elegans. WormBook, 1–11. 10.1895/wormbook.1.101.1.

45. Helf, M.J., Fox, B.W., Artyukhin, A.B., Zhang, Y.K., and Schroeder, F.C. (2022). Comparative metabolomics with Metaboseek reveals functions of a conserved fat metabolism pathway in C. elegans. Nat Commun 13, 782. 10.1038/s41467-022-28391-9.

46. Titov, D.V., Cracan, V., Goodman, R.P., Peng, J., Grabarek, Z., and Mootha, V.K. (2016). Complementation of mitochondrial electron transport chain by manipulation of the NAD+/NADH ratio. Science 352, 231–235. 10.1126/science.aad4017.

47. Liu, S., Fu, S., Wang, G., Cao, Y., Li, L., Li, X., Yang, J., Li, N., Shan, Y., Cao, Y., et al. (2021). Glycerol-3-phosphate biosynthesis regenerates cytosolic NAD+ to alleviate mitochondrial disease. Cell Metabolism 33, 1974–1987.e9. 10.1016/j.cmet.2021.06.013.

48. Link, C.D., and Johnson, C.J. (2002). [42] Reporter Transgenes for Study of Oxidant Stress in Caenorhabditis elegans. Methods Enzym. 353, 497–505. 10.1016/s0076-6879(02)53072-x.

49. Leiers, B., Kampkötter, A., Grevelding, C.G., Link, C.D., Johnson, T.E., and Henkle- Dührsen, K. (2003). A stress-responsive glutathione S-transferase confers resistance to oxidative stress in Caenorhabditis elegans. Free Radic. Biol. Med. 34, 1405–1415. 10.1016/s0891-5849(03)00102-3.

50. Paek, J., Lo, J.Y., Narasimhan, S.D., Nguyen, T.N., Glover-Cutter, K., Robida-Stubbs, S., Suzuki, T., Yamamoto, M., Blackwell, T.K., and Curran, S.P. (2012). Mitochondrial SKN- 1/Nrf Mediates a Conserved Starvation Response. Cell Metab 16, 526–537. 10.1016/j.cmet.2012.09.007.

51. Tullet, J.M.A., Hertweck, M., An, J.H., Baker, J., Hwang, J.Y., Liu, S., Oliveira, R.P., Baumeister, R., and Blackwell, T.K. (2008). Direct inhibition of the longevity-promoting factor SKN-1 by insulin-like signaling in C. elegans. Cell 132, 1025–1038. 10.1016/j.cell.2008.01.030.

52. Gusarov, I., Shamovsky, I., Pani, B., Gautier, L., Eremina, S., Katkova-Zhukotskaya, O., Mironov, A., Makarov, A.А., and Nudler, E. (2021). Dietary thiols accelerate aging of C. elegans. Nature Communications, 1–14. 10.1038/s41467-021-24634-3.

53. Fletcher, M., Tillman, E.J., Butty, V.L., Levine, S.S., and Kim, D.H. (2019). Global transcriptional regulation of innate immunity by ATF-7 in C. elegans. PLoS Genet. 15, e1007830. 10.1371/journal.pgen.1007830.

54. Cai, X., Ng, C.P., Jones, O., Fung, T.S., Ryu, K.W., Li, D., and Thompson, C.B. (2023). Lactate activates the mitochondrial electron transport chain independently of its metabolism. Mol. Cell. 10.1016/j.molcel.2023.09.034.

55. Dall, K.B., Havelund, J.F., Harvald, E.B., Witting, M., and Faergeman, N.J. (2021). HLH-30-dependent rewiring of metabolism during starvation in C. elegans. Aging Cell, e13342. 10.1111/acel.13342.

56. Lapierre, L.R., Filho, C.D.D.M., McQuary, P.R., Chu, C.-C., Visvikis, O., Chang, J.T., Gelino, S., Ong, B., Davis, A.E., Irazoqui, J.E., et al. (2013). The TFEB orthologue HLH-30 regulates autophagy and modulates longevity in Caenorhabditis elegans. Nature Communications 4, 2267. 10.1038/ncomms3267.

57. Ju, S., Chen, H., Wang, S., Lin, J., Ma, Y., Aroian, R.V., Peng, D., and Sun, M. (2022). C. elegans monitor energy status via the AMPK pathway to trigger innate immune responses against bacterial pathogens. Commun Biology 5, 643. 10.1038/s42003-022-03589-1.

58. Kenyon, C.J. (2010). The genetics of ageing. Nature 464, 504–512. 10.1038/nature08980.

59. Huang, Y., Luo, G., Peng, K., Song, Y., Wang, Y., Zhang, H., Li, J., Qiu, X., Pu, M., Liu, X., et al. (2024). Lactylation stabilizes TFEB to elevate autophagy and lysosomal activity. J. Cell Biol. 223, e202308099. 10.1083/jcb.202308099.

60. Zhang, D., Tang, Z., Huang, H., Zhou, G., Cui, C., Weng, Y., Liu, W., Kim, S., Lee, S., Perez-Neut, M., et al. (2019). Metabolic regulation of gene expression by histone lactylation. Nature 574, 1–25. 10.1038/s41586-019-1678-1.

61. Pan, R.-Y., He, L., Zhang, J., Liu, X., Liao, Y., Gao, J., Liao, Y., Yan, Y., Li, Q., Zhou, X., et al. (2022). Positive feedback regulation of microglial glucose metabolism by histone H4 lysine 12 lactylation in Alzheimer’s disease. Cell Metab. 10.1016/j.cmet.2022.02.013.

62. Liu, X., Zhang, Y., Li, W., and Zhou, X. (2022). Lactylation, an emerging hallmark of metabolic reprogramming: Current progress and open challenges. Frontiers Cell Dev Biology 10, 972020. 10.3389/fcell.2022.972020.

63. Ganner, A., Gerber, J., Ziegler, A.-K., Li, Y., Kandzia, J., Matulenski, T., Kreis, S., Breves, G., Klein, M., Walz, G., et al. (2019). CBP-1/p300 acetyltransferase regulates SKN- 1/Nrf cellular levels, nuclear localization, and activity in C. elegans. Exp Gerontol 126, 110690. 10.1016/j.exger.2019.110690.

64. Zhou, L., He, B., Deng, J., Pang, S., and Tang, H. (2019). Histone acetylation promotes long-lasting defense responses and longevity following early life heat stress. PLoS Genet 15, e1008122. 10.1371/journal.pgen.1008122.

65. Li, T.Y., Sleiman, M.B., Li, H., Gao, A.W., Mottis, A., Bachmann, A.M., Alam, G.E., Li, X., Goeminne, L.J.E., Schoonjans, K., et al. (2021). The transcriptional coactivator CBP/p300 is an evolutionarily conserved node that promotes longevity in response to mitochondrial stress. Nature Aging, 1–26. 10.1038/s43587-020-00025-z.

66. Schmeisser, K., Mansfeld, J., Kuhlow, D., Weimer, S., Priebe, S., Heiland, I., Birringer, M., Groth, M., Segref, A., Kanfi, Y., et al. (2013). Role of sirtuins in lifespan regulation is linked to methylation of nicotinamide. Nature Chemical Biology 9, 693–700. 10.1038/nchembio.1352.

67. Schmeisser, K., and Parker, J.A. (2018). Nicotinamide-N-methyltransferase controls behavior, neurodegeneration and lifespan by regulating neuronal autophagy. PLoS Genet 14, e1007561. 10.1371/journal.pgen.1007561.

68. Mutlu, A.S., Duffy, J., and Wang, M.C. (2021). Lipid metabolism and lipid signals in aging and longevity. Developmental Cell, 1–14. 10.1016/j.devcel.2021.03.034.

69. Engel, K.M., Schiller, J., Galuska, C.E., and Fuchs, B. (2021). Phospholipases and Reactive Oxygen Species Derived Lipid Biomarkers in Healthy and Diseased Humans and Animals – A Focus on Lysophosphatidylcholine. Front. Physiol. 12, 732319. 10.3389/fphys.2021.732319.

70. Lucanic, M., Held, J.M., Vantipalli, M.C., Klang, I.M., Graham, J.B., Gibson, B.W., Lithgow, G.J., and Gill, M.S. (2011). N-acylethanolamine signalling mediates the effect of diet on lifespan in Caenorhabditis elegans. Nature 473, 226–229. 10.1038/nature10007.

71. Harrison, N., Lone, M.A., Kaul, T.K., Rodrigues, P.R., Ogungbe, I.V., and Gill, M.S. (2014). Characterization of N-Acyl Phosphatidylethanolamine-Specific Phospholipase-D Isoforms in the Nematode Caenorhabditis elegans. Plos One 9, e113007. 10.1371/journal.pone.0113007.

72. Bazopoulou, D., Knoefler, D., Zheng, Y., Ulrich, K., Oleson, B.J., Xie, L., Kim, M., Kaufmann, A., Lee, Y.-T., Dou, Y., et al. (2019). Developmental ROS individualizes organismal stress resistance and lifespan. Nature 576, 301–305. 10.1038/s41586-019-1814-y.

73. Oleson, B.J., Bazopoulou, D., and Jakob, U. (2021). Shaping longevity early in life: developmental ROS and H3K4me3 set the clock. Cell Cycle 20, 2337–2347. 10.1080/15384101.2021.1986317.

74. Johnson, A.A., and Stolzing, A. (2019). The role of lipid metabolism in aging, lifespan regulation, and age-related disease. Aging Cell 18, 16335–26. 10.1111/acel.13048.

75. Ratnappan, R., Amrit, F.R.G., Chen, S.-W., Gill, H., Holden, K., Ward, J., Yamamoto, K.R., Olsen, C.P., and Ghazi, A. (2014). Germline Signals Deploy NHR-49 to Modulate Fatty-Acid β-Oxidation and Desaturation in Somatic Tissues of C. elegans. Plos Genet 10, e1004829. 10.1371/journal.pgen.1004829.

76. Goh, G.Y.S., Winter, J.J., Bhanshali, F., Doering, K.R.S., Lai, R., Lee, K., Veal, E.A., and Taubert, S. (2018). NHR-49/HNF4 integrates regulation of fatty acid metabolism with a protective transcriptional response to oxidative stress and fasting. Aging Cell 13, e12743–14. 10.1111/acel.12743.

77. Gilst, M.R.V., Hadjivassiliou, H., Jolly, A., and Yamamoto, K.R. (2005). Nuclear Hormone Receptor NHR-49 Controls Fat Consumption and Fatty Acid Composition in C. elegans. Plos Biol 3, e53. 10.1371/journal.pbio.0030053.

78. Naim, N., Amrit, F.R.G., Ratnappan, R., DelBuono, N., Loose, J.A., and Ghazi, A. (2021). Cell nonautonomous roles of NHR-49 in promoting longevity and innate immunity. Aging Cell, e13413. 10.1111/acel.13413.

79. Han, H.-F., Nien, S.-F., Jiang, H.-S., Wu, J.-C., Chiang, C.-Y., Li, M.-T., Huang, L.-J., Chiang, S., Lin, L.-C., Chuang, Y.-T., et al. (2024). Dietary bacteria control C. elegans fat content through pathways converging at phosphatidylcholine. 10.7554/elife.96473.

80. Qin, S., Wang, Y., Li, L., Liu, J., Xiao, C., Duan, D., Hao, W., Qin, C., Chen, J., Yao, L., et al. (2022). Early-life vitamin B12 orchestrates lipid peroxidation to ensure reproductive success via SBP-1/SREBP1 in Caenorhabditis elegans. Cell Reports 40, 111381. 10.1016/j.celrep.2022.111381.

81. Walker, A.K., Jacobs, R.L., Watts, J.L., Rottiers, V., Jiang, K., Finnegan, D.M., Shioda, T., Hansen, M., Yang, F., Niebergall, L.J., et al. (2011). A Conserved SREBP- 1/Phosphatidylcholine Feedback Circuit Regulates Lipogenesis in Metazoans. Cell 147, 840– 852. 10.1016/j.cell.2011.09.045.

82. Lee, D., An, S.W.A., Jung, Y., Yamaoka, Y., Ryu, Y., Goh, G.Y.S., Beigi, A., Yang, J.- S., Jung, G.Y., Ma, D.K., et al. (2019). MDT-15/MED15 permits longevity at low temperature via enhancing lipidostasis and proteostasis. PLoS Biol. 17, e3000415. 10.1371/journal.pbio.3000415.

83. Shomer, N., Kadhim, A.Z., Grants, J.M., Cheng, X., Alhusari, D., Bhanshali, F., Poon, A.F.-Y., Lee, M.Y.Y., Muhuri, A., Park, J.I., et al. (2019). Mediator subunit MDT- 15/MED15 and Nuclear Receptor HIZR-1/HNF4 cooperate to regulate toxic metal stress responses in Caenorhabditis elegans. PLoS Genet. 15, e1008508. 10.1371/journal.pgen.1008508.

84. Soukas, A.A., Kane, E.A., Carr, C.E., Melo, J.A., and Ruvkun, G. (2009). Rictor/TORC2 regulates fat metabolism, feeding, growth, and life span in Caenorhabditis elegans. Genes & Development 23, 496–511. 10.1101/gad.1775409.

85. Aspernig, H., Heimbucher, T., Qi, W., Gangurde, D., Curic, S., Yan, Y., Gromoff, E.D. von, Baumeister, R., and Thien, A. (2019). Mitochondrial Perturbations Couple mTORC2 to Autophagy in C. elegans. Cell Reports 29, 1399–1409.e5. 10.1016/j.celrep.2019.09.072.

86. Blackwell, T.K., Sewell, A.K., Wu, Z., and Han, M. (2019). TOR Signaling in Caenorhabditis elegans Development, Metabolism, and Aging. Genetics 213, 329–360. 10.1534/genetics.119.302504.

87. Ruf, V., Holzem, C., Peyman, T., Walz, G., Blackwell, T.K., and Neumann-Haefelin, E. (2013). TORC2 signaling antagonizes SKN-1 to induce C. elegans mesendodermal embryonic development. Developmental Biology 384, 214–227. 10.1016/j.ydbio.2013.08.011.

88. Sheaffer, K.L., Updike, D.L., and Mango, S.E. (2008). The Target of Rapamycin Pathway Antagonizes pha-4/FoxA to Control Development and Aging. Curr. Biol. 18, 1355– 1364. 10.1016/j.cub.2008.07.097.

89. VanDerMolen, K.R., Newman, M.A., Breen, P.C., Huff, L.A., and Dowen, R.H. (2024). Non-cell-autonomous regulation of mTORC2 by Hedgehog signaling maintains lipid homeostasis. bioRxiv, 2024.05.06.592795. 10.1101/2024.05.06.592795.

90. Li, X., Yang, Y., Zhang, B., Lin, X., Fu, X., An, Y., Zou, Y., Wang, J.-X., Wang, Z., and Yu, T. (2022). Lactate metabolism in human health and disease. Signal Transduct. Target. Ther. 7, 305. 10.1038/s41392-022-01151-3.

91. Hayek, L.E., Khalifeh, M., Zibara, V., Assaad, R.A., Emmanuel, N., Karnib, N., El- Ghandour, R., Nasrallah, P., Bilen, M., Ibrahim, P., et al. (2019). Lactate Mediates the Effects of Exercise on Learning and Memory through SIRT1-Dependent Activation of Hippocampal Brain-Derived Neurotrophic Factor (BDNF). J Neurosci 39, 2369–2382. 10.1523/jneurosci.1661-18.2019.

92. Lian, B., Zhang, J., Yin, X., Wang, J., Li, L., Ju, Q., Wang, Y., Jiang, Y., Liu, X., Chen, Y., et al. (2024). SIRT1 improves lactate homeostasis in the brain to alleviate parkinsonism via deacetylation and inhibition of PKM2. Cell Rep. Med. 5, 101684. 10.1016/j.xcrm.2024.101684.

93. Choi, H.J., Jang, S.-Y., and Hwang, E.S. (2015). High-Dose Nicotinamide Suppresses ROS Generation and Augments Population Expansion during CD8+ T Cell Activation. Mol. Cells 38, 918–924. 10.14348/molcells.2015.0168.

94. Song, Z., Zhong, X., Li, M., Gao, P., Ning, Z., Sun, Z., and Song, X. (2021). 1-MNA Ameliorates High Fat Diet-Induced Heart Injury by Upregulating Nrf2 Expression and Inhibiting NF-κB in vivo and in vitro. Front. Cardiovasc. Med. 8, 721814. 10.3389/fcvm.2021.721814.

95. Barrett, L.N., and Westerheide, S.D. (2022). The CBP-1/p300 Lysine Acetyltransferase Regulates the Heat Shock Response in C. elegans. Frontiers Aging 3, 861761. 10.3389/fragi.2022.861761.

96. Babetto, E., Wong, K.M., and Beirowski, B. (2020). A glycolytic shift in Schwann cells supports injured axons. Nat. Neurosci. 23, 1215–1228. 10.1038/s41593-020-0689-4.

97. Morrison, B.M., Tsingalia, A., Vidensky, S., Lee, Y., Jin, L., Farah, M.H., Lengacher, S., Magistretti, P.J., Pellerin, L., and Rothstein, J.D. (2015). Deficiency in monocarboxylate transporter 1 (MCT1) in mice delays regeneration of peripheral nerves following sciatic nerve crush. Experimental Neurology 263, 325–338. 10.1016/j.expneurol.2014.10.018.

98. Jia, L., Liao, M., Mou, A., Zheng, Q., Yang, W., Yu, Z., Cui, Y., Xia, X., Qin, Y., Chen, M., et al. (2021). Rheb-regulated mitochondrial pyruvate metabolism of Schwann cells linked to axon stability. Developmental Cell, 1–22. 10.1016/j.devcel.2021.09.013.

99. San-Millan, I., Sparagna, G.C., Chapman, H.L., Warkins, V.L., Chatfield, K.C., Shuff, S.R., Martinez, J.L., and Brooks, G.A. (2022). Chronic Lactate Exposure Decreases Mitochondrial Function by Inhibition of Fatty Acid Uptake and Cardiolipin Alterations in Neonatal Rat Cardiomyocytes. Front. Nutr. 9, 809485. 10.3389/fnut.2022.809485.

100. Fang, Y., Li, Z., Yang, L., Li, W., Wang, Y., Kong, Z., Miao, J., Chen, Y., Bian, Y., and Zeng, L. (2024). Emerging roles of lactate in acute and chronic inflammation. Cell Commun. Signal. 22, 276. 10.1186/s12964-024-01624-8.

101. Turner, C.D., and Curran, S.P. (2025). Activated SKN-1 alters the aging trajectories of long-lived C. elegans mutants. GENETICS, iyaf016. 10.1093/genetics/iyaf016.

102. Bitto, A., Ito, T.K., Pineda, V.V., LeTexier, N.J., Huang, H.Z., Sutlief, E., Tung, H., Vizzini, N., Chen, B., Smith, K., et al. (2016). Transient rapamycin treatment can increase lifespan and healthspan in middle-aged mice. Elife 5, e16351. 10.7554/elife.16351.

103. Espada, L. (2020). Loss of metabolic plasticity underlies metformin toxicity in aged Caenorhabditis elegans. Nature Metabolism 21, 1–36. 10.1038/s42255-020-00307-1.

104. Oleson, B.J., Bhattrai, J., Zalubas, S.L., Kravchenko, T.R., Ji, Y., Jiang, E.L., Lu, C.C., Madden, C.R., Coffman, J.G., Bazopoulou, D., et al. (2023). Early life changes in histone landscape protect against age-associated amyloid toxicities through HSF-1-dependent regulation of lipid metabolism. Nat. Aging, 1–14. 10.1038/s43587-023-00537-4.

105. Torre, A. la, Vecchio, F.L., and Greco, A. (2023). Epigenetic Mechanisms of Aging and Aging-Associated Diseases. Cells 12, 1163. 10.3390/cells12081163.

106. Wang, W., Zheng, Y., Sun, S., Li, W., Song, M., Ji, Q., Wu, Z., Liu, Z., Fan, Y., Liu, F., et al. (2021). A genome-wide CRISPR-based screen identifies KAT7 as a driver of cellular senescence. Sci. Transl. Med. 13. 10.1126/scitranslmed.abd2655.

107. Zhao, S., Xu, W., Jiang, W., Yu, W., Lin, Y., Zhang, T., Yao, J., Zhou, L., Zeng, Y., Li, H., et al. (2010). Regulation of Cellular Metabolism by Protein Lysine Acetylation. Science 327, 1000–1004. 10.1126/science.1179689.

108. Guan, K.-L., and Xiong, Y. (2011). Regulation of intermediary metabolism by protein acetylation. Trends Biochem. Sci. 36, 108–116. 10.1016/j.tibs.2010.09.003.

109. Chavez-Guevara, I.A., Fermandez-Escabias, M., Hernandez-Lepe, M.A., and Amaro- Gahete, F.J. (2025). Modulation of Fatty Acid Metabolism via Lactate-HCA1 Signaling: Potential Therapeutic Implications. Am. J. Physiol.-Cell Physiol. 10.1152/ajpcell.00969.2024.

110. Lee, W.D., Weilandt, D.R., Liang, L., MacArthur, M.R., Jaiswal, N., Ong, O., Mann, C.G., Chu, Q., Hunter, C.J., Ryseck, R.-P., et al. (2025). Lactate homeostasis is maintained through regulation of glycolysis and lipolysis. Cell Metab. 10.1016/j.cmet.2024.12.009.

