## Supplementary material for "Lactate promotes longevity through redox-driven lipid remodeling in *Caenorhabditis elegans*": Supp. Figures and legends

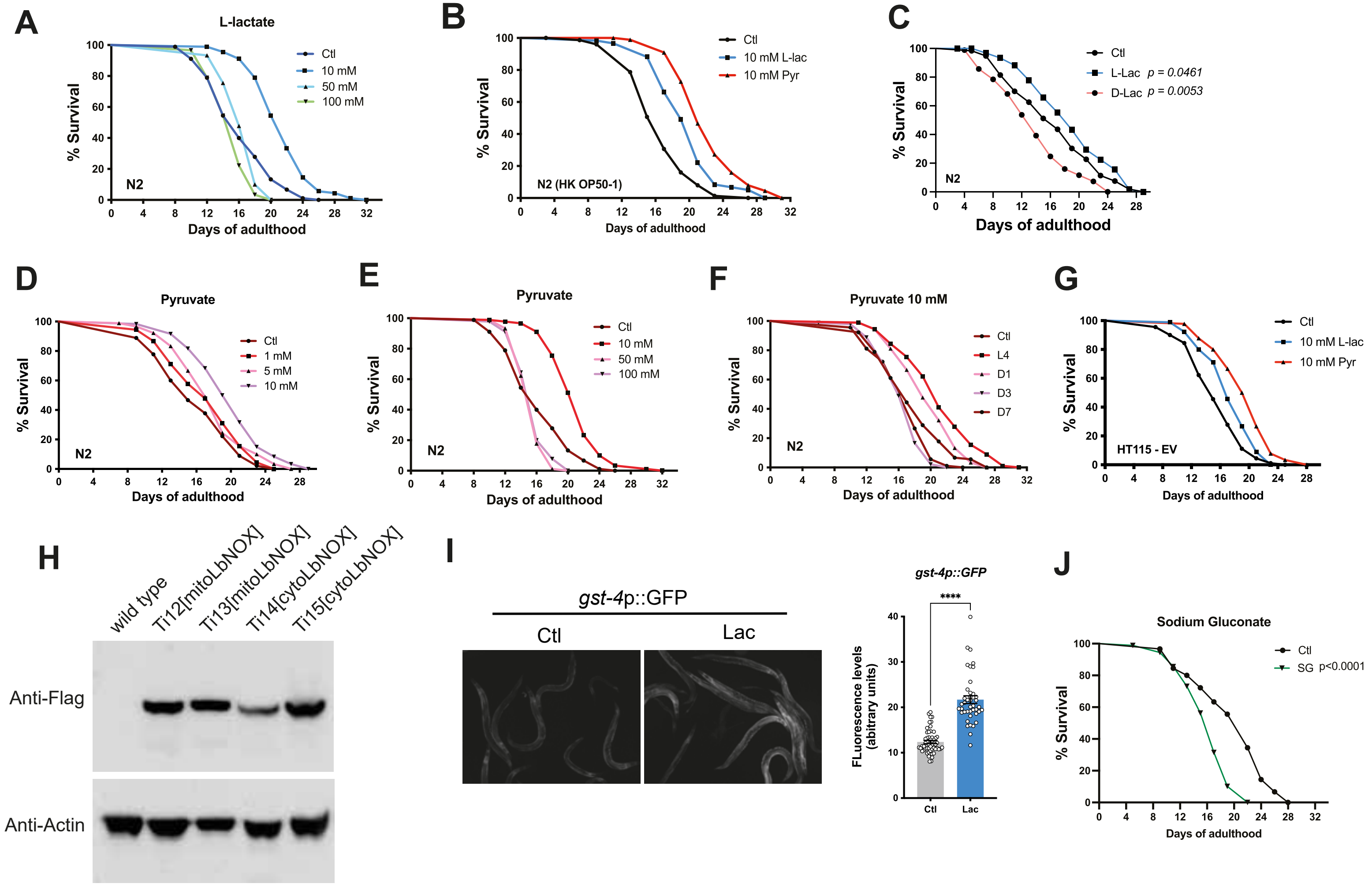
A
L-lactate
Ctl
10 mM
50 mM
100 mM
N2
% Survival
Days of adulthood
B
Ctl
10 mM L-lac
10 mM Pyr
N2 (HK OP50-1)
% Survival
Days of adulthood
C
Ctl
L-Lac p = 0.0461
D-Lac p = 0.0053
N2
% Survival
Days of adulthood
D
Pyruvate
Ctl
1 mM
5 mM
10 mM
N2
% Survival
Days of adulthood
E
Pyruvate
Ctl
10 mM
50 mM
100 mM
N2
% Survival
Days of adulthood
F
Pyruvate 10 mM
Ctl
L4
D1
D3
D7
N2
% Survival
Days of adulthood
G
Ctl
10 mM L-lac
10 mM Pyr
HT115 - EV
% Survival
Days of adulthood
H
wild type
Ti12[mitoLbNOX]
Ti13[mitoLbNOX]
Ti14[cytoLbNOX]
Ti15[cytoLbNOX]
Anti-Flag
Anti-Actin
I
gst-4p::GFP
Ctl
Lac
gst-4p::GFP
****
FLuorescence levels (abitrary units)
Ctl
Lac
J
Sodium Gluconate
Ctl
SG p<0.0001
% Survival
Days of adulthood

FIG. S1

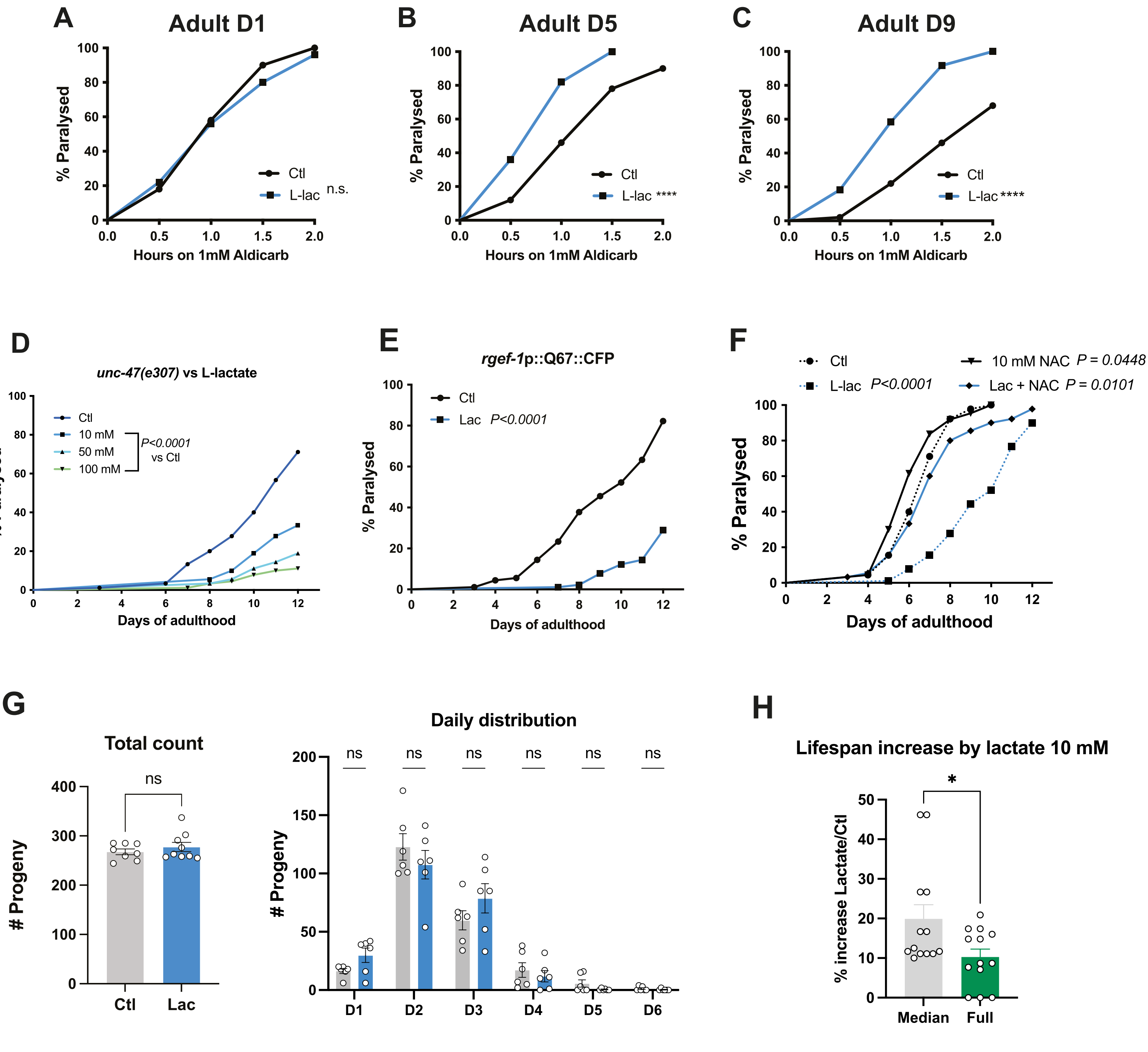
A
Adult D1
% Paralysed
Hours on 1mM Aldicarb
Ctl
L-lac n.s.
B
Adult D5
Ctl
L-lac ****
C
Adult D9
Ctl
L-lac ****
D
unc-47(e307) vs L-lactate
Ctl
10 mM
50 mM
100 mM
P<0.0001 vs Ctl
Days of adulthood
E
rgef-1p::Q67::CFP
Ctl
Lac P<0.0001
F
Ctl
L-lac P<0.0001
10 mM NAC P = 0.0448
Lac + NAC P = 0.0101
G
Total count
ns
### Progeny
Ctl
Lac
Daily distribution
D1
D2
D3
D4
D5
D6
H
Lifespan increase by lactate 10 mM
% increase Lactate/Ctl
Median
Full

FIG. S2

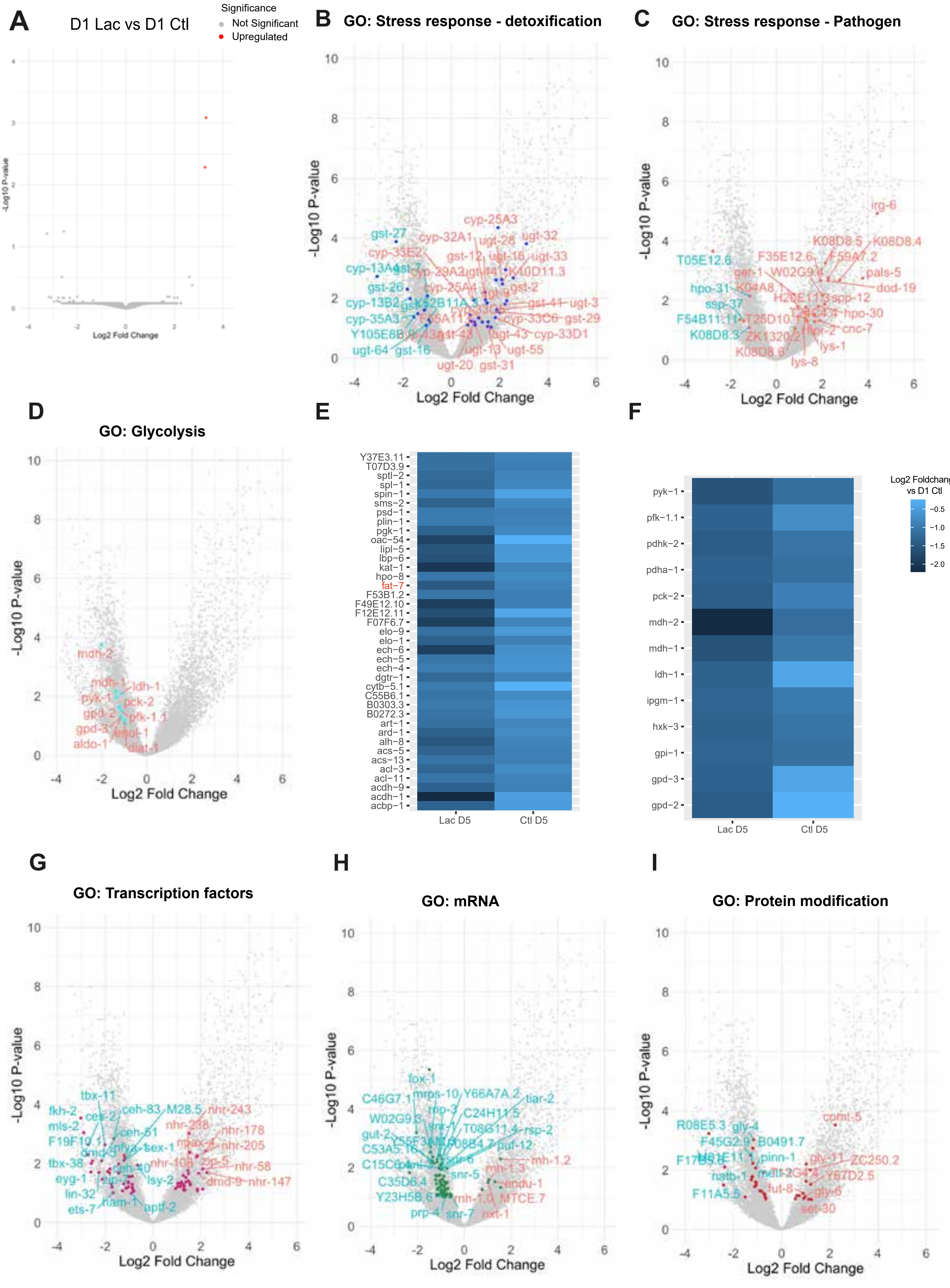
A
D1 Lac vs D1 Ctl
Significance
Not Significant
Upregulated
-Log10 P-value
Log2 Fold Change
B
GO: Stress response - detoxification
C
GO: Stress response - Pathogen
D
GO: Glycolysis
E
Y37E3.11
T07D3.9
sptl−2
spl−1
spin−1
sms−2
psd−1
plin−1
pgk−1
oac−54
lipl−5
lbp−6
kat−1
hpo−8
fat−7
F53B1.2
F49E12.10
F12E12.11
F07F6.7
elo−9
elo−1
ech−6
ech−5
ech−4
dgtr−1
cytb−5.1
C55B6.1
B0303.3
B0272.3
art−1
ard−1
alh−8
acs−5
acs−13
acl−3
acl−11
acdh−9
acdh−1
acbp−1
Lac D5
Ctl D5
F
pyk−1
pfk−1.1
pdhk−2
pdha−1
pck−2
mdh−2
mdh−1
ldh−1
ipgm−1
hxk−3
gpi−1
gpd−3
gpd−2
Lac D5
Ctl D5
Log2 Foldchange vs D1 Ctl
−0.5
−1.0
−1.5
−2.0
G
GO: Transcription factors
H
GO: mRNA
I
GO: Protein modification
-Log10 P-value
Log2 Fold Change

Fig. S3

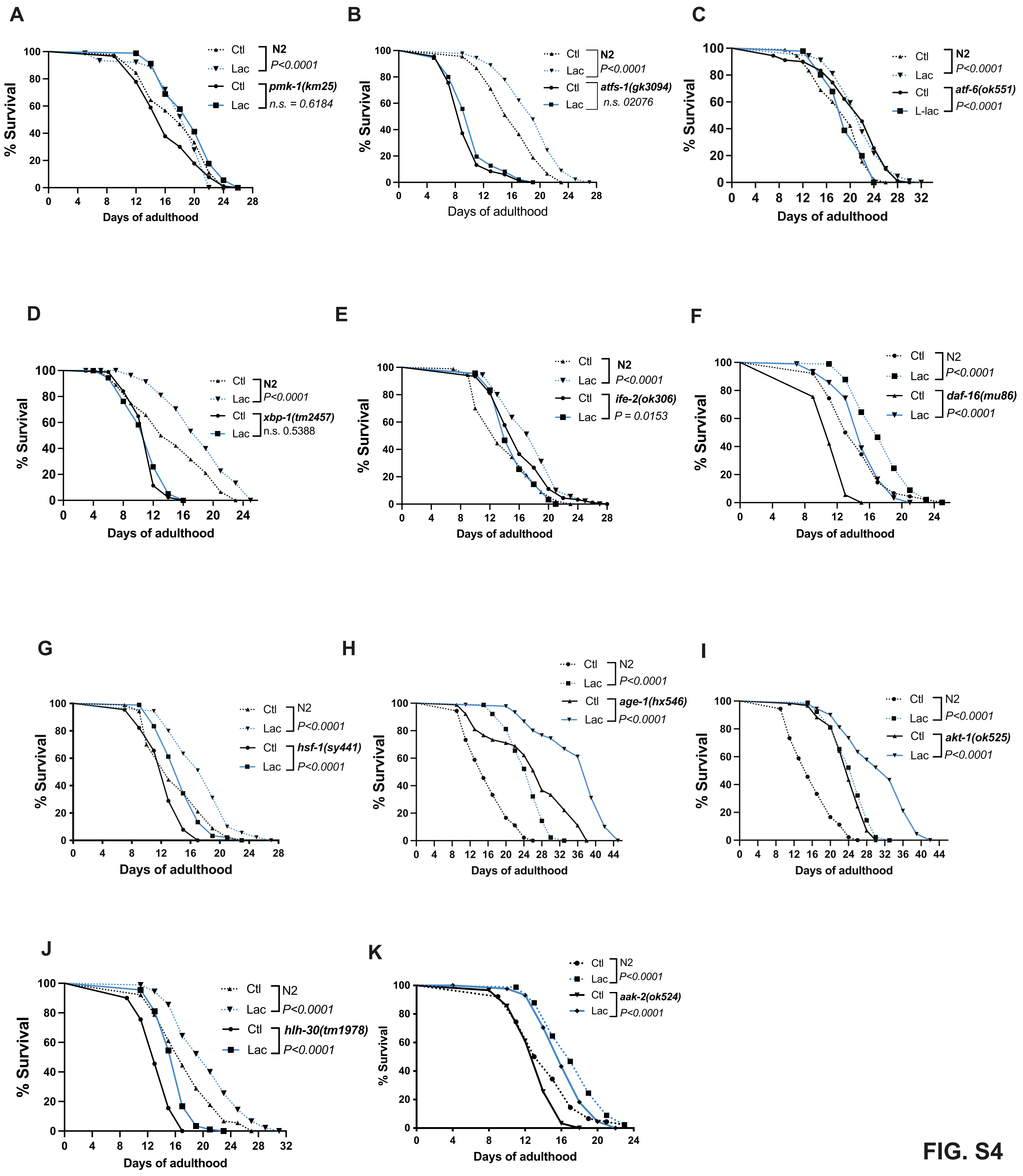

A
Ctl
Lac
N2
P<0.0001
Ctl
Lac
pmk-1(km25)
n.s. = 0.6184
% Survival
Days of adulthood
B
Ctl
Lac
N2
P<0.0001
Ctl
Lac
atfs-1(gk3094)
n.s. 02076
% Survival
Days of adulthood
C
Ctl
Lac
N2
P<0.0001
Ctl
L-lac
atf-6(ok551)
P<0.0001
% Survival
Days of adulthood
D
Ctl
Lac
N2
P<0.0001
Ctl
Lac
xbp-1(tm2457)
n.s. 0.5388
% Survival
Days of adulthood
E
Ctl
Lac
N2
P<0.0001
Ctl
Lac
ife-2(ok306)
P = 0.0153
% Survival
Days of adulthood
F
Ctl
Lac
N2
P<0.0001
Ctl
Lac
daf-16(mu86)
P<0.0001
% Survival
Days of adulthood
G
Ctl
Lac
N2
P<0.0001
Ctl
Lac
hsf-1(sy441)
P<0.0001
% Survival
Days of adulthood
H
Ctl
Lac
N2
P<0.0001
Ctl
Lac
age-1(hx546)
P<0.0001
% Survival
Days of adulthood
I
Ctl
Lac
N2
P<0.0001
Ctl
Lac
akt-1(ok525)
P<0.0001
% Survival
Days of adulthood
J
Ctl
Lac
N2
P<0.0001
Ctl
Lac
hlh-30(tm1978)
P<0.0001
% Survival
Days of adulthood
K
Ctl
Lac
N2
P<0.0001
Ctl
Lac
aak-2(ok524)
P<0.0001
% Survival
Days of adulthood


**FIG. S4**

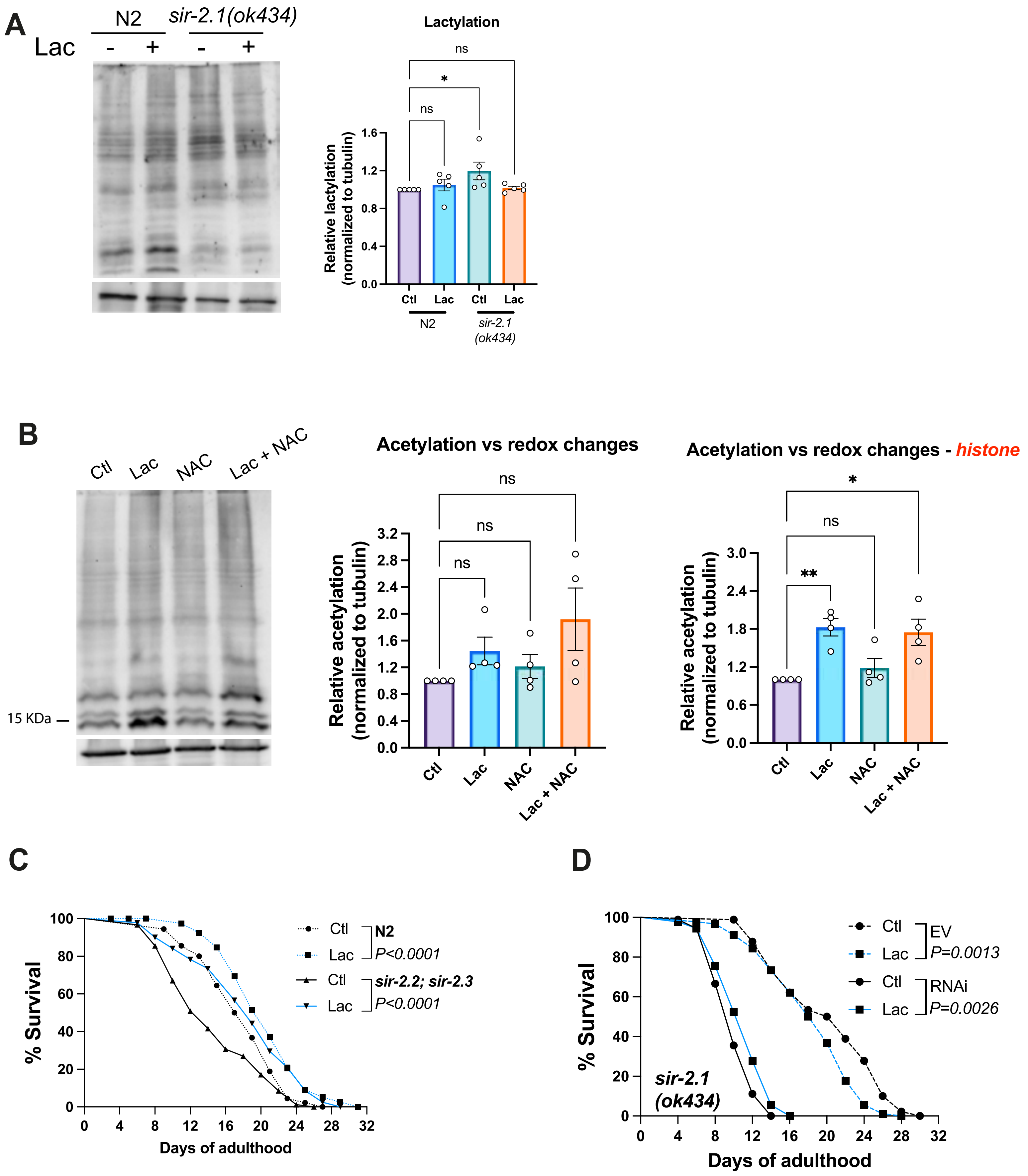
A
Lac
N2
sir-2.1(ok434)
Lactylation
ns
*
ns
Relative lactylation (normalized to tubulin)
Ctl
Lac
Ctl
Lac
N2
sir-2.1 (ok434)
B
Ctl
Lac
NAC
Lac + NAC
15 KDa
Acetylation vs redox changes
Relative acetylation (normalized to tubulin)
Acetylation vs redox changes - histone
**
C
% Survival
Days of adulthood
Ctl
Lac
N2
P<0.0001
sir-2.2; sir-2.3
P<0.0001
D
EV
P=0.0013
RNAi
P=0.0026
sir-2.1 (ok434)

Fig. S5

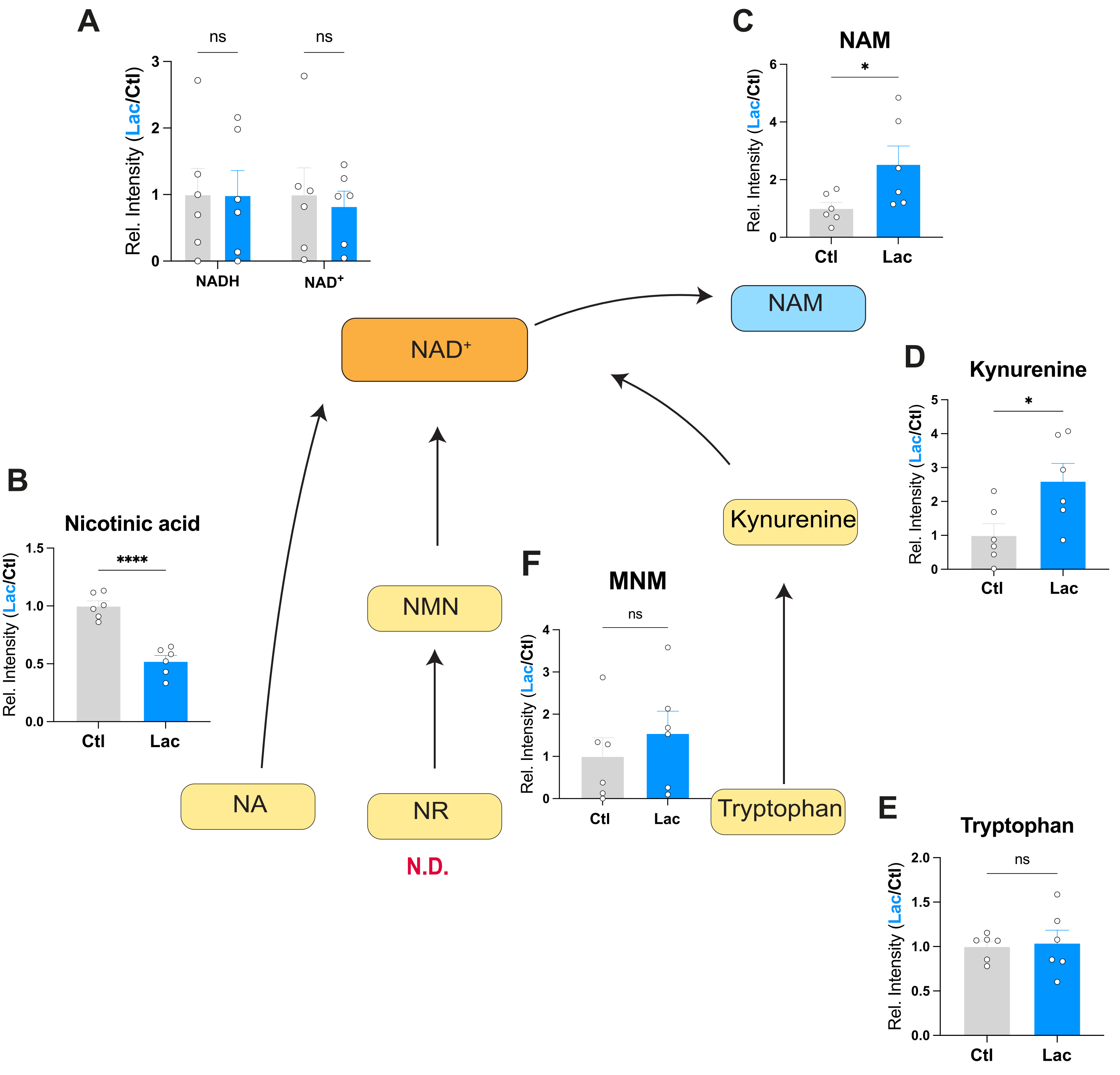
A
ns
ns
Rel. Intensity (Lac/Ctl)
NADH
NAD+
C
NAM
*
Rel. Intensity (Lac/Ctl)
Ctl
Lac
NAM
NAD+
D
Kynurenine
*
Rel. Intensity (Lac/Ctl)
Ctl
Lac
B
Nicotinic acid
****
Rel. Intensity (Lac/Ctl)
Ctl
Lac
Kynurenine
NMN
F
MNM
ns
Rel. Intensity (Lac/Ctl)
Ctl
Lac
NA
NR
N.D.
Tryptophan
E
Tryptophan
ns
Rel. Intensity (Lac/Ctl)
Ctl
Lac

Fig. S6

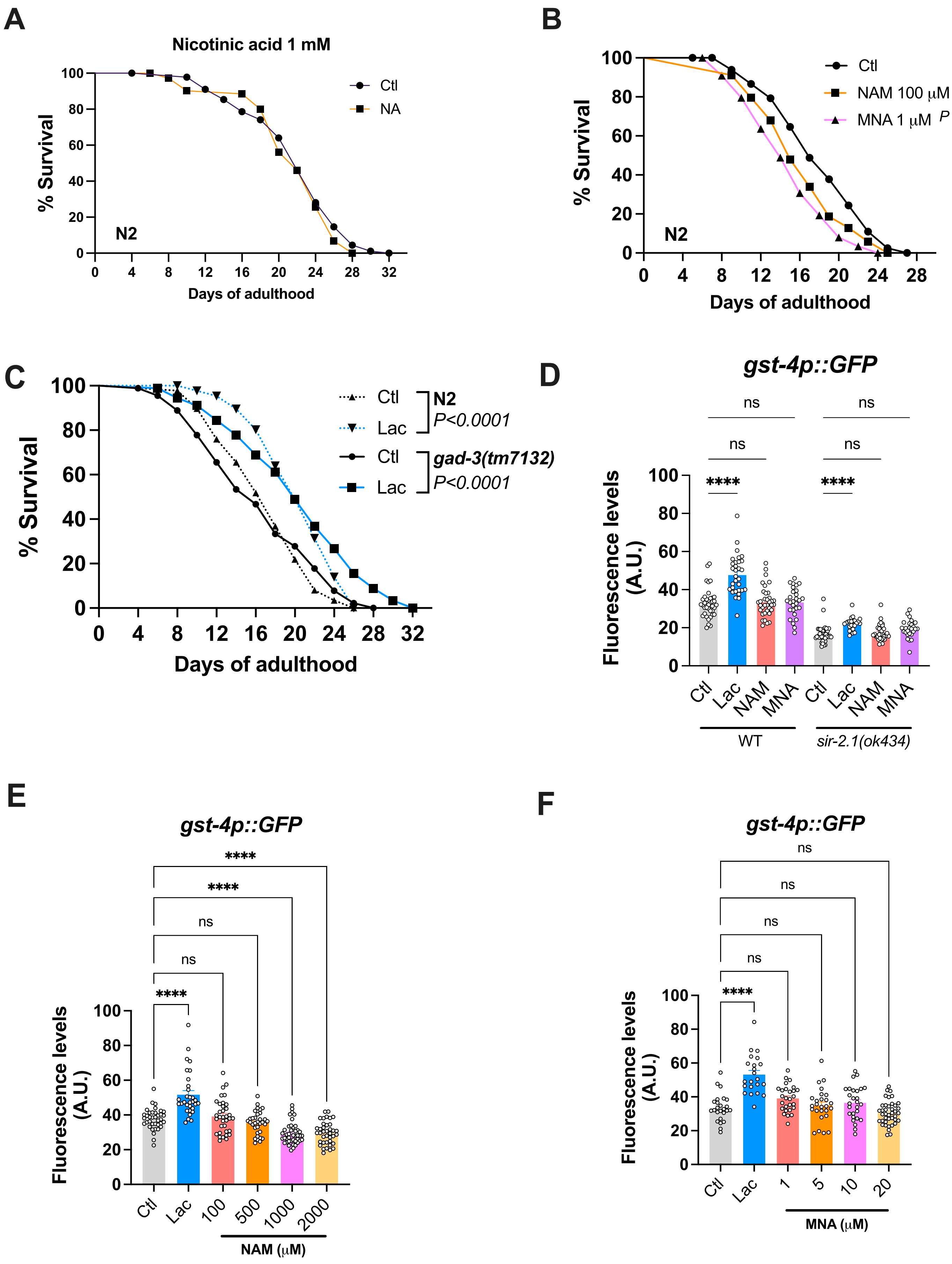
A
Nicotinic acid 1 mM
% Survival
Days of adulthood
Ctl
NA
N2
B
% Survival
Days of adulthood
Ctl
NAM 100 μM P = 0.0107
MNA 1 μM P < 0.0001
N2
C
% Survival
Days of adulthood
Ctl
Lac
N2
P<0.0001
Ctl
Lac
gad-3(tm7132)
P<0.0001
D
gst-4p::GFP
Fluorescence levels (A.U.)
ns
ns
ns
ns
****
****
Ctl
Lac
NAM
MNA
Ctl
Lac
NAM
MNA
WT
sir-2.1(ok434)
E
gst-4p::GFP
Fluorescence levels (A.U.)
****
****
ns
ns
****
Ctl
Lac
100
500
1000
2000
NAM (μM)
F
gst-4p::GFP
Fluorescence levels (A.U.)
ns
ns
ns
ns
****
Ctl
Lac
1
5
10
20
MNA (μM)

Fig. S7

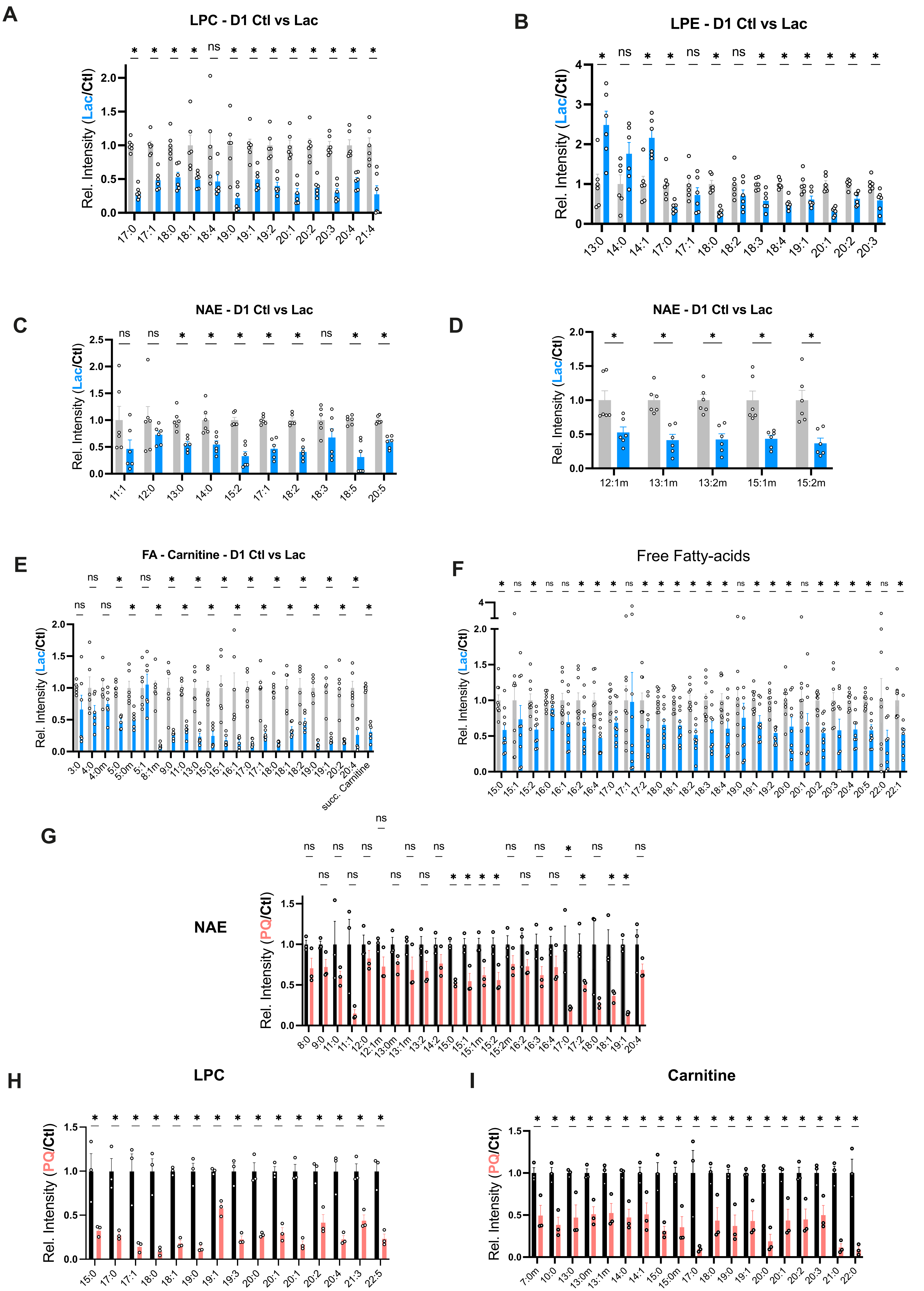
A
LPC - D1 Ctl vs Lac
Rel. Intensity (Lac/Ctl)
17:0 17:1 18:0 18:1 18:4 19:0 19:1 19:2 20:1 20:2 20:3 20:4 21:4
B
LPE - D1 Ctl vs Lac
Rel. Intensity (Lac/Ctl)
13:0 14:0 14:1 17:0 17:1 18:0 18:2 18:3 18:4 19:1 20:1 20:2 20:3
C
NAE - D1 Ctl vs Lac
Rel. Intensity (Lac/Ctl)
11:1 12:0 13:0 14:0 15:2 17:1 18:2 18:3 18:5 20:5
D
NAE - D1 Ctl vs Lac
Rel. Intensity (Lac/Ctl)
12:1m 13:1m 13:2m 15:1m 15:2m
E
FA - Carnitine - D1 Ctl vs Lac
Rel. Intensity (Lac/Ctl)
3:0 4:0 4:0m 5:0 5:0m 5:1 8:1m 9:0 11:0 13:0 15:0 15:1 16:1 17:0 17:1 18:0 18:1 18:2 19:0 19:1 20:2 20:4 succ. Carnitine
F
Free Fatty-acids
Rel. Intensity (Lac/Ctl)
15:0 15:1 15:2 16:0 16:1 16:2 16:4 17:0 17:1 17:2 18:0 18:1 18:2 18:3 18:4 19:0 19:1 19:2 20:0 20:1 20:2 20:3 20:4 20:5 22:0 22:1
G
NAE
Rel. Intensity (PQ/Ctl)
8:0 9:0 11:0 11:1 12:0 12:1m 13:0m 13:1m 13:2 14:2 15:0 15:1 15:1m 15:2 15:2m 16:2 16:3 16:4 17:0 17:2 18:0 18:1 19:1 20:4
H
LPC
Rel. Intensity (PQ/Ctl)
15:0 17:0 17:1 18:0 18:1 19:0 19:1 19:3 20:0 20:1 20:1 20:2 20:4 21:3 22:5
I
Carnitine
Rel. Intensity (PQ/Ctl)
7:0m 10:0 13:0 13:0m 13:1m 14:0 14:1 15:0 15:0m 17:0 18:0 19:0 19:1 20:0 20:1 20:2 20:3 21:0 22:0

Fig. S8

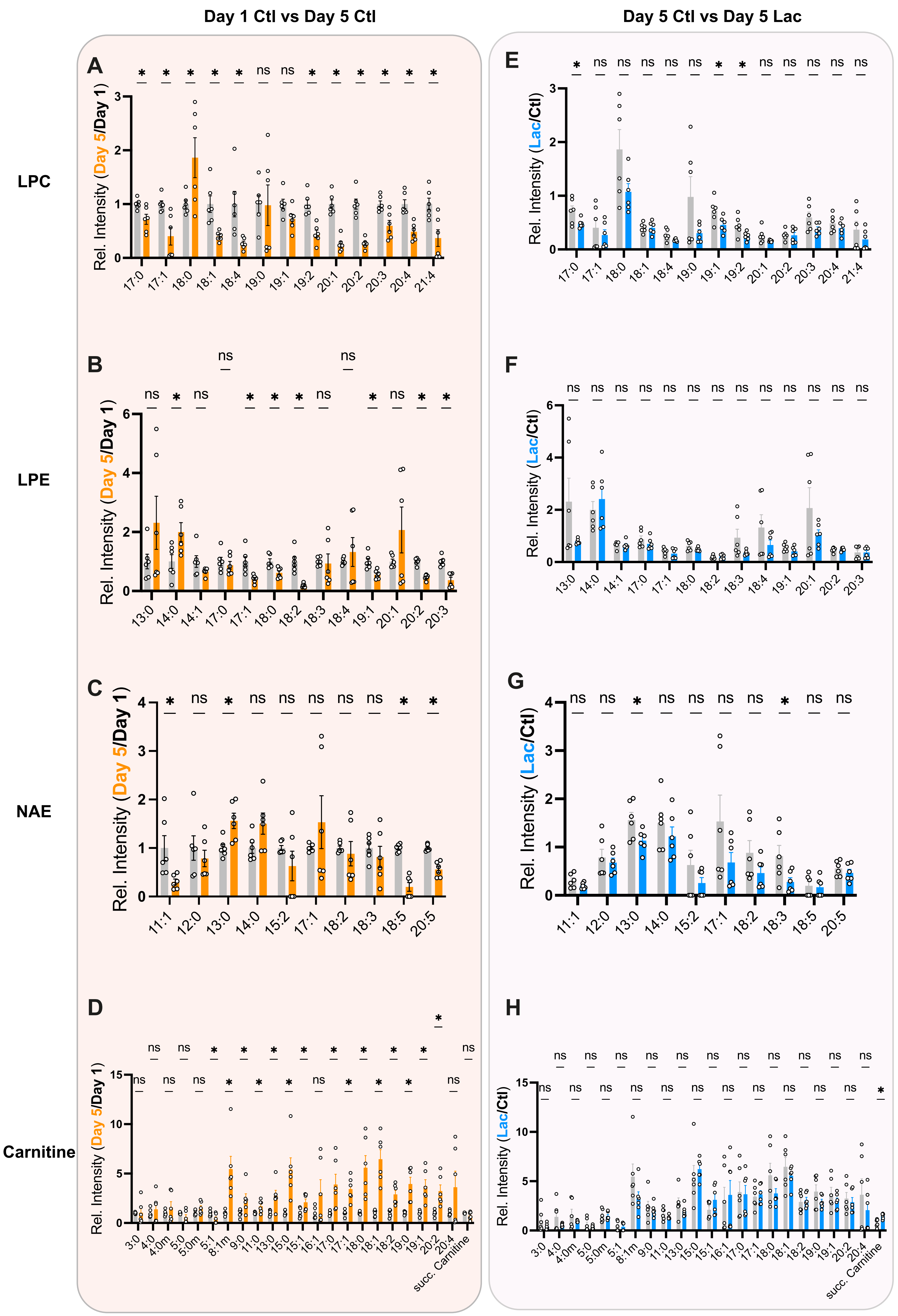

Day 1 Ctl vs Day 5 Ctl
Day 5 Ctl vs Day 5 Lac
LPC
LPE
NAE
Carnitine
A
Rel. Intensity (Day 5/Day 1)
17:0 17:1 18:0 18:1 18:4 19:0 19:1 19:2 20:1 20:2 20:3 20:4 21:4
* * * * * ns ns * * * * * *
B
Rel. Intensity (Day 5/Day 1)
13:0 14:0 14:1 17:0 17:1 18:0 18:2 18:3 18:4 19:1 20:1 20:2 20:3
ns * ns ns * * * ns ns * ns * *
C
Rel. Intensity (Day 5/Day 1)
11:1 12:0 13:0 14:0 15:2 17:1 18:2 18:3 18:5 20:5
* ns * ns ns ns ns ns * *
D
Rel. Intensity (Day 5/Day 1)
3:0 4:0 4:0m 5:0 5:0m 5:1 8:1m 9:0 11:0 13:0 15:0 15:1 16:1 17:0 17:1 18:0 18:1 18:2 19:0 19:1 20:2 20:4 succ. Carnitine
ns ns * * * * * * * * ns
ns ns ns * * * ns * * * ns
*
E
Rel. Intensity (Lac/Ctl)
17:0 17:1 18:0 18:1 18:4 19:0 19:1 19:2 20:1 20:2 20:3 20:4 21:4
* ns ns ns ns ns * * ns ns ns ns ns
F
Rel. Intensity (Lac/Ctl)
13:0 14:0 14:1 17:0 17:1 18:0 18:2 18:3 18:4 19:1 20:1 20:2 20:3
ns ns ns ns ns ns ns ns ns ns ns ns ns
G
Rel. Intensity (Lac/Ctl)
11:1 12:0 13:0 14:0 15:2 17:1 18:2 18:3 18:5 20:5
ns ns * ns ns ns ns * ns ns
H
Rel. Intensity (Lac/Ctl)
3:0 4:0 4:0m 5:0 5:0m 5:1 8:1m 9:0 11:0 13:0 15:0 15:1 16:1 17:0 17:1 18:0 18:1 18:2 19:0 19:1 20:2 20:4 succ. Carnitine
ns ns ns ns ns ns ns ns ns ns ns
ns ns ns ns ns ns ns ns ns ns ns *


Fig. S9

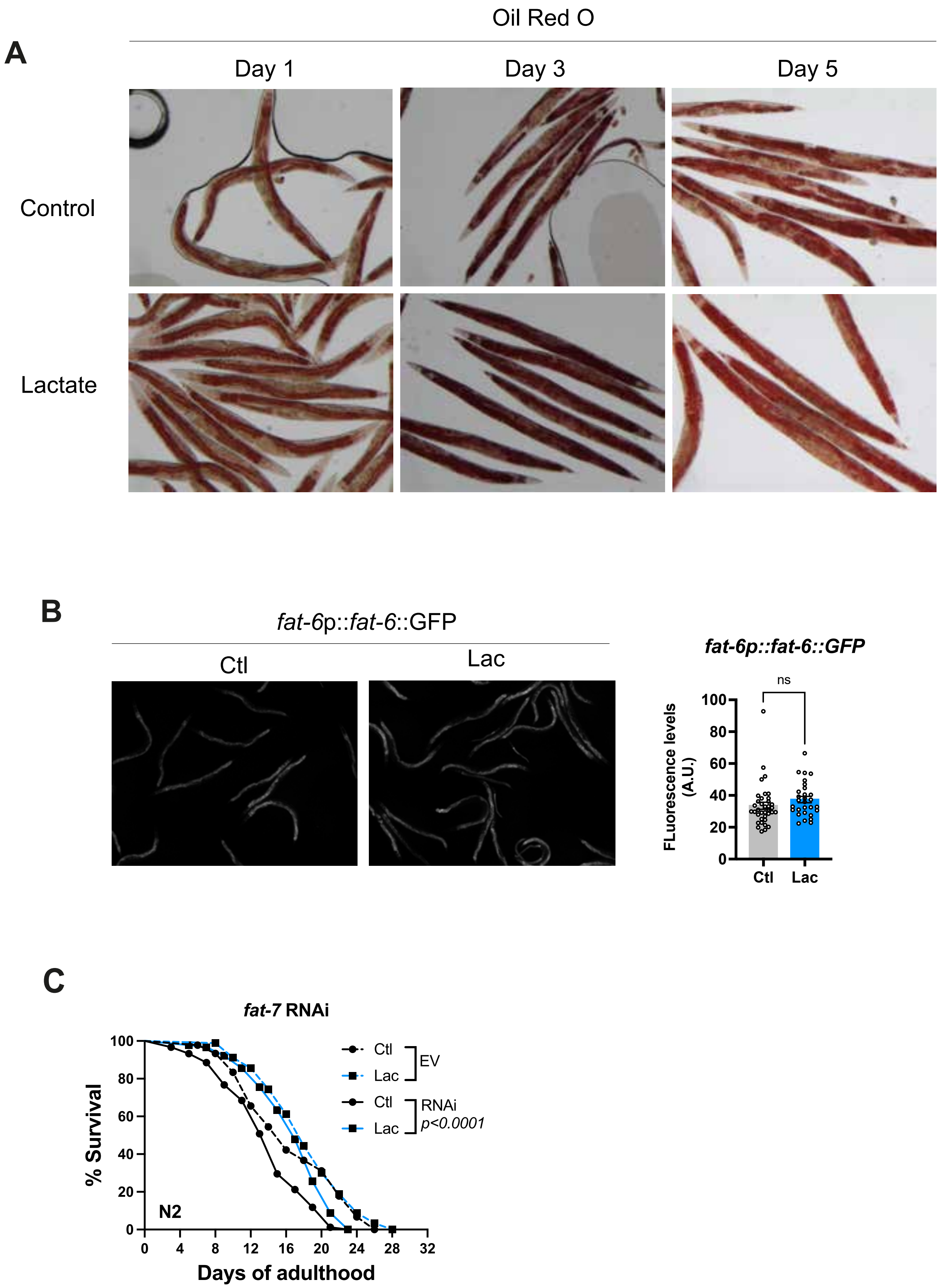
A
Oil Red O
Day 1
Day 3
Day 5
Control
Lactate
B
fat-6p::fat-6::GFP
Ctl
Lac
fat-6p::fat-6::GFP
ns
FLuorescence levels (A.U.)
100
80
60
40
20
0
Ctl
Lac
C
fat-7 RNAi
% Survival
Days of adulthood
N2
Ctl
Lac
EV
Ctl
Lac
RNAi
p<0.0001
0
4
8
12
16
20
24
28
32

Fig. S10

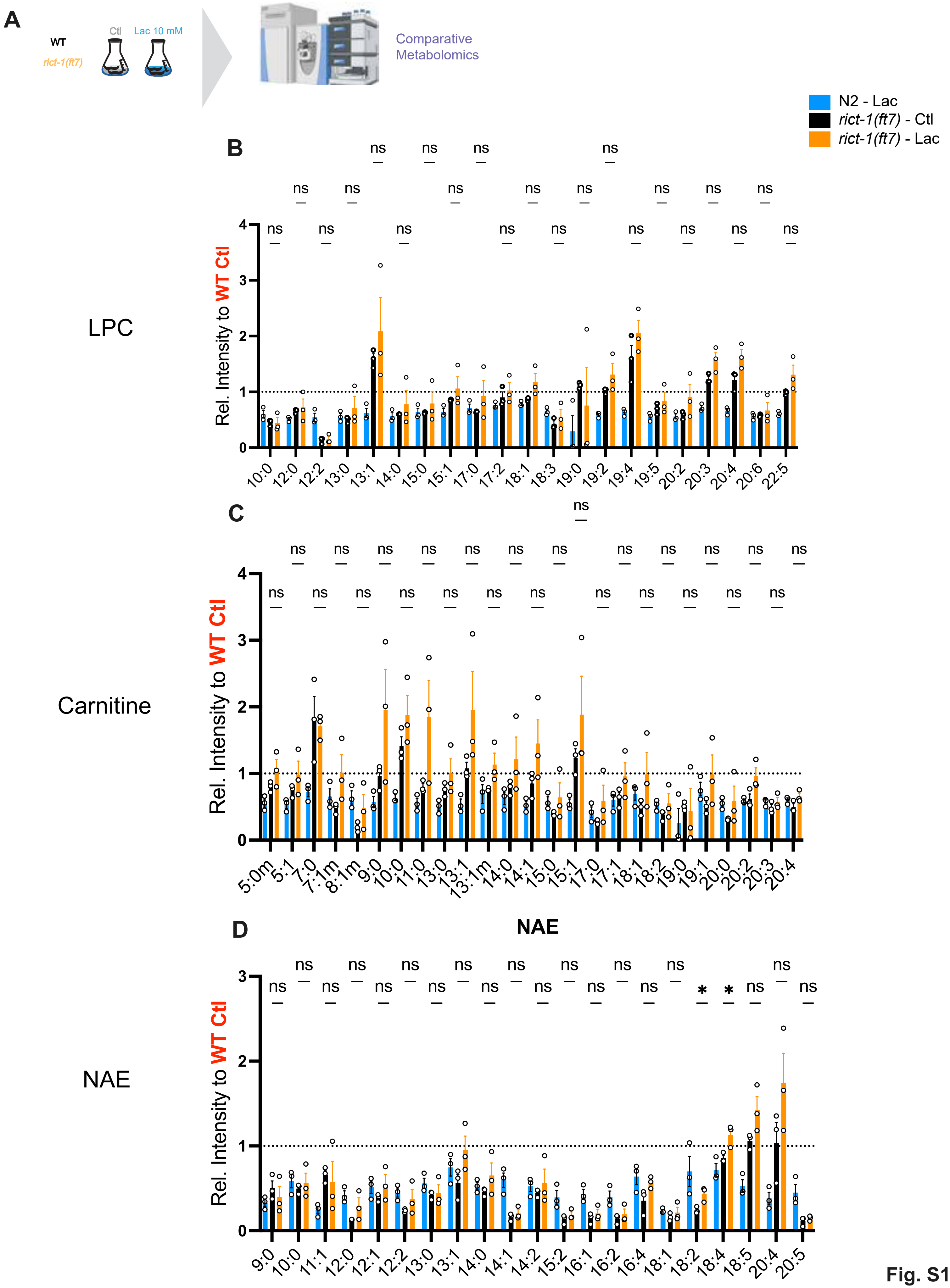
A
WT
rict-1(ft7)
Ctl
Lac 10 mM
Comparative Metabolomics
N2 - Lac
rict-1(ft7) - Ctl
rict-1(ft7) - Lac
B
LPC
Rel. Intensity to WT Ctl
10:0 12:0 12:2 13:0 13:1 14:0 15:0 15:1 17:0 17:2 18:1 18:3 19:0 19:2 19:4 19:5 20:2 20:3 20:4 20:6 22:5
C
Carnitine
Rel. Intensity to WT Ctl
5:0m 5:1 7:0 7:1m 8:1m 9:0 10:0 11:0 13:0 13:1 13:1m 14:0 14:1 15:0 15:1 17:0 17:1 18:1 18:2 19:0 19:1 20:0 20:2 20:3 20:4
D
NAE
NAE
Rel. Intensity to WT Ctl
9:0 10:0 11:1 12:0 12:1 12:2 13:0 13:1 14:0 14:1 14:2 15:2 16:1 16:2 16:4 18:1 18:2 18:4 18:5 20:4 20:5
ns
*

Fig. S11

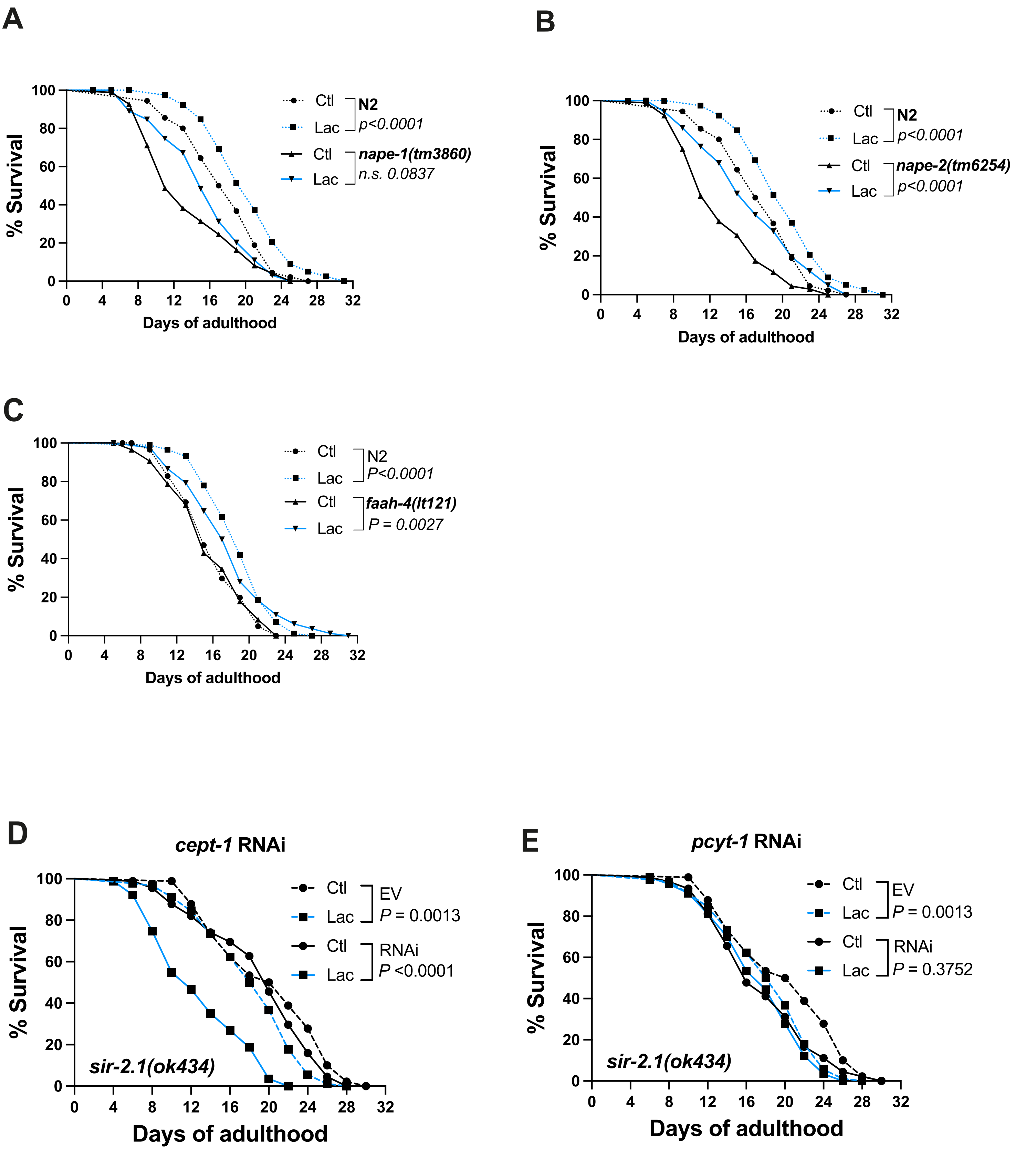
A
% Survival
Days of adulthood
Ctl
Lac
N2
p<0.0001
Ctl
Lac
nape-1(tm3860)
n.s. 0.0837
B
% Survival
Days of adulthood
Ctl
Lac
N2
p<0.0001
Ctl
Lac
nape-2(tm6254)
p<0.0001
C
% Survival
Days of adulthood
Ctl
Lac
N2
P<0.0001
Ctl
Lac
faah-4(lt121)
P = 0.0027
D
cept-1 RNAi
% Survival
Days of adulthood
Ctl
Lac
EV
P = 0.0013
Ctl
Lac
RNAi
P <0.0001
sir-2.1(ok434)
E
pcyt-1 RNAi
% Survival
Days of adulthood
Ctl
Lac
EV
P = 0.0013
Ctl
Lac
RNAi
P = 0.3752
sir-2.1(ok434)

**Fig. S12**

**Figures legend** :

**Fig S1: Characterization of longevity phenotype by lactate and pyruvate.** (*related to figure 1*)

**A)** Lifespan curves for N2 animals supplemented with different doses of lactate.

**B)** Longevity curves for N2 animals supplemented with either 10 mM lactate or pyruvate. Animals were grown on heat-killed bacteria (OP50-1)

**C)** Lifespan curves for N2 animals treated with 10 mM of with (**L**)-lactate or (**D**)-lactate.

**D-E)** Lifespan curves for N2 animals treated with 10 mM pyruvate. Treatments were started **at** indicated stages and continued until death.

**F)** Lifespan curves for N2 animals under control diet (black) or different temporal pyruvate supplementation. Full treatment was applied at embryonic stage until death. The life-stage in legend indicates when pyruvate treatment was applied.

**G)** Longevity curves for N2 animals supplemented with either 10 mM lactate or pyruvate. Animals were grown on L4440 RNAi

**H)** Immunoblot for JDM458(Ti12), JDM459(Ti13), JDM460(Ti14) and JDM460 (ti15). Expression of LbNOX in each strain was measured using anti-FLAG antibody. Wild-type stain (N2) was used as a negative control

**I)** Images of *gst-4*p::GFP reporter animals at day 1 adult stage, under control or lactate supplemented diet. Quantification *gst-4* reporter expression in day 1 animals under control or lactate-supplemented diet. Each dot represents one animal.

**J)** Lifespan curves for N2 animals treated with 10 mM sodium gluconate.

Statistics for lifespan curves were done using log-rank (Mantel–Cox) method. For each condition, 90 animals were analyzed (in triplicates) each experiment was repeated at least twice. P values for fluorescence were calculated by unpaired, two-sided t-test with Welch correction; ****P<0.0001. ns: Not significant. Each fluorescence experiment was performed at least 3 times and between 25-40 animals were measured per condition.

**Fig S2: Characterization of lactate healthspan impact.** *(related to figure 1).*

**A-C)** Paralysis curves for N2 animals treated with 1 mM aldicarb. Animals were grown on control or lactate-enriched diet and paralysis assessed at day 1, 5 and 9 of adulthood. 30 animals were scored per condition

**D)** Paralysis curves for *unc-47(e307)* supplemented with different doses of lactate.

**E-F)** Paralysis curves for *rgef-1p::Q67::CFP* animals supplemented with 10 mM L-lactate **(E)**. Paralysis rate of Q67::CFP grown on N-acetyl-L-cysteine (NAC 5 mM) was also measured together with lactate enrichment **(F)**.

**G)** Brood size of N2 animals grown on control or lactate-enriched diet. Number of viable progeny was measured over 6 days.

**H)** Quantification of median and full lifespan extension by 10 mM lactate in N2 animals. . Animals were grown on control or lactate (10 mM) enriched diets. Each measure represent an independent experiment and is represented as a percentage (%) of lactate-treated over control animals.

Statistics for paralysis curves were done using log-rank (Mantel–Cox) method. For each condition, 90 animals were analyzed (in triplicates) each experiment was repeated at least twice. ns: Not significant. Brood size was measured over 25-30 animals, over 4 independent experiments P values for median and full lifespan comparisons were calculated by paired, two-sided t-test with Welch correction

**Fig S3**: **Characterization of Lactate-mediated transcriptomic changes.** *(related to figure 3)*

**A)** Volcano plot for differentially expressed transcripts in D1 lactate vs D1 control. Statistical analysis uses Benjamini–Hochberg correction.

**B-C)** Volcano plot for differentially expressed transcripts in D5 lactate vs D1 control. Transcripts specifically regulated by D5 lactate for the gene ontology (GO) category stress response – detoxification **(B)**, stress response – pathogen **(C)** are highlighted in blue and orange, respectively.

**D)** Volcano plot for differentially expressed transcripts in D5 lactate vs D1 control. Transcripts specifically regulated by D5 lactate for the gene ontology (GO) category metabolism – glycolysis.

**E-F)** List of transcripts differentially in day 5 adults animals supplemented with 10 mM lactate. Pathway enrichment analysis was performed using *Wormcat*. Day 5 animals treated with lactate show decrease level of glycolysis **(D)** and lipid metabolism **(E)** related transcripts.

**G-I)** Volcano plot for differentially expressed transcripts in D5 lactate vs D1 control. Transcripts specifically regulated by D5 lactate for the gene ontology (GO) Transcription factors **(G)**, mRNA **(H)** and Protein modifications **(I)** are highlighted in pink, green and orange, respectively. Statistical analysis uses Benjamini–Hochberg correction. P-adjusted value cutoff was *padj<0.05*.

Differential gene expression (DEG) between D5 lactate and D1 control conditions was assessed using the unpaired, nonparametric Mann–Whitney test followed by Benjamini–Hochberg correction for multiple comparisons. Cutoff for significance was fold change ≥1.5 and P-adjusted value of ≤0.05. List of transcripts specifically regulated by lactate were generated by comparing DEG D5 Lactate vs D1 control vs D5 control vs D1 control. Gene ontology analysis for differentially regulated between day 5 ctl and day 5 lac animals was performed using *Wormcat*[43] . Enriched categories with an adjusted P-value ≤0.05 were considered significant.

**Fig S4**: **Targeted screen for metabolic signals under lactate diet** (*Related to figure 3*).

**A-I)** Lifespan curves for *pmk-1(km25)* **(A)**, *atfs-1(gk3094)* **(B)**, *atf-6(ok551)* **(C)**, *xbp-1(zc12)* **(D)**, ife-2(ok306) **(E)**, *daf-16(mu86)* **(F),** *hsf-1(sy441)* **(G),** *age-1(hx546)* **(H)**, *akt-1(ok525)* **(I)**, *hlh-30(tm1978)* **(J)** and *aak2(ok524)* **(K)** loss of function mutants. Animals were grown on control diet or supplemented with 10 mM lactate.

Statistics for paralysis curves were done using log-rank (Mantel–Cox) method. For each condition, 90 animals were analyzed (in triplicates) each experiment was repeated at least twice

**Fig S5: Characterization of protein modification by lactate** *(related to figure 4)***.**

**A)** Immunoblot for lysine lactylation. N2 and *sir-2.1(ok434)* were grown on control and lactate enriched diet (10 mM) and protein extracted at adult day 1 stage. Lactylation level were normalized to tubulin and represented relative to control condition.

**B)** Immunoblot for lysine acetylation. N2 animals were grown on control, lactate enriched diet (10 mM), N-acetylcysteine (5 mM) or NAC + lactate. Protein extracted at adult day 1 stage. Acetylation level were normalized to tubulin and represented relative to control condition.

**C)** Lifespan curves of *sir-2.2; 2.3* loss of function mutants. Animals were grown on control diet or supplemented with 10 mM lactate

**D)** Lifespan curves of *sir-2.1(ok434)* animals treated with *cbp-1* RNAi. Animals were grown on control or lactate-enriched diet, on control RNAi. Animals were transferred on *cbp-1* RNAi at L4 stage.

Statistics for lifespan curves were done using log-rank (Mantel–Cox) method. For each condition, 90 animals were analyzed (in triplicates) each experiment was repeated at least twice

**Fig S6: Metabolic profile of NAD-associated metabolites** *(related to figure 4)*.

**A-E)** Quantification several $NAD^+$ related metabolites. Hydrophilic Interaction Liquid Chromatography (HILIC) LC-MS was used to measure metabolites level over 3 independent experiments. Metabolite level were normalized to total ion chromatography (TIC).

For metabolite quantification, P values were calculated by unpaired, two-sided t-test with Welch correction. *P<0.05; **P<0.01; ****P<0.0001. ns: Not significant, ND: Not Detected.

**Fig. S7**: **Validation of sir-2.1-dependent factors.**

**A-B)** Lifespan curves for N2 animals supplemented with nicotinic acid (NA – 1 mM) **(C)**, nicotinamide (NAM – 100 µM) and 1-N-methylnicotinamide (MNA 1 µM) **(D)**. Animals were grown on control or supplemented diets.

**C)** Lifespan curves for *gad-3(tm7132)* animals. Animals were grown on control diet or supplemented with 10 mM lactate

**D)** Quantification of gst-4::GFP expression under different treatment. Animals were grown on control, lactate (10 mM), NAM (100 µM) or MNA (1 µM) enriched diets. GFP level were measured in day 1 adults.

**E-F)** Quantification of gst-4::GFP expression under different doses of NAM and MNA. Animals were grown on control, lactate (10 mM) or different doses of NAM (100, 500, 1000 and 2000 µM) or MNA (1, 5, 10 and 20 µM). GFP level were measured in day 1 adults.

Statistics for lifespan curves were done using log-rank (Mantel–Cox) method. For each condition, 90 animals were analyzed (in triplicates) each experiment was repeated at least twice. P values for fluorescence measure were calculated by unpaired, two-sided t-test with Welch correction. Each fluorescence experiment was performed at least 3 times and between 25-40 animals were measured per condition. *P<0.05; **P<0.01; ****P<0.0001. ns: Not significant.

**Fig S8: Characterization of lipid derived metabolites regulated by lactate and under stress.** (*related to figure 5*)

**A-E)** Quantification of lipid-associated molecules regulated by lactate relative to control diet.

**F)** Quantification of free fatty acid in lactate-treated animals. Metabolites were measured using post-column ionization (PCI) in ESI(-) over 3 independent experiments.

**G-I)** Quantification of lipid-associated molecules under paraquat treatment relative to untreated controls. N2 animals were grown on control diet or supplemented with (100 µM) paraquat during development.

For metabolite quantification, P values were calculated by unpaired, two-sided t-test with Welch correction. *P<0.05; **P<0.01; ****P<0.0001. ns: Not significant. Metabolite measure represent one experiment, performed in triplicates. Each experiment was performed three times.

**Fig. S9: Characterization of lipid derived metabolites regulated by lactate during aging.** *(related to figure 5)*

**A-D)** Relative concentration of LPC **(A)**, LPE **(B)**, NAE **(C)** and Carnitine **(D)** in day 5 control diet, compared to day 1 control.

**E-H)** Relative concentration of LPC **(E)**, LPE **(F)**, NAE **(G)** and Carnitine **(H)** in day 5 lactate samples, compared to day 5 control.

For metabolite quantification, P values were calculated by unpaired, two-sided t-test with Welch correction. *P<0.05; **P<0.01; ****P<0.0001. ns: Not significant. Metabolite measure represent one experiment, performed in triplicates. Each experiment was performed three times.

**Fig S10 Characterization of lipid metabolism under lactate diet** *(related to figure 7)*

**A)** Images of N2 animals stained with Oil Red O (ORO). Animals were grown on control or lactate enriched diet (10 mM) and imaged as adult day 1, 3 and 5.

**B)** Images of fat-6::GFP reporter. Animals were grown on control or lactate enriched diet (10 mM) and imaged as day 1 adult. Quantification of fat-6::GFP.

**C)** Lifespan curves of N2 animals treated with *fat-7* RNAi. Animals were grown on control or lactate-enriched diet, on control RNAi.

P values for fluorescence measure were calculated by unpaired, two-sided t-test with Welch correction. Each fluorescence experiment was performed at least 3 times and between 25-40 animals were measured per condition. *P<0.05; **P<0.01; ****P<0.0001. ns: Not significant. Statistics for lifespan curves were done using log-rank (Mantel–Cox) method. For each condition, 90 animals were analyzed (in triplicates) each experiment was repeated at least twice

**Fig S11: Changes in lipid-associated metabolites are dependent on rict-1*.***

**A)** Schematic of metabolomics experiments comparing N2 and *rict-1(ft7)* animals on control or lactate-enriched diet. Samples were harvested at day 1 adult stage.

**B-D)** Quantification of lactate-regulated LPC **(B)**, Carnitine **(C)** and N-acetylethanolamine (NAE) **(D)** in *rict-1(ft7)* mutants.

For metabolite quantification, P values were calculated by unpaired, two-sided t-test with Welch correction. *P<0.05; **P<0.01; ****P<0.0001. ns: Not significant. Metabolite measure represent one experiment, performed in triplicates. Each experiment was performed three times.

**Fig S12: Analysis of NAE and phospholipids biosynthesis pathways under lactate diet.**

**A-B)** Lifespan curves for NAE-related factors. Loss of function mutant *nape-1(tm3860)* and *nape-2(tm6254)* supplemented with 10 mM L-lactate. Animals were grown on control diet or lactate-enriched media.

**C)** Lifespan curves for *faah-4(lt121)* supplemented with 10 mM L-lactate. Animals were grown on control diet or lactate-enriched media.

**D-E)** Lifespan curves for LPC/LPE-related factors. Loss of function mutant *sir-2.1(ok434)* were grown on control RNAi, on control or lactate-enriched (10 mM) media. Animals were transferred on *cept-1* **(C)** or *pcyt-1* **(D)** RNAi at L4 stages.

Statistics for paralysis curves were done using log-rank (Mantel–Cox) method. For each condition, 90 animals were analyzed (in triplicates) each experiment was repeated at least twice. ns: Not significant
