## Supplementary material for "Lactate promotes longevity through redox-driven lipid remodeling in *Caenorhabditis elegans*": Table I

Table I: Strains list used in the study

| Strain name | Genotype | Source |
|---|---|---|
| **N2** | WT | CGC |
| **JDM458** | *unc-119(ed3)* III; veyTi12[*Prpl-28*::mitoLbNOX + *unc-119*] | Joshua D. Meisel |
| **JDM459** | *unc-119(ed3)* III; veyTi13[*Prpl-28*::mitoLbNOX + *unc-119*] | Joshua D. Meisel |
| **JDM460** | *unc-119(ed3)* III; veyTi14[*Prpl-28*::cytoLbNOX + *unc-119*] | Joshua D. Meisel |
| **JDM461** | *unc-119(ed3)* III; veyTi15[*Prpl-28*::cytoLbNOX + *unc-119*] | Joshua D. Meisel |
| **CL2166** | dvIs19 [(pAF15)*gst-4p*::GFP::NLS] | CGC |
| **CB307** | *unc-47(e307)* | CGC |
| **AM44** | rmIs190 [F25B3.3p::Q67::CFP] | CGC |
| **LD1** | *skn-1(zu67)* | CGC |
| **VC1518** | *atf-7(gk715)* | CGC |
| **RB1206** | *rsks-1(ok1255)* | CGC |
| **MQ887** | *isp-1(qm150)* | CGC |
| **KX15** | *ife-2(ok3206)* | CGC |
| **CF1038** | *daf-16(mu86)* | CGC |
| **PS3551** | *hsf-1(sy441)* | CGC |
| **RB772** | *atf-6(ok551)* | CGC |
| **RB759** | *akt-1(ok525)* | CGC |
| **TJ1052** | *age-1(hx546)* | CGC |
| **JIN1375** | *hlh-30(tm1978)* | CGC |
| **RB754** | *aak-2(ok524)* | CGC |
| **KU25** | *pmk-1(km25)* | CGC |
| **VC3201** | *atfs-1(gk3094)* | CGC |
| **TM2457** | *xbp-1(tm2457)* | NBRP |
| **VC199** | *sir-2.1(ok434)* | CGC |
| **VC1061** | *anmt-1(gk457)* | CGC |
| **FCS83** | dvIs19 [(pAF15)*gst-4p*::GFP::NLS]; *sir-2.1(ok434)* | This study |
| **TM7132** | *gad-3(tm7132)* | NBRP |
| **KQ1366** | *rict-1(ft7)* | CGC |
| **XA7702** | *mdt-15(tm2182)* | CGC |
| **CE541** | *sbp-1(ep79)* | CGC |
| **VC870** | *nhr-49(gk405)* | CGC |
| **FCS100** | *sir-2.1(ok434); rict-1(ft7)* | This study |
| **FCS75** | dvIs19 [(pAF15)*gst-4p*::GFP::NLS]; *rict-1(ft7)* | This study |
| **FCS95** | nIs590 [fat-7p::fat-7::GFP + lin15(+)]; *sir-2.1(ok434)* | This study |
| **FCS97** | nIs590 [fat-7p::fat-7::GFP + lin15(+)]; *rict-1(ft7)* | This study |
| **FCS98** | dvIs19 [(pAF15)*gst-4p*::GFP::NLS]; *sir-2.1(ok434)*; *rict-1(ft7)* | This study |
| **TM5011** | *faah-1(tm5011)* | NBRP |
| **OD3609** | *faah-4(lt121)* | CGC |
| **TM3860** | *nape-1(tm3860)* | NBRP |
| **TM6254** | *nape-2(6254)* | NBRP |
